# Ensovibep, a novel trispecific DARPin candidate that protects against SARS-CoV-2 variants

**DOI:** 10.1101/2021.02.03.429164

**Authors:** Sylvia Rothenberger, Daniel L. Hurdiss, Marcel Walser, Francesca Malvezzi, Jennifer Mayor, Sarah Ryter, Hector Moreno, Nicole Liechti, Andreas Bosshart, Chloe Iss, Valérie Calabro, Andreas Cornelius, Tanja Hospodarsch, Alexandra Neculcea, Thamar Looser, Anja Schlegel, Simon Fontaine, Denis Villemagne, Maria Paladino, Yvonne Kaufmann, Doris Schaible, Iris Schlegel, Dieter Schiegg, Christof Zitt, Gabriel Sigrist, Marcel Straumann, Feyza Sacarcelik, Julia Wolter, Marco Comby, Julia M. Adler, Kathrin Eschke, Mariana Nascimento, Azza Abdelgawad, Achim D. Gruber, Judith Bushe, Olivia Kershaw, Heyrhyoung Lyoo, Chunyan Wang, Wentao Li, Ieva Drulyte, Wenjuan Du, H. Kaspar Binz, Rachel Herrup, Sabrina Lusvarghi, Sabari Nath Neerukonda, Russell Vassell, Wei Wang, Susanne Mangold, Christian Reichen, Filip Radom, Charles G. Knutson, Kamal K. Balavenkatraman, Krishnan Ramanathan, Seth Lewis, Randall Watson, Micha A. Haeuptle, Alexander Zürcher, Keith M. Dawson, Daniel Steiner, Carol D. Weiss, Patrick Amstutz, Frank J.M. van Kuppeveld, Michael T. Stumpp, Berend-Jan Bosch, Olivier Engler, Jakob Trimpert

**Affiliations:** Spiez Laboratory, Austrasse, 3700 Spiez, Switzerland; Institute of Microbiology, University Hospital Center and University of Lausanne, Rue du Bugnon 48, 1011 Lausanne, Switzerland; Molecular Partners AG, Wagistrasse 14, 8952 Zurich-Schlieren, Switzerland; Department Biomolecular Health Sciences, Division Infectious Diseases & Immunology - Virology section, Faculty of Veterinary Medicine, Utrecht University, 3584 CL, Utrecht, The Netherlands.; Cryo-Electron Microscopy, Bijvoet Center for Biomolecular Research, Department of Chemistry, Faculty of Science, Utrecht University, Padualaan 8, 3584 CH Utrecht, The Netherlands; Materials and Structural Analysis, Thermo Fisher Scientific, Eindhoven, 5651 GG, The Netherlands; Binz Biotech Consulting, Lüssirainstrasse 52, 6300 Zug, Switzerland; Laboratory of Immunoregulation, Division of Viral Products, Center for Biologics Evaluation and Research, U.S. Food and Drug Administration, Silver Spring, Maryland, USA; Freie Universität Berlin, Institut für Virologie, Robert-von Ostertag-Straße 7-13, 14163 Berlin, Germany; Freie Universität Berlin, Institut für Tierpathologie, Robert-von Ostertag-Straße 15, 14163 Berlin, Germany; Novartis Institutes for BioMedical Research, PK Sciences, Cambridge, MA, USA; Novartis Institutes for BioMedical Research, Preclinical Safety, Basel, Switzerland; Novartis Pharma AG, Basel, Switzerland

**Keywords:** SARS-CoV-2, COVID-19, coronavirus, mutations, emerging variants, antiviral therapy, ensovibep, MP0420, DARPin drug, ankyrin repeat protein, DARPin, multispecific, K417N, K417T, L452R E484K, N501Y, B.1.1.7, B.1.1.529, B.1.351, P.1, B.1.429, B.1.526, B.1.617, B.1.618, B.1.621, AY.1, alpha, beta, gamma, delta, mu, omicron, Roborovski dwarf hamster

## Abstract

SARS-CoV-2 has infected millions of people globally and continues to undergo evolution. Emerging variants can be partially resistant to vaccine induced immunity and therapeutic antibodies, emphasizing the urgent need for accessible, broad-spectrum therapeutics. Here, we report a comprehensive study of ensovibep, the first trispecific clinical DARPin candidate, that can simultaneously engage all three units of the spike protein trimer to potently inhibit ACE2 interaction, as revealed by structural analyses. The cooperative binding of the individual modules enables ensovibep to retain inhibitory potency against all frequent SARS-CoV-2 variants, including Omicron BA.1 and BA.2, as of February 2022. Moreover, viral passaging experiments show that ensovibep, when used as a single agent, can prevent development of escape mutations comparably to a cocktail of monoclonal antibodies (mAb). Finally, we demonstrate that the very high in vitro antiviral potency also translates into significant therapeutic protection and reduction of pathogenesis in Roborovski dwarf hamsters infected with either the SARS-CoV-2 wild-type or the Alpha variant. In this model, ensovibep prevents fatality and provides substantial protection equivalent to the standard of care mAb cocktail. These results support further clinical evaluation and indicate that ensovibep could be a valuable alternative to mAb cocktails and other treatments for COVID-19.

## Introduction

The extent of the COVID-19 pandemic allowed SARS-CoV-2 to quickly undergo adaptive evolution. The main mutations localize to the spike protein, a metastable prefusion trimer on the viral membrane that mediates virus entry into the host cell. The spike protein comprises multiple functional subunits: S1, which includes the N-terminal domain (NTD) and the receptor binding domain (RBD), responsible for interaction with the angiotensin-converting enzyme 2 (ACE2) host receptor^1–4^, and the S2 subunit, which is responsible for virus-host cell membrane fusion via extensive, irreversible conformational changes^5–8^. In the first months of the pandemic, a single mutation, D614G, located in the S2 domain, became prevalent. This mutation impairs premature conformational change of the spike protein, thus increasing the number of infectious viral particles and therefore overall viral infectivity^9^. By November 2021, more viral lineages have been identified and designated as Variants of Interest (VOIs) or Variants of Concern (VOCs) based on their associated increased risk to public health. These were first isolated in the UK (Alpha, B.1.1.7 lineage), South Africa (Beta, B.1.351), Brazil (Gamma, P.1), South California (Epsilon, B.1.429), Nigeria (Eta, B.1.525), New York, (Iota, B.1.526), Peru (Lambda, C.37), Japan (R.1), India (Kappa, B.1.617.1 and Delta, B.1.617.2), Uganda (A.23.1), and, more recently, in Colombia (Mu, B.1.621), the UK (Delta Plus, AY.1), as well as Africa (Omicron, B.1.1.529) ^10–22^.

Many of these variants harbor mutations in the RBD domain of the spike protein, mainly in the ACE2 binding site (K417T/N, N439K, L452R, E484K/Q, N501Y). Since this region is also highly immunogenic, these mutations have been linked to a dual effect: either increasing the affinity to the human ACE2 receptor (N439K, N501Y) and therefore transmissibility, and/or facilitating immune escape of the virus (K417T/N, L452R, E484K/Q)^10,11,15–17,23–25^. In particular, the E484K substitution has been shown to play a key role in attenuating the potency gain and resistance to the majority of antibodies, according to a study analyzing clinical-stage therapeutic antibodies^12^.

Fighting the COVID-19 pandemic requires a coordinated global effort to maximize the benefits of vaccinations and therapeutics^1^. The presence of an unvaccinated portion of the population and the evolution of escape mutants highlights the medical need for globally accessible therapeutics^26^. Neutralizing mAbs are a critically important therapeutic approach against COVID-19. To circumvent their loss of potency due to viral mutational escape, antibody cocktails were generated to provide increased protection against variants^27–29^.

We have applied the DARPin platform^30^, which allows fast generation and cost-effective production of biological therapeutics, to generate ensovibep, an anti-SARS-CoV-2 multispecific DARPin antiviral clinical candidate^31,32^. DARPins are an emerging class of novel therapeutics that are actively being developed in ophthalmology and oncology^33,34^. They are structurally fully differentiated from antibodies and consist of a single chain of linked DARPin binding domains. In the case of ensovibep, the molecule comprises two human serum albumin binding DARPin domains for systemic half-life extension^35^ (H1 and H2) and three spike protein RBD-binding DARPin domains at the C-terminus (R1, R2 and R3). The relatively small size of ensovibep (85 kDa), in conjunction with high thermal stability^31^, high production yields^31^ and demonstrated high protection against viral escape mutations and variants makes this molecule an attractive alternative to other treatments.

Using structural analysis, we provide an explanation for ensovibep-mediated neutralization of the SARS-CoV-2 spike protein. The three distinct DARPin domains can simultaneously target the receptor binding ridge on each RBD of the spike trimer, locking the spike in an open-conformation and occluding the ACE2 binding site. Thanks to the cooperative binding of this novel trispecific design, ensovibep confers very high protection against a panel of relevant spike mutants as well as all frequent SARS-CoV-2 variants identified around the globe to date. We show in a viral passaging experiment that the protection provided by ensovibep against development of viral escape mutants is equivalent to that of a well characterized and clinically evaluated monoclonal antibody cocktail^27,36,37^.

Following our *in vitro* characterization, we demonstrate high *in vivo* efficacy in a therapeutic Roborovski dwarf hamster model of COVID-19. Here ensovibep protects against severe disease induced by either wild-type or the Alpha variant SARS-CoV-2. The Roborovski dwarf hamster is highly susceptible to SARS-CoV-2 infection and develops strong lung pathology, with most animals reaching a defined humane endpoint within two to five days after infection^38^. In the presented study, ensovibep protects the animals to an extent equivalent to a standard-of-care mAb cocktail. For both therapeutic agents, a significant reduction of fulminant disease, as well as significantly reduced viral loads and attenuated lung pathology was observed.

In brief, the trispecific design of ensovibep provides great protection against all currently known SARS-CoV-2 variants with the potential to protect against emerging variants in the future. Our findings strongly support the progressing clinical development of ensovibep as a potential therapeutic for COVID-19.

## Results

### Structural basis for ensovibep-mediated neutralization of the SARS-CoV-2 spike

Ensovibep comprises of five covalently linked DARPin domains. Three of them (R1, R2 and R3) bind the RBD of SARS-CoV-2 with picomolar affinity (Supplementary Figure 1) and two of them (H1-H2) bind to human serum albumin (HSA), extending the systemic half-life (Figure 1A). To understand how ensovibep binds to the SARS-CoV-2 spike (S), we selected one of the three RBD-targeting DARPin domains of ensovibep for cryo-EM analysis in complex with the trimeric S-ectodomain. The RBD-binding domains are from the same sequence family and are thus expected to target a common epitope (Figure 1B). Upon incubation of the trimeric spike protein with the monovalent DARPin R2 for 15 seconds prior to vitrification, 3D classification revealed that 65% of the S-ectodomains were in the closed conformation, 20% had two RBDs in the open conformation and 15% had all three RBDs in the open conformation (Supplementary Figure 2A, B). For the open RBD classes, additional density, consistent with the size of the monovalent DARPin molecule, was present on the RBD receptor binding ridge (RBR). When the incubation time was increased to 60 seconds, 66% of S-ectodomains had three monovalent DARPin molecule-bound RBDs in the open conformation (Supplementary Figure 2C). Interestingly, 18% of the S-ectodomains had two DARPin-bound RBDs in the open conformation and one trapped in a partially closed conformation (Supplementary Figure 2C and 3A-B). These results demonstrate that monovalent DARPin domain binding prevents closure of the RBD through a previously described ratcheting mechanism^39^. 3D refinement of the fully open class, from the 60 second incubated sample, produced a 4.2 Å global resolution map (Figure 1C and Supplementary Figure 2D-F). Following focused refinement of the RBD region, the quality of the map was sufficient to unambiguously assign the pose of the monovalent DARPin domain, which binds perpendicular to the RBD receptor binding motif (RBM), with its N-terminus orientated toward the spike three-fold symmetry axis (Figure 1C). The concave DARPin binding surface covers the RBD and would prevent ACE2 binding through steric hindrance (Figure 1D). Guided by the cryo-EM data, molecular docking experiments were performed between the RBD of SARS-CoV-2 and DARPin R2.

**Figure 1:**
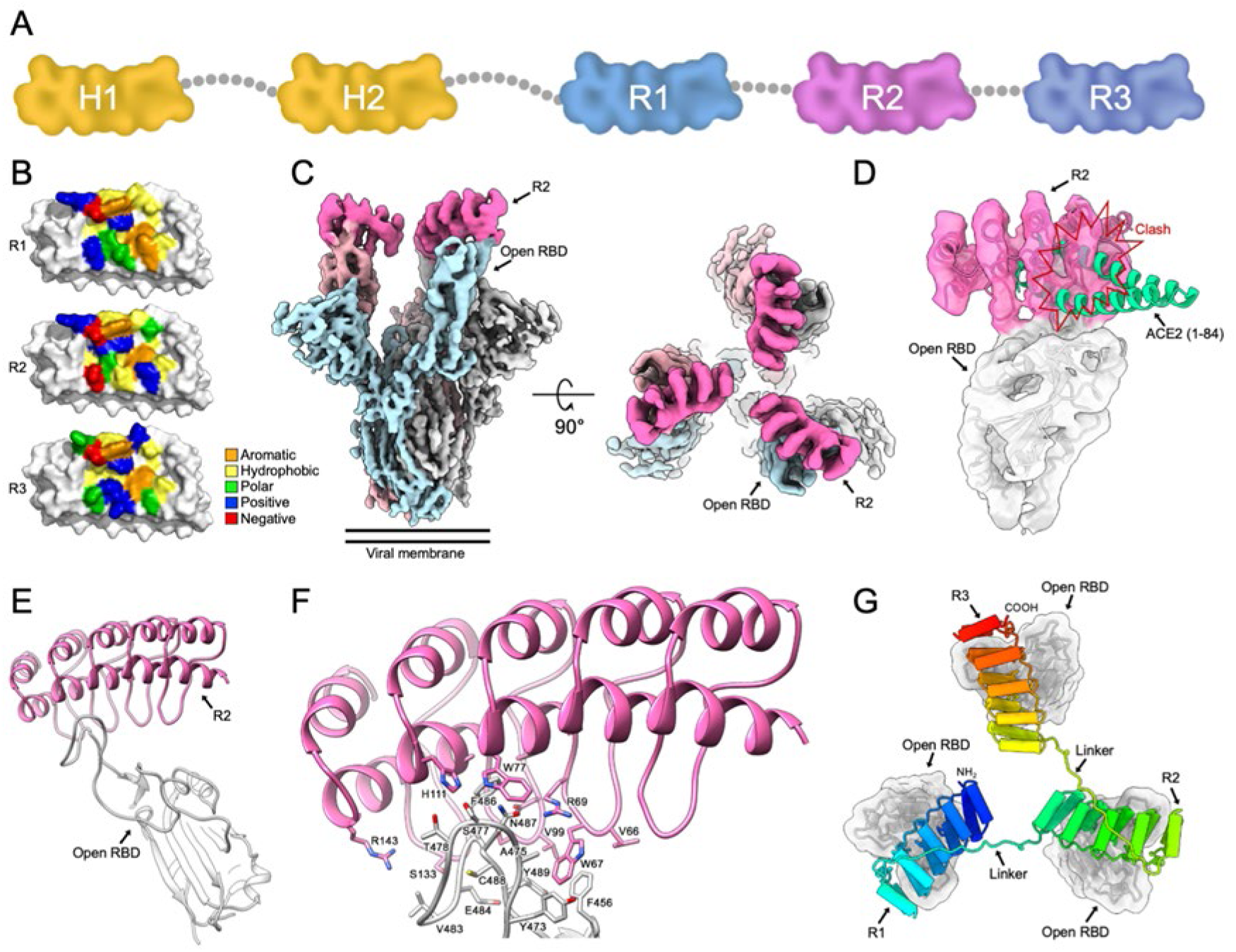
Structural modelling of ensovibep. A) Schematic overview of the ensovibep construct. Protein linkers are depicted as gray dashed lines and the half-life extending human serum albumin binding monovalent DARPins (H1 and H2) are colored yellow. B) Surface representations of the three monovalent DARPin molecules binding to the RBD, with the amino acid residues in the paratope colored according to their biophysical characteristics as indicated. C) Cryo-EM density for the SARS-CoV-2 spike ectodomain in complex with the RBD-targeting monovalent DARPin R2, shown as two orthogonal views. The DARPin density is colored magenta and the three spike protomers are colored light blue, grey and pale pink. D) Zoomed in view of an RBD-bound to DARPin R2 with the cryo-EM density shown semi-transparent. The atomic coordinates for the fitted open RBD (PDB ID: 6XCN) and the DARPin model are overlaid. The atomic coordinates for residues 1-84 of the RBD-bound ACE2 (PDB ID: 6M0J), colored green, is superimposed. E) Pseudo-atomic model of the monovalent DARPin R2 in complex with the RBD, colored pink and grey, respectively. F) Zoomed in view of the interface between monovalent DARPin R2 and RBD. G) Proposed model of the three covalently linked RBD-targeting monovalent DARPin molecules of ensovibep bound to the trimeric spike protein RBD domains. The three DARPin domains are shown in a rainbow color scheme from the N terminus (blue) to the C terminus (red).

The top scoring model indicated that the interface area is ∼700 Å^2^ and that key epitope residues are F456, Y473, F486, N487 and Y489, which form an interface of hydrophobic interactions and hydrogen-bonds with the DARPin domain (Figure 1E-F). Because the three DARPin domains share a similar paratope composition and architecture, we were able to conceptually model the entire ensovibep molecule bound to the fully open S-ectodomain (Figure 1G). This demonstrated that the linkers would permit simultaneous binding of all three DARPin modules, allowing very high avidity of ensovibep (Supplementary Figure 1), and that the half-life extension modules have sufficient space to bind HSA (not shown). Taken together, these data suggest that ensovibep inhibits SARS-CoV-2 by blocking ACE2 binding and promoting the premature conversion of spike to the post-fusion state. This mechanism of inhibition through receptor functional mimicry was observed for a number of SARS-CoV-2 neutralizing antibodies^39,40^.

### Ensovibep is highly potent against globally identified SARS-CoV-2 variants as well as the most frequent spike protein point mutations

In order to assess the neutralizing potencies of ensovibep against the initial SARS-CoV-2 (Wuhan) and emerging variants, we used vesicular stomatitis virus (VSV)-based as well as lentivirus-based pseudoviruses carrying the SARS-CoV-2 wild-type or mutant spike protein at their surface. In addition, we tested the authentic SARS-CoV-2 variants for the Wuhan reference and for lineages B.1.1.7, B.1.351 and P.1. Ensovibep is able to neutralize the reference wild type strain with an IC_50_ of ∼1 ng/mL, when either the authentic SARS-CoV-2 or the pseudovirus is used (Figure 2A). Remarkably, the high neutralization efficacy is retained in all the frequent variants circulating to date, which display a diverse set of mutations over the entire length of the spike protein (Figure 2A and 2B; Supplementary Table 2; Supplementary Figure 4). In particular, ensovibep can neutralize the variants of concern (VOC) and variants of interest (VOI) of the lineage B.1.1.7/Alpha (69-70 del, del145, E484K, N501Y, A570D, D614G, P681H, T716I, S982A, D1118H and with the addition of E484K or S494P), lineage B.1.351/Beta (L18F, D80A, D215G, Del242-244, R246I, K417N, E484K, N501Y, D614G, A701V), lineage P.1/Gamma (L18F, T20N, P26S, D138Y, R190S, K417T, E484K, N501Y, D614G, H655Y, T1027I, V1176F), B.1.617.2/Delta (T19R, G142D, del156-157, R158G, L452R, T478K, D614G, P681R, D950N), AY.2/Delta Plus (T19R, G142D, del156-157, R158G, K417N, L452R, T478K, D614G, P681R, D950N), AY.4.2/Delta Plus (T19R, T95I, G142D, Y145H, E156G, F157-, R158-, A222V, L452R, T478K, D614G, P681R, D950N), Lambda (C.37; G75V, T76I, del246-252, D253N, L452Q, F490S, D614G, T859N), Mu (B.1.621; T95I, Y144S, Y145N, R346K, E484K, N501Y, D614G, P681H, D950N), Omicron (B.1.1.529, BA.1; A67V, Δ69-70, T95I, G142D, Δ143-145, Δ211, L212I, ins214EPE, G339D, S371L, S373P, S375F, K417N, N440K, G446S, S477N, T478K, E484A, Q493K/R, G496S, Q498R, N501Y, Y505H, T547K, D614G, H655Y, N679K, P681H, N764K, D796Y, N856K, Q954H, N969K, L981F) and Omicron (B.1.1.529, BA.2; T19I, L24-, P25-, P26-, A27S, G142D, V213G, G339D, S371F, S373P, S375F, T376A, D405N, R408S, K417N, N440K, S477N, T478K, E484A, Q493R, Q498R, N501Y, Y505H, D614G, H655Y, N679K, P681H, N764K, D796Y, Q954H, N969K). The neutralization potencies of ensovibep remain within 10-fold difference from the reference virus (Wuhan or D614G variant) with IC_50_ values in the low single-digit ng/mL range, even against those variants that have been shown to be, to a large extent, refractory to vaccine- or infection-related antibody neutralization, such as Beta, Gamma, Delta, Delta Plus, and the newly evolved Omicron variants BA.1 and BA.2.^25,41–43^ When testing the neutralizing potency in a VSV-based pseudotype assay, containing more than 30 substitutions of the Omicron spike protein, ensovibep maintained neutralization at low single digit ng/mL IC_50_ values without loss in potency, when compared to the wild type. In contrast, many of the tested clinically relevant monoclonal antibodies and antibody cocktails demonstrated a major loss in neutralization (Figure 2D).

**Figure 2:**
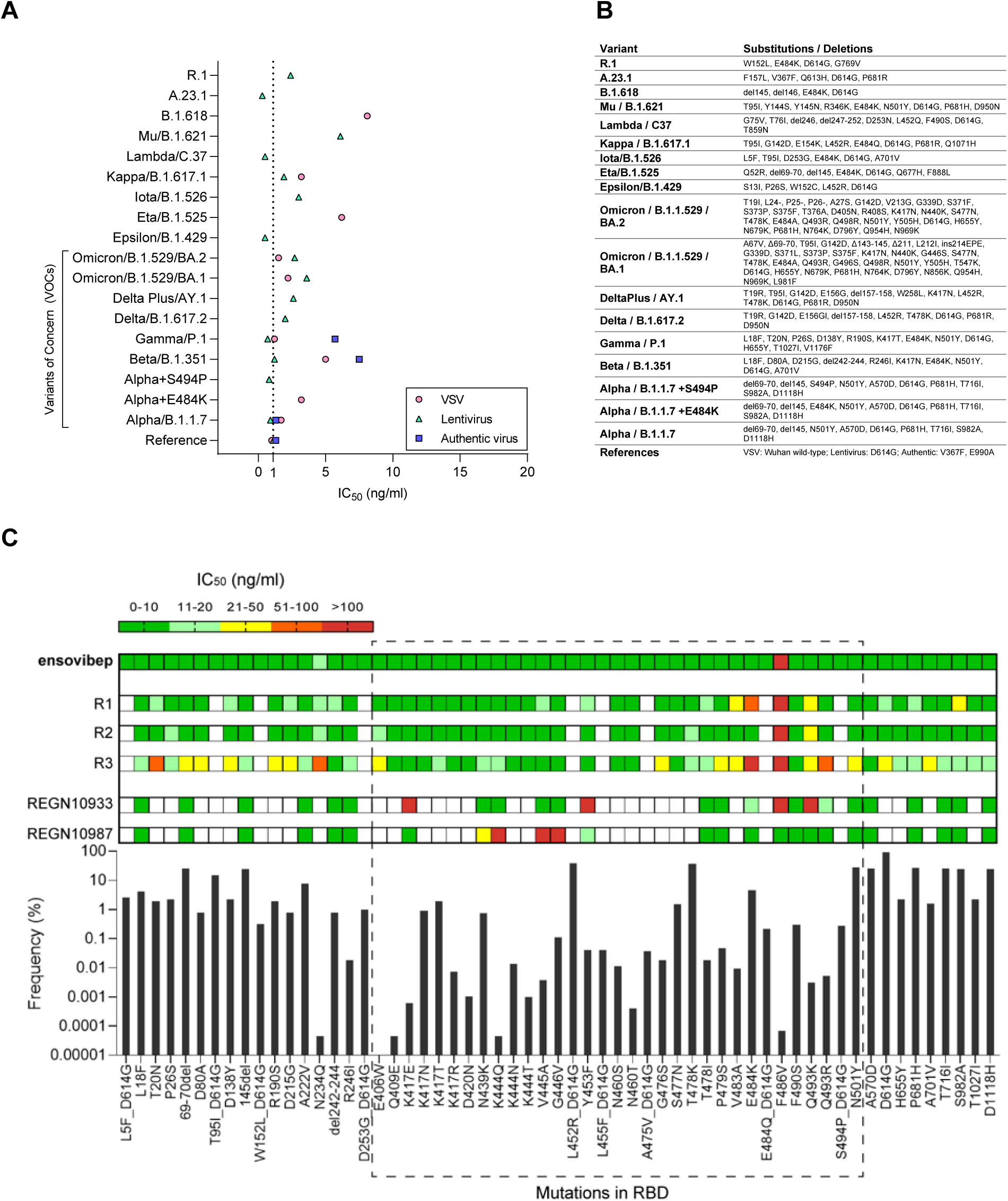
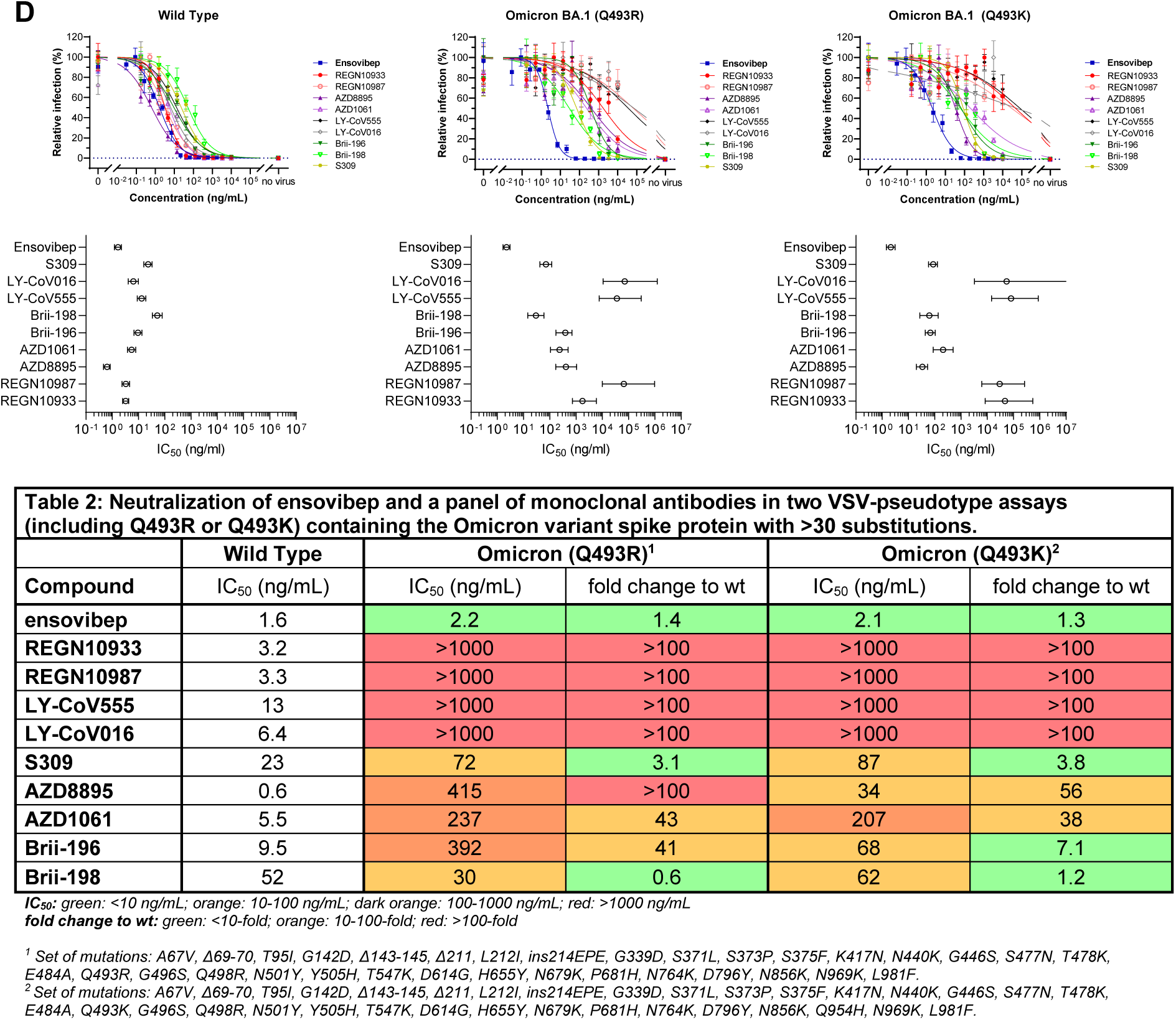
A) Graph reporting IC_50_ values (ng/mL) for ensovibep measured in neutralization assays performed with lentivirus-, VSV-based pseudoviruses or authentic viruses for the variants indicated. Reference variant is the Wuhan strain for VSV-based pseudovirus, a D614G variant for the lentivirus-based pseudovirus or a patient isolate from the early pandemic for the authentic virus. B) Schematic representation of the residues modified in the SARS-CoV-2 spike protein for the different variants tested compared to the Wuhan strain. C) Graph with global frequencies of point mutations in the spike protein of SARS-CoV-2 according to the GISAID database (as of October 2021) including a heat map table with IC_50_ values for ensovibep, R1, R2, R3, REGN10933, REGN10987 for all point mutations tested (VSV/Lentivirus-based pseudovirus assays). Dashed box: mutations in RBD. D) Titration curves (mean ±SEM) and IC_50_ values (mean ±CI at 95%) for VSV-pseudotype neutralization assays with wild-type and two different Omicron BA.1 variant spike proteins containing either an arginine or a lysine in position Q493. Ensovibep was tested together with a panel of clinically validated monoclonal antibodies. The table provides the numeric IC_50_ values as well as the fold change towards the wild-type values.

Using the VSV- and lentivirus-based pseudovirus neutralization assays, we also evaluated the influence of single mutations on the neutralization potency of ensovibep, of the monovalent DARPin molecules and of the mAbs REGN10933 and REGN10987, as a reference within the same experiment. The panel included mutations present on variants of interest/concern, appearing frequently, or located within the binding epitope of ensovibep. Most notably, ensovibep protected well against all point mutations tested, in contrast to the single monoclonal antibodies, with the only exception of substitution F486V, which affects all three monovalent DARPin RBD binders incorporated in ensovibep (Figure 2C). A major impact of this mutation is not surprising, as our structural analysis and modelling identifies F486 as a core interacting residue for the three related but distinct RBD binders^31^ (Figure 1B,F). Consequently, the mutation F486V destabilizes the binding of the entire tri-specific ensovibep molecule to the spike protein. However, F486 is also a critical residue for the interaction between the RBD of SARS-CoV-2 and human ACE2 and its mutation leads to a ∼8.5-fold reduction of the binding affinity as well as a ∼17-fold reduction of the ability of ACE2 to reduce the infection of a VSV-based pseudovirus carrying the F486L mutation (Supplementary Figure 6). The functional importance of F486 is reflected by a low frequency of naturally occurring substitutions at this site (Figure 2C; Supplementary Table 3; Supplementary Figure 4) where the selective pressure on the virus favors a phenylalanine, thus maintaining the key anchoring element for ensovibep binding. A reduction of the potency of ensovibep from one-digit to double-digit ng/mL IC_50_ was also observed for mutation N234Q. This residue is located outside of the RBD binding region of ensovibep. This minor effect of substitution N234Q could be related to the loss of the conserved glycosylation site at this position, favoring the kinetics of the down conformation of the RBD domain and thus reducing binding of ensovibep as well as ACE2 to the RBD, which only bind the RBD up confirmation.^44^

It is interesting to note that ensovibep retains potency against spike proteins carrying mutations at locations where the single DARPin domains partially lose activity, such as E484K and Q493K/R. We hypothesize that the cooperative binding in combination with the complementarity of the three independent RBD-binding DARPin modules provides resistance to mutation escape. Taken together, our analysis demonstrates that the trispecific design of ensovibep enables very high potencies against spike proteins carrying the most frequently observed mutations as well as mutations known to impact the binding of neutralizing antibodies.

#### Passaging of SARS-CoV-2 under therapeutic pressure of DARPin antivirals and monoclonal antibodies

Previous studies have shown that SARS-CoV-2 escape mutants may arise under selective pressure of a therapy^29,45^. Using a viral passaging model, we compared the risk of mutational escape from therapeutic pressure of ensovibep compared to that of its monovalent R2 module, the mAbs REGN10933 and REGN10987, singly and as a 1:1 mixture, as well as the mAb S309.

In order to generate a stringent therapeutic pressure, a relatively high viral load of 1.5 x 10^6^ pfu of an authentic French SARS-CoV-2 WT isolate (with the following differences to the Wuhan wild-type spike protein: V367F; E990A) was serially passaged in the presence of increasing concentrations of DARPin molecules and antibodies (Figure 3A, 3B). Resistant escape variants were further selected by passaging the supernatant of cultures showing significant virus-induced cytopathic effect (CPE) under the selection pressure of the highest therapeutic concentration onto fresh cells while maintaining the selective pressure of increasing concentrations of therapeutic antivirals (Supplementary Figure 5). After the first incubation cycle of four days (passage 1), ensovibep, DARPin R2, REGN10933 and the antibody mixture conferred protection at the same concentration of 0.4 µg/mL. S309 was less efficient, requiring a higher concentration (10 µg/mL) for protection and REGN10987 was not protective up to the highest tested concentration of 50 µg/mL. Under continuous selective pressure through passage 2 to 4, DARPin R2 and the individual mAbs S309 and REGN10933 lost the capacity to protect cells, which manifested in complete CPE up to 50 µg/mL. In contrast, ensovibep and the cocktail of two mAbs remained effective and protected cells from CPE throughout the four passages (Figure 3A).

**Figure 3:**
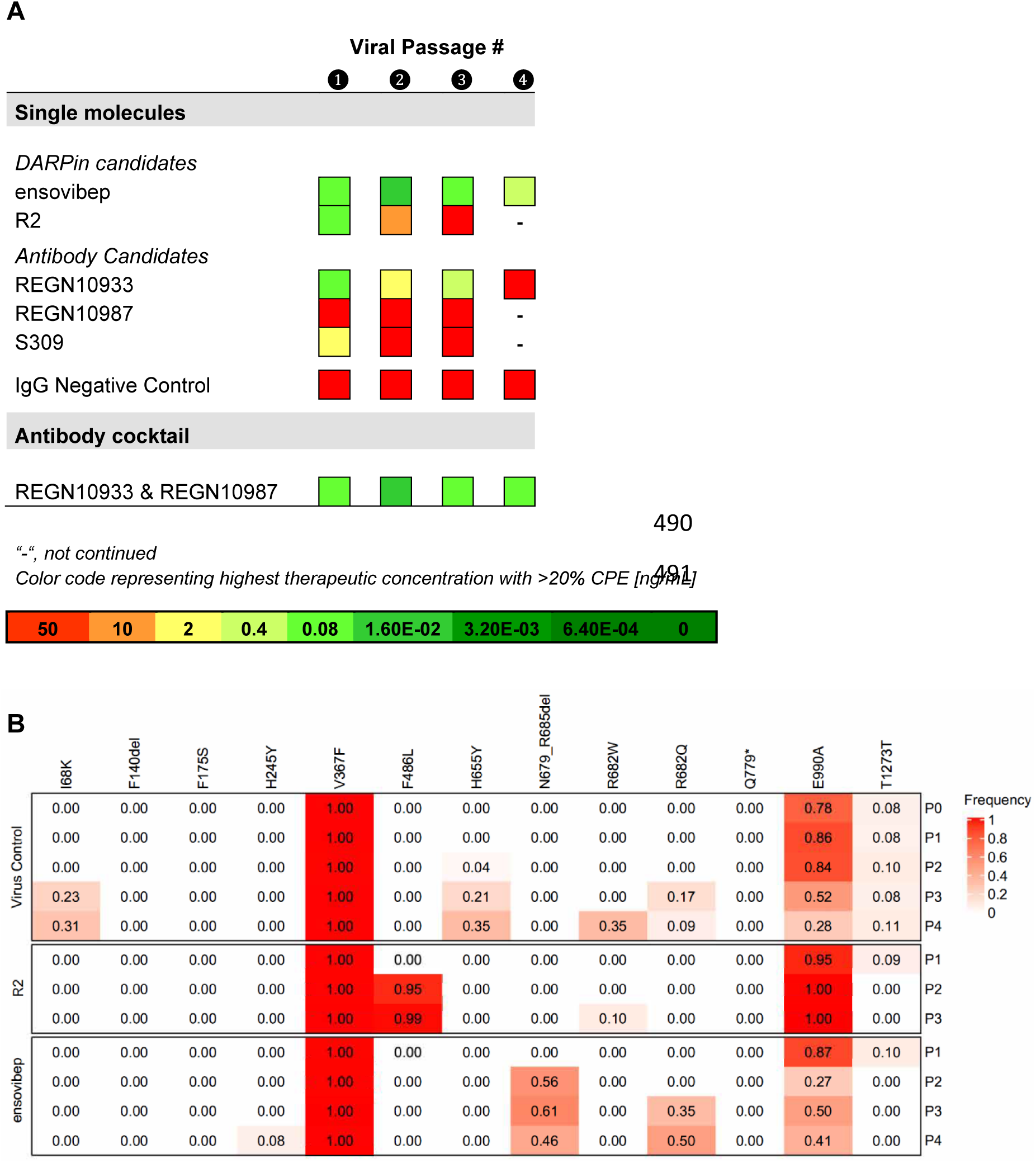
Protection against SARS-CoV-2 escape mutations generated over four viral passages. A) Tabular representation of the cytopathic effects induced by SARS-CoV-2 cultured in the presence of increasing concentrations of mono-valent DARPin binder R2, multi-specific DARPin antiviral ensovibep and the antibody antivirals REGN10933, REGN10987 and S309 or a cocktail of REGN10933 and REGN10987 through passage 1 to 4. Color code represents the highest concentration showing ≥20% CPE, for which the culture supernatants was passaged to the next round and deep sequenced for the identification of potential escape mutations. B) Identification of escape mutations in viral passages using deep sequencing. SARS-CoV-2 virus was serially passaged with the mono-valent DARPin binder R2 and ensovibep. To identify putative escape mutations in the spike protein, RNA was extracted and sequenced from supernatant of wells with the greatest selective pressure showing a significant cytopathic effect. All variants in the spike protein relative to the reference genome (NC_045512.2) are shown. Passage 0 of the virus control corresponds to the inoculum used for all experiments. The color of the fields is proportional to the fraction of the reads containing the respective variant (red= 1.0 white=0.0).

To identify putative escape mutations in the spike protein upon therapeutic pressure of the DARPins, RNA was extracted and deep-sequenced from the supernatant of wells with the greatest selective pressure showing a significant cytopathic effect in each passage (Figure 3B). Mutations were found near the spike protein cleavage site (H655Y, N679_R685del, R682W, R682Q), which are likely related to adaptations to the experimental cell system and thus would not account for escape mutations due to the therapeutic pressure of the DARPin^36,37^, as well as a potential escape mutation, F486L, which was found for the monovalent DARPin R2 but not for ensovibep, up to passaging round four. Still, supporting this finding, mutations in F486 were shown to influence also the potency of ensovibep, when analyzed separately.

#### In vivo antiviral efficacy of ensovibep in a COVID-19 SARS-CoV-2 Roborovski dwarf hamster model

To test the *in vivo* efficacy of ensovibep in treating SARS-CoV-2 infection, we employed the Roborovski dwarf hamster, a species susceptible to severe COVID-19 like illness^46^. Unlike the more commonly used Syrian golden hamster^47^, this species is prone to develop a lethal course of disease, notably without the extrapulmonary disease manifestations observed in highly susceptible transgenic mice^48^. We used this particular animal model to judge the *in vivo* efficacy of ensovibep and to compare it to the REGN10933 & REGN10987 antibody mixture. Moreover, evaluation of the virological and histopathological outcome of infection enabled comparison across a variety of important parameters of infection.

We first aimed to determine *in vivo* protection conferred by ensovibep against a SARS-CoV-2 wild type reference strain (BetaCoV/Germany/BavPat1/2020). In an initial series of experiments, we determined both dose and time dependency of treatment efficacy based on clinical and virological parameters. In absence of venous access in dwarf hamsters, we choose intraperitoneal (i.p.) treatment for delivery of ensovibep. It is important to note, that the course of disease in Roborovski dwarf hamsters is rapid, with first animals developing severe disease and reaching termination criteria within 48 hours of infection. For this reason, we considered 24 hours post-infection (p.i.) the latest possible intervention time point. Both dose and time of ensovibep administration (relative to time of infection) were found to positively affect the outcome of infection. Specifically, the use of ensovibep resulted in markedly reduced virus loads in the respiratory tract of treated animals (Supplementary Figure 7).

From these initial results, we determined 10 mg/kg to be the optimal dose for ensovibep treatment and in further studies compared this dose with the same dose of the REGN10933 & REGN10987 cocktail using the SARS-CoV-2 alpha (B.1.1.7) variant of a more recent isolate (BetaCoV/Germany/ChVir21652/2020) for infection of animals. We chose two treatment time points, the first at the time of infection to mimic clinical post exposure prophylaxis and the second at 24 h p.i. to mimic treatment at the onset of clinical symptoms (Figure 4A). For the post exposure prophylaxis dosed directly after infection (0 h p.i.), we confirmed full protection for both treatments with notable reduction of viral loads, particularly in the lungs of treated animals compared to placebo treated controls at all time points (Figure 5A). There were no obvious differences between the two agents, however, based on virological parameters, a slight trend towards lower viral load in the antibody cocktail group was observed at 5 days p.i. (Figure 5A).

**Figure 4:**
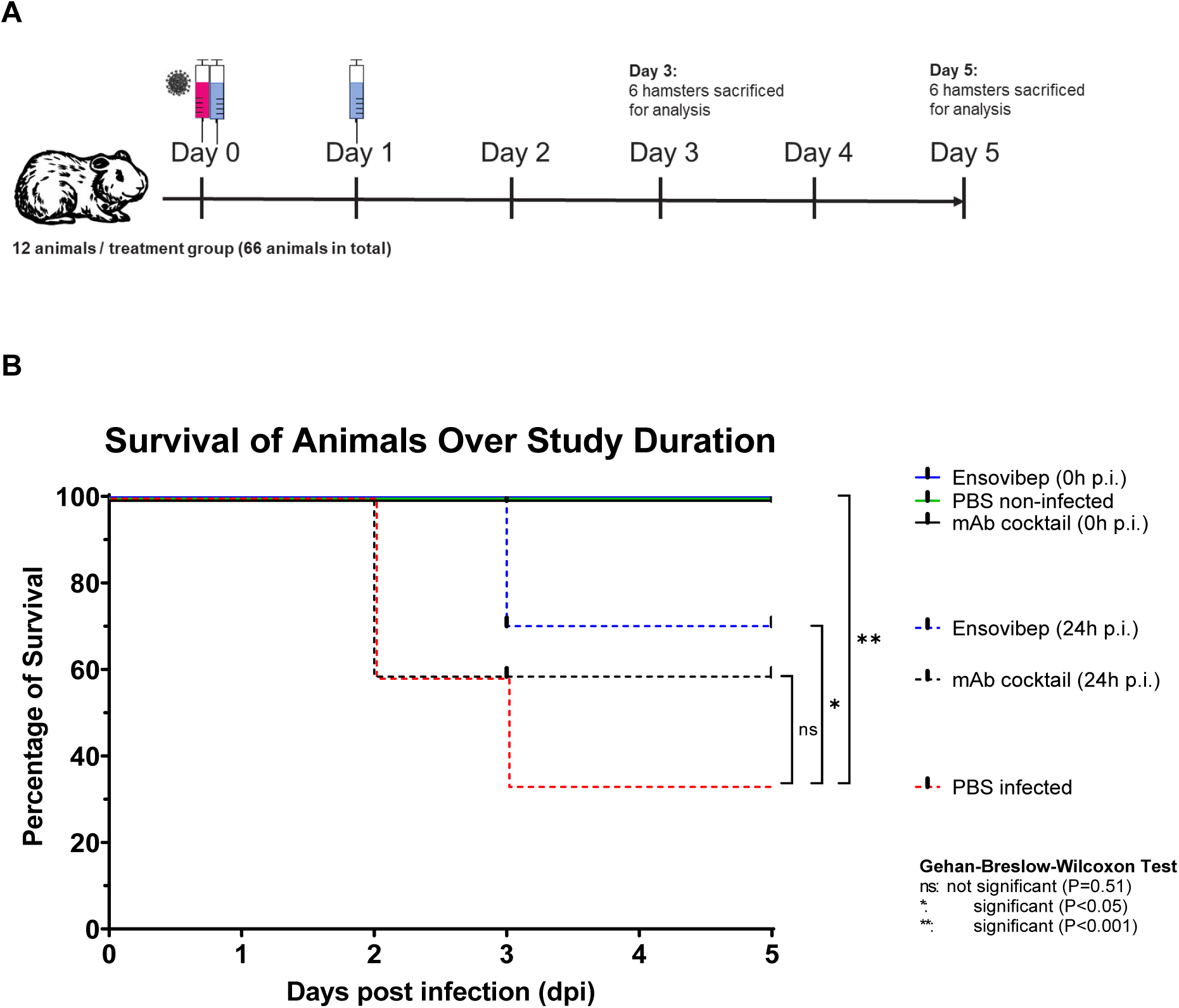
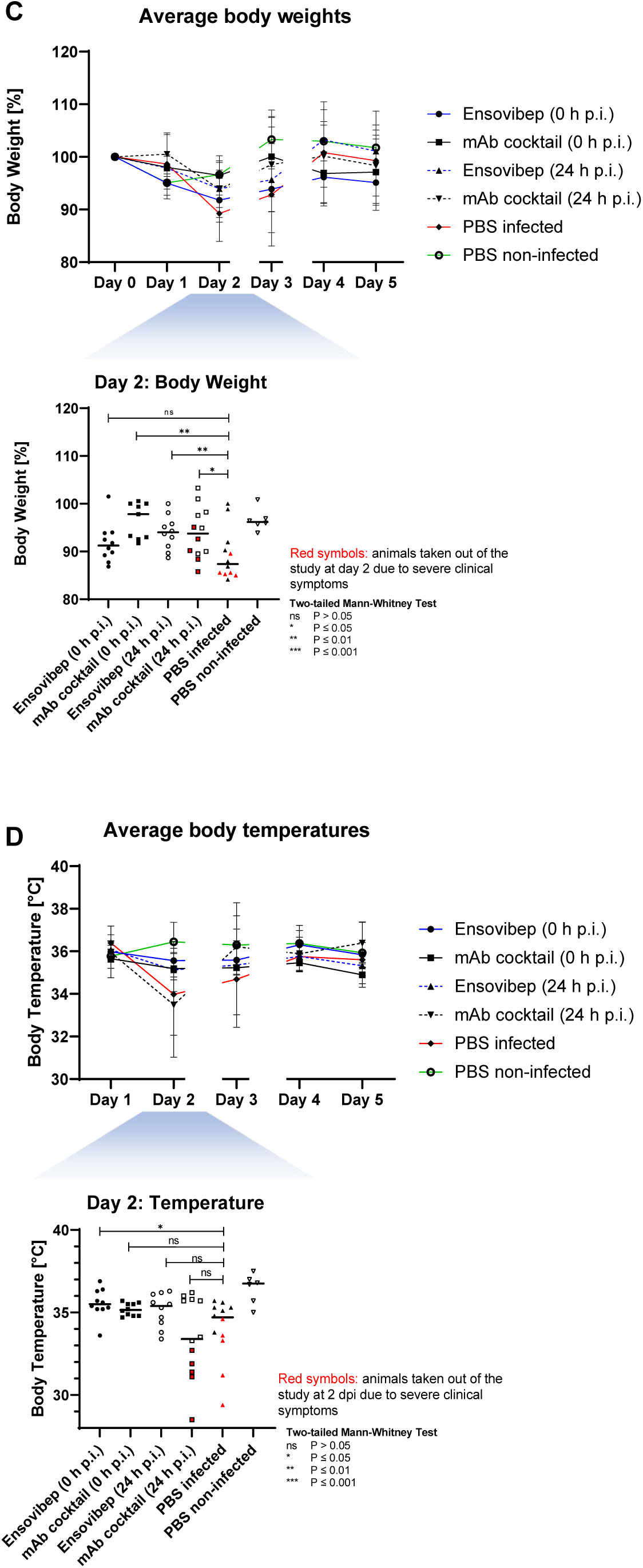
A) Design of the Roborovski dwarf hamster study. Animals were infected on day 0 with 10^5^ pfu of SARS-CoV-2 alpha (B.1.1.7) variant. Treatment was administered either directly following infection (0 h p.i.) or one day post infection (24 h p.i). For each treatment group, twelve animals were injected i.p. with either 10 mg/kg of ensovibep, 10 mg/kg monoclonal antibody cocktail (5 mg/kg REGN10933 & 5 mg/kg REGN10987), or PBS (placebo). Additionally, a group of six non-infected and non-treated control animals were included as comparators for the infected and treated groups. Daily measurement of body weight and temperatures as well observation of vital symptoms was undertaken. Animals were sacrificed on day 3 or 5 p.i. or immediately once an individual reached a defined humane endpoint. B) Survival of animals for 5 days p.i.. Animals that had to be euthanized according to defined humane endpoints were considered as non-survived. C) Body weight and D) body temperatures throughout the study duration. Data points show mean +/- SD of the following number of animals analyzed per treatment group at (0/1/2/3/4/5) days p.i.: Esovibep 0h: n=10/10/10/10/5/5; mAb cocktail 0h: n=9/9/9/9/5/5; Esovibep 24h: n=10/10/10/10/5/5; mAb cocktail 24h: n=12/12/12/7/6/6; Placebo, infected: n=12/12/12/7/4/4; Placebo, non-infected: n=6/6/6/6/6/6. The rational for excluding animals is the identification of animals with low drug exposure, likely due to a failure of i.p. injections. These animals were excluded from all analyses. Lines connecting dots are interrupted for any change in animal numbers between consecutive days. Since a considerable number of animals in the mAb cocktail and placebo groups reached defined humane endpoints by day 2 p.i., This day is zoomed-in and values are presented the median and for each individual animal with red symbols marking animals that had to be euthanized at day 2.

**Figure 5:**
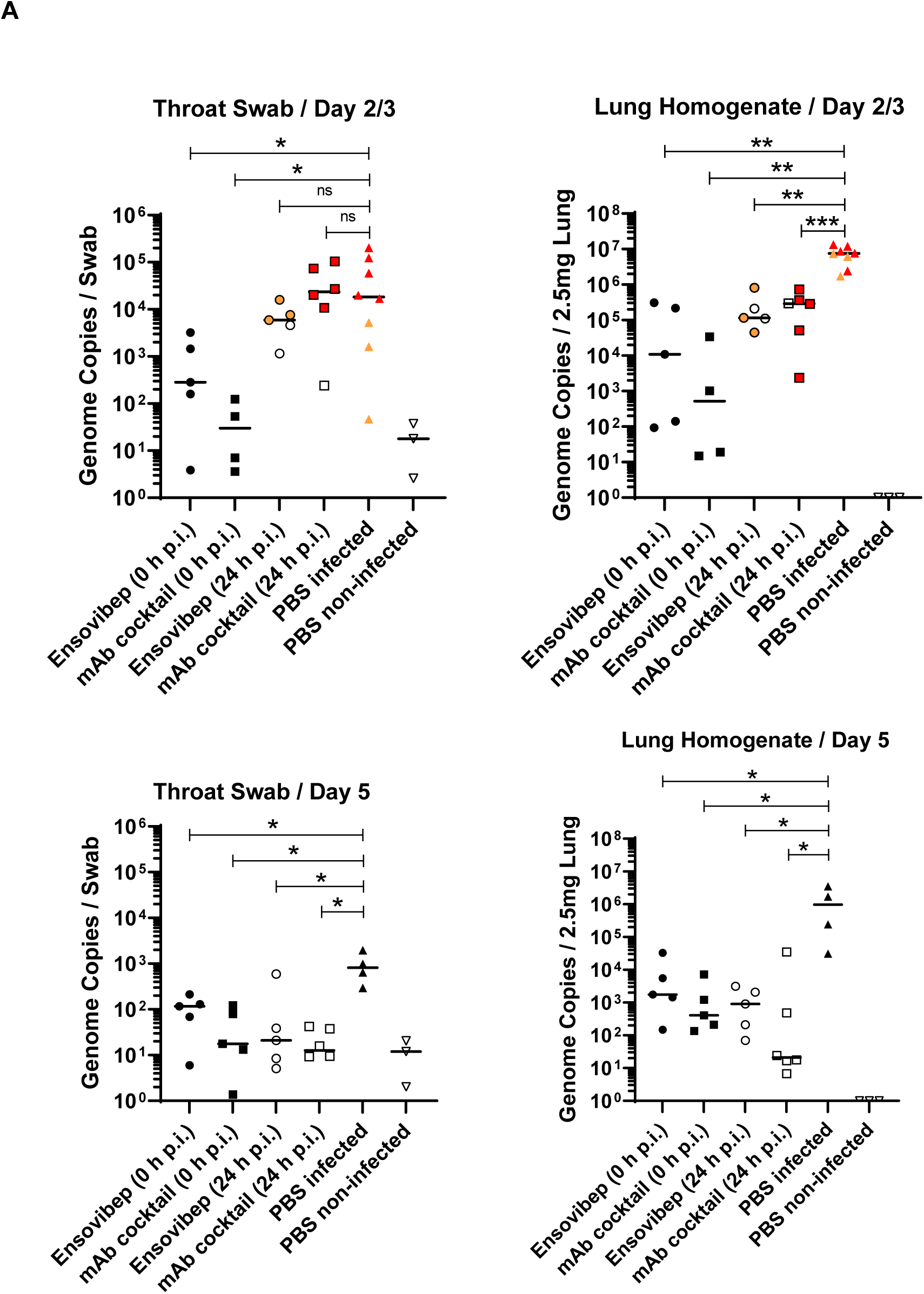
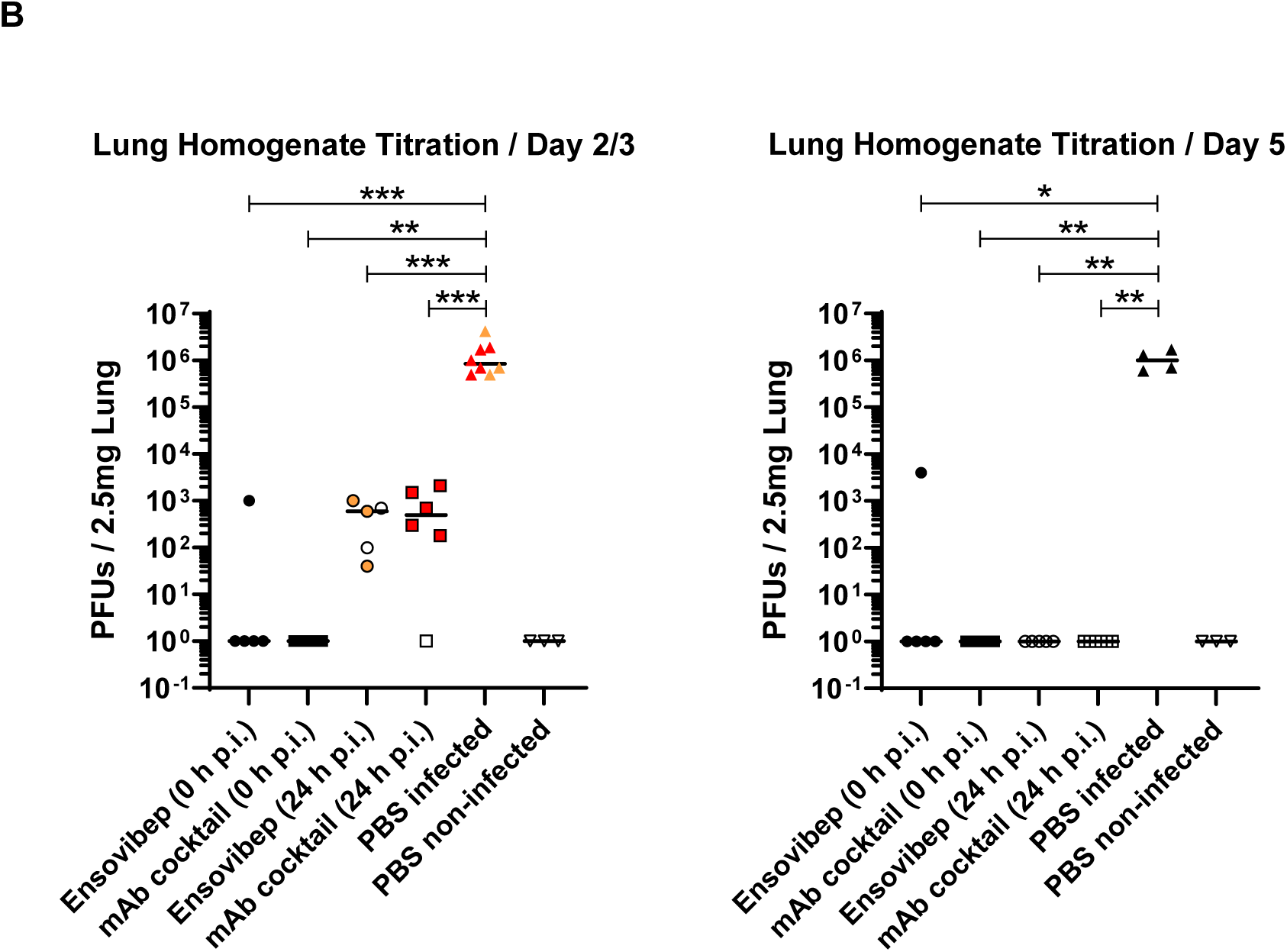
A) qPCR analysis of virus gRNA copy numbers in oropharyngeal swabs and lung homogenates at day 2/3 or day 5 p.i. B) Titration of replication competent virus from lung homogenates as plaque assay on Vero E6 cells at day 2/3 or day 5 post infection. Red symbols: animals taken out of the study at day 2 due to severe clinical symptoms. Orange symbols: animals taken out of the study at day 3 due to severe clinical symptoms. Data is represented by the median and values for individual animals. Statistics: two-tailed Mann-Whitney Test: ns P > 0.05; * P ≤ 0.05; ** P ≤ 0.01; *** P ≤ 0.001.

In contrast to the similarities in the post exposure prophylaxis setting we observed differences between the groups treated 24 hours p.i. (Figure 4B, C). In this scenario, animals treated with ensovibep presented with improved condition at 2 days p.i. with 0/12 of the animals reaching a defined humane endpoint, while 5/12 animals were euthanized in the mAb cocktail group and 5/12 in the placebo group (Figure 5B) due to reached humane endpoints. Nevertheless, 3/10 hamsters in the ensovipeb group and an additional three hamsters in the placebo group reached defined endpoints at day 3 p.i., while no further animals in the mAb cocktail group developed severe illness after day 3 p.i. (Figure 5B). Following 24 h p.i. treatment, no significant differences in average body weights or temperatures were observed in any of the treatment groups (Figure 5C, Supplementary Figure 8). This is likely a result of the early termination of severely sick animals, while the healthier animals remained in the study. However, examination of these parameters on day 2 p.i. revealed significant trends towards reduced body weight loss in both treatment groups compared to the placebo and a similar trend towards higher body temperatures in the ensovibep group compared to the other groups (Figure 5C). As body temperature decrease is a very sensitive parameter of disease in this species^46^, this in particular is reflective of the improved condition in the ensovibep treated group at 24h p.i., when compared to the antibody cocktail treated or the placebo treated animals. Virological readouts were not significantly different between groups treated with ensovibep and the mAb cocktail at 24 hours post-infection. Both treatments resulted in drastic reductions of viral load compared to the placebo group (Figure 5A, B). This result was more pronounced at the level of replicating virus, indicating efficient neutralization of cell-free virus in both treatment groups (Figure 5B). These trends were likewise reflected by the results of histopathological examinations of animals treated at 24 h p.i.. While the histological outcome of infection was similar between both treatment groups (Figure 6), semi-quantitative assessment of SARS-CoV-2 induced lesions revealed consistently higher scores for the mAb treated group compared to ensovibep. Interestingly, scores for inflammation in the mAb treated group were on average exceeding the scores obtained for the placebo group. These findings need to be interpreted knowing that 5/6 animals in the mAb treated group which had been scheduled for termination and analysis at day 3 had to be taken out of the study already on day 2 due to rapid onset of fulminant disease, which is reflected by these readouts.

**Figure 6:**
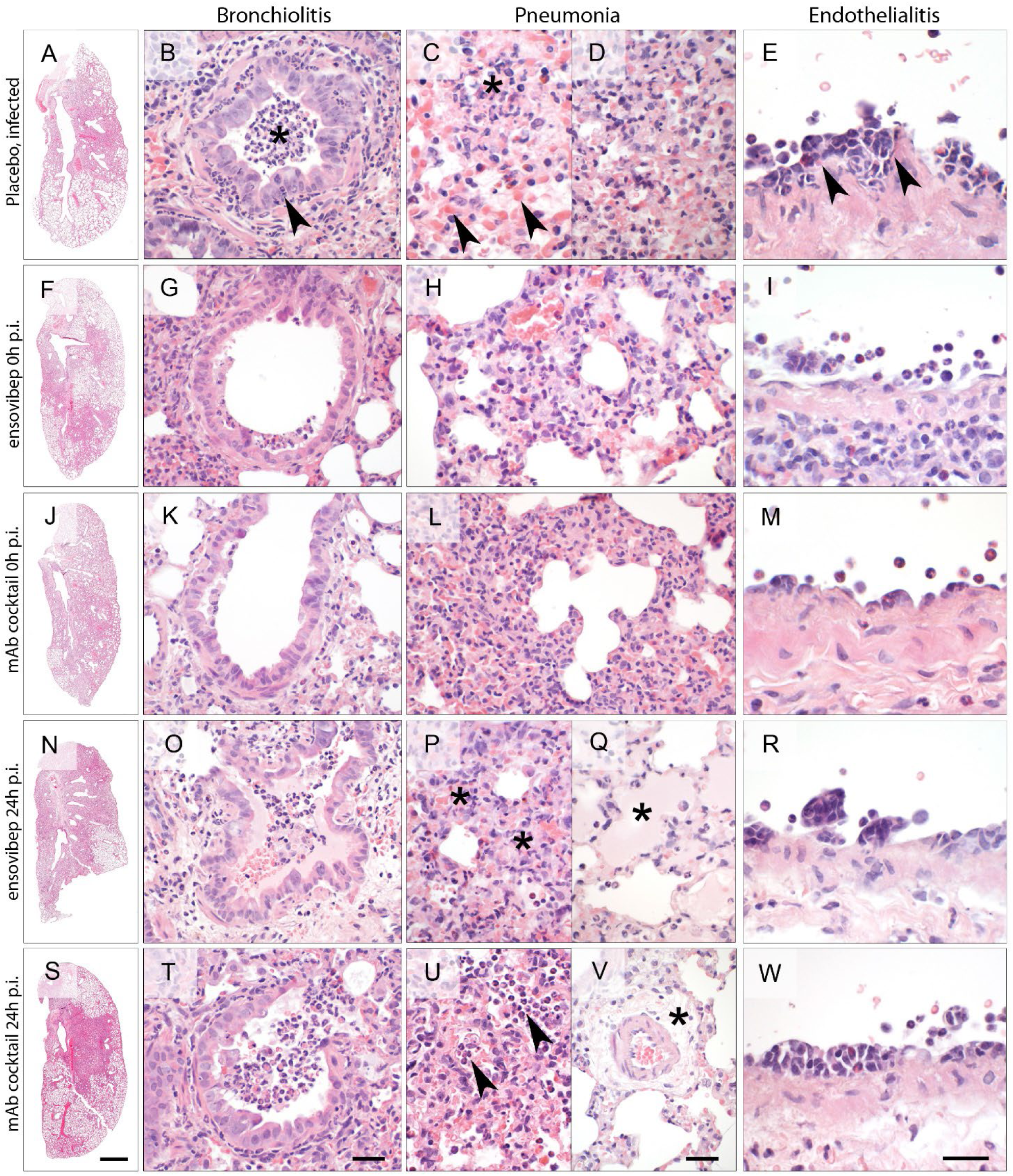
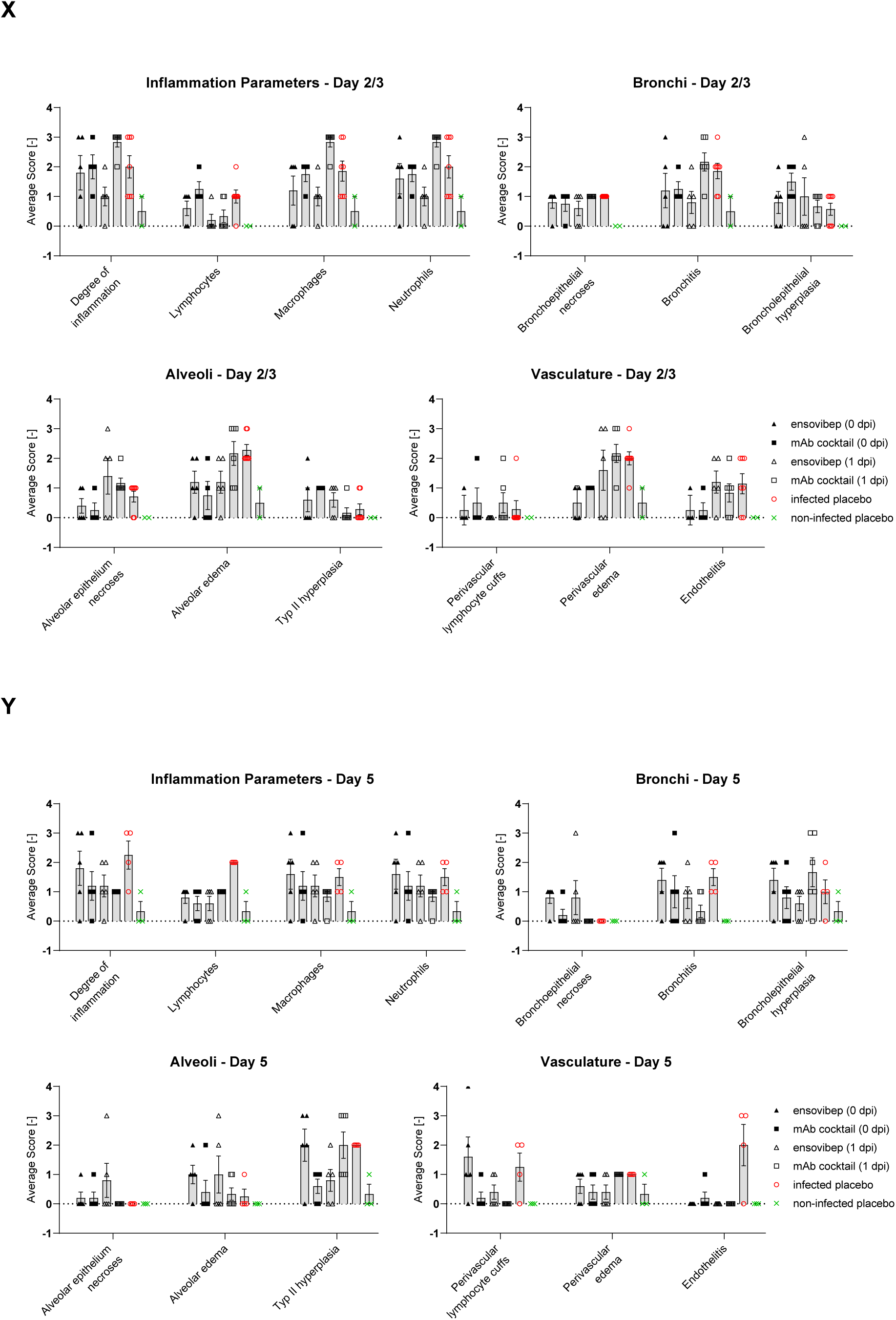
A-W. Lung histopathology of Roborovski dwarf hamsters at 2 or 3 days p.i. with SARS-CoV-2, hematoxylin and eosin stain. (A-E) Lungs of untreated hamsters at 3 days p.i. developed marked inflammation with lesion patterns as described earlier. (A) Whole slide scan revealing consolidation of approximately 60% of the left lung. (B) Untreated hamsters had moderate necro-suppurative and hyperplastic bronchiolitis with intraluminal accumulation of neutrophils and cellular debris (asterisk) as well as neutrophils transmigrating through the bronchial epithelium into the lumen (arrowhead). The lung parenchyma presented with a patchy distribution of acute necrosis (C, asterisk) with microvascular thrombosis (arrowheads) or (D) with areas of dense infiltration by macrophages and neutrophils. (E) Pulmonary blood vessels had mild to moderate endothelialitis. (F-I) In contrast, lungs of hamsters treated with ensovibep on the day of infection developed (F) moderately less consolidation of their lungs. (G) Bronchiolitis was milder with less inflammatory cell infiltrate compared to the untreated group. Neutrophils were mostly absent. (H) Alveolar walls were only moderately expanded by neutrophils and macrophages with less alveolar edema compared to untreated hamsters. (I) Endothelialitis was virtually absent with marginating neutrophils as only immune cells interacting with the vascular lining. (J-M) Hamsters treated with the antibody cocktail at the day of infection developed lesions that were similar to those as described for the ensovibep treated group. (N-W) In contrast, lungs of hamsters treated at 1 dpi had lesions similar to the untreated hamsters at that time, regardless of their treatments. (O, T): Both treatment groups developed moderate bronchiolitis similar to the untreated group. (P, U): Lung parenchyma were characterized by interstitial (asterisks) and alveolar (arrowheads) infiltration with neutrophils and macrophages with variable necrosis of alveolar epithelial cells. Additional lesions in both treatment groups included (Q) moderate to marked alveolar edema (asterisk), here shown for the ensovibep group, and (V) moderate interstitial edema (asterisk), here shown for the antibody group. (R, W): Both treatment groups developed moderate endothelialitis with monomorphonuclear infiltrates underneath detached endothelial cells, similar to the untreated group. Scale bars: A, F, J, N, S = 1 mm; B, G, K, O, T = 50 µm; C, D, H, L, P, Q, U, V = 20 µm; E, I, M, R, W = 20 µm Histopathologic lesions were scored semi-quantitatively and scores plotted as graphs for histologic signs of general inflammation and histologic parameters of bronchiolar, alveolar and vascular lesions at day 2/3 p.i (X) or day 5 p.i. (Y).

To account for possible differences in exposure, we performed pharmacokinetic analysis for both treatments. These assessments identified that overall, comparable exposures were achieved in non-infected hamster following i.p. administration. It was noted that, ensovibep achieved a higher maximal serum concentration (Cmax) and a shorter systemic half-life compared to the mAb cocktail (Supplementary Figure 9).

Considering the small size of the Roborovski dwarf hamster, failure of i.p. injection due to an accidental injection into body compartments other than the peritoneum may occur. We thus screened for animals which lacked a proper drug exposure in terminal serum samples and removed data of these animals from all other analyses (Supplementary Table 4).

Whole genome sequencing using virus RNA recovered from lungs and upper respiratory tract was performed to investigate whether SARS-CoV-2 escape mutants were selected under ensovibep treatment. Viral RNA from individual animals with higher viral load compared to other animals of the same treatment group was analyzed and no escape mutations affecting the ensovibep epitope located in the RBD were discovered (Supplementary Table 5).

## Discussion

Multiple strategies are urgently needed to combat the COVID-19 pandemic. Next to preventive vaccination approaches and small molecules, mAbs are showing therapeutic promise, based on highly potent virus inhibition and encouraging animal and clinical efficacy. However, manufacturing capacities are limiting a global supply and novel emerging variants of SARS-CoV-2 are an ever-present threat, as they may escape the antibodies generated during immunization or in response to therapeutics. A number of alternative molecules are being developed to complement and partially overcome these limitations.

In the present study, we provide the structural and functional analysis of ensovibep, a trispecific DARPin designed as a potential alternative to antibodies and other therapeutics^32,49–53^. The structural analysis provides insights into the mode of action, which enables low picomolar neutralizing activity against the currently most frequent SARS-CoV-2 mutations as well as recently identified variants. We measured the effect of ensovibep on a panel of single spike protein mutations which have been shown to be of concern because they may be associated with increased transmissibility, disease severity, or affect neutralization of some monoclonal- or polyclonal antibodies^27,54,55^. Among all mutations tested, only F486 substitutions caused a strong decrease in ensovibep potency when compared to the wild-type or reference virus. The effect of this mutation was also noted in the viral passaging study: sequencing of mutations allowing escape from inhibition by the monovalent RBD binder (R2, incorporated in ensovibep) identified F486L (Figure 3B). These findings are in line with our structural analysis (Figure 1F) showing that F486 is one of the key binding residues for the interaction of ensovibep with the RBD. Most importantly, F486 is a critical residue for the virus itself, allowing an efficient binding to the ACE2 receptor and thus cell infection. Therefore, mutations of the phenylalanine at position 486 will decrease the affinity between the RBD and human ACE2 receptor and lower the infectivity of the virus^17,56–58^ (Supplementary Figure 6). We thus expect that position F486 in the SARS-CoV-2 spike protein will remain conserved to maintain efficient binding to the human ACE2 receptor or that the virus might lose fitness if mutated at this position. So far, based on the global SARS-CoV-2 database sequences published in the GISAID database (https://www.gisaid.org/hcov19-variants/; visited November 2021), mutations in position F486 occur at very low frequencies.

A small reduction of neutralization potency observed for ensovibep and its single DARPin moieties for viruses bearing the N234Q mutation outside of the RBD might be explained by the impact of the mutation on the RBD conformational dynamics. An *in-silico* simulation study showed that this conserved glycosylation site, together with N165, might be involved in the stabilization of the RBD up-conformation. Since the epitope of ensovibep is exposed only in the up-conformation, a mutation in one of these glycosylation sites might affect its binding equilibrium, as indicated in our neutralization assays. The N234Q mutation might thus impact all protein binding scaffolds that are binding exclusively to the up-conformation of the RBD. By the same token, reduced affinity of the spike protein for the human ACE2 receptor was demonstrated elsewhere in *in vitro* assays^44^. Accordingly, mutations of the N165 and N234 amino acids have been observed only at low frequencies (<0.02%).

Some mutations that are not predicted to be key interaction residues for the three distinct RBD binders of ensovibep (e.g., E484K or Q493K), led to a reduction in potency for one or several of the RBD-binding monovalent DARPins, while the trispecific ensovibep molecule maintained full neutralization capacity. This demonstrates that the trispecific DARPin design of ensovibep, with cooperative binding of three distinct paratopes (Supplementary Figure 1), permits high neutralizing potency, even in the case when an individual monovalent DARPin domain exhibits decreased affinity (Figure 2A). This cooperative binding of multiple paratopes is a hallmark of the trispecific nature of ensovibep and differentiates the molecule from mAb candidates to allow full neutralization of highly mutated SARS-CoV-2 variants such as Omicron BA.1 and BA.2, that are substantially different from the original virus that the mAb was selected against^59^.

The high level of protection against viral escape mutations by ensovibep demonstrated in the virus challenge studies was also clearly apparent in a viral passaging experiment. The single mAbs and the monovalent DARPin binder were rapidly overcome by escape mutants whereas ensovibep maintained potency to an extent comparable to a clinically validated mAb cocktails.

Translatability of the observed *in vitro* activity of ensovibep against SARS-CoV-2 was evaluated in a COVID-19 model using the highly susceptible Roborovski dwarf hamsters. Using this *in vivo* model, we confirmed the therapeutic benefit of ensovibep, which displayed comparable outcomes to a clinically validated antibody cocktail (REGN10933 & REGN10987). In our comparison, we found evidence for a better performance of ensovibep in a late intervention scenario with prolonged survival of animals and reduced inflammation of the lungs. Potential reasons for this difference include differences in pharmacokinetics where ensovibep demonstrated a higher maximal concentration compared to the antibody cocktail (Supplementary Figure 9). Another possible explanation could be that ensovibep lacks an Fc-fragment when compared to antibodies stimulating pro-inflammatory immune responses mostly via their Fc-fragment. Regardless of this, we clearly demonstrate that ensovibep has great potential to prevent disease and eliminate the virus in a highly susceptible *in vivo* model under different treatment scenarios. The clinical translatability of these results is currently being investigated in the EMPATHY trial for the treatment of ambulatory COVID-19 patients.

In conclusion, ensovibep, has been shown to have highly potent neutralization against the currently most frequent SARS-CoV-2 variants due to its cooperative and complementary binding to a highly conserved epitope region on the spike RBD. *In vitro* and *in vivo* single agent efficacies closely match the performance of one of the best clinically validated mAb cocktails. In addition, the albumin binding domains of the molecule have been demonstrated to confer a plasma half-life compatible with single dose treatment. Translation of these preclinical findings into the clinic is currently under investigation and if successful, the *E. coli* based-manufacturing of the agent will allow rapid and large-scale production for global access to this alternative class of therapeutics as an addition to other treatment approaches for COVID-19.

## Data availability

The EM density maps for the SARS-CoV-2 spike ectodomain in complex with monovalent DARPin R2 (state 1 and state 2), have been deposited to the Electron Microscopy Data Bank under the accession codes EMD-11953 and EMD-11954, respectively. The monovalent DARPin and multivalent DARPin sequences, and pseudo-atomic models derived from molecular docking experiments, are available here, to allow the use of the data for non-commercial purposes: https://www.guidetopharmacology.org/GRAC/LigandDisplayForward?tab=structure&ligandId=11470

## Acknowledgements

S.R. & J.M. were supported by Swiss Federal Office for Civil Protection (Grants Nr. 353008564/Stm, 353008218/Stm, and 353008560/Stm to Olivier Engler and Stefan Kunz).

D.L.H. is funded by the European Union’s Horizon 2020 research and innovation program under the Marie Skłodowska-Curie grant agreement (No 842333) and holds an EMBO non-stipendiary long-term Fellowship (ALTF 1172-2018). Cryo-EM data processing was carried out on the Dutch national e-infrastructure with the support of the SURF Cooperative.

The authors also thank Dr. Gert Zimmer for the gift of the recombinant VSV (Institute of Virology and Immunology (IVI), CH-3147 Mittelhäusern, Switzerland, Department of Infectious Diseases and Pathobiology, Vetsuisse Faculty, University of Bern, CH-3012 Bern, Switzerland).

The expression plasmid for the SARS-CoV-2 spike protein was kindly provided by Dr. Giulia Torriani and Dr. Isabella Eckerle (Department of Medicine, University of Geneva, Switzerland).

We would like to thank Dr. Sylvie van der Werf for the supply of 2019-nCoV/IDF0372/2020 (National Reference Centre for Respiratory Viruses hosted by Institut Pasteur (Paris, France)). Strain 2019-nCoV/IDF0372/2020 was generously provided by Dr. X. Lescure and Pr. Y. Yazdanpanah from the Bichat Hospital.Additionally, we would like to thank William Lee, former board member of Molecular Partners - and the Virology group at Gilead Sciences for their helpful input.

We would like to thank the Centre for AIDS Reagents (National Institute for Biological Standards and Control, Herts, UK) for providing VeroE6/TMPRSS2 cells.

Lentivirus pseudotype investigations were performed independently by investigators at the US Food and Drug Administration, Center for Biologics Evaluation and Research as part of Therapeutics Research Team for the US government COVID-19 response efforts. The work was supported by US government research funds.

## Supplementary Materials for

## Materials and Methods

### Generation of His-tagged monovalent RBD binders and ensovibep

DARPin constructs selected and cloned as described in Walser et al.^31^ were transformed in *E.coli* BL21 cells, plated on LB-agar (containing 1% glucose and 50 μg/mL ampicillin) and then incubated overnight at 37°C. A single colony was picked into TB medium (containing 1% glucose and 50 μg/mL ampicillin) and incubated overnight at 37°C, shaking at 230 rpm. Fresh TB medium (containing 50 μg/mL ampicillin) was inoculated with 1:20 of overnight culture and incubated at 37°C at 230 rpm. At OD600 = 1.1 the culture was induced by addition of IPTG (0.5 mM final concentration) and incubated further for 5 h at 37°C and 230 rpm. Harvest was done by centrifugation (10 min, 5000 x g). After cell disruption by sonication, primary recovery was done by heat treatment for 30 min at 62.5°C and subsequent centrifugation (15 min, 12000 x g). 20 mM Imidazole and 1% Triton X-100 was added to the supernatant and the 0.22 µm filtered supernatant was further purified by immobilized metal affinity chromatography (IMAC) (HisTrap FF crude, Cytiva, Sweden) using the N-terminal His-tag and including a wash step with 1% Triton X-100 and a step elution with 250 mM Imidazole. In a subsequent step, the elution fraction of the IMAC step was applied on a size exclusion chromatography (Superdex 200, Cytiva, Sweden) and fractions of interest were pooled and concentrated. Finally, the concentrated sample was filtered through a 0.22 µm Mustang E filter for Endotoxin removal and sterile filtration and quality controlled.

### Cryo-electron microscopy

4 μl of purified S-ectodomain (9 μM) was mixed with 1 μl of 50 μM mono-DARPin R2, and incubated for 15 seconds at room temperature. 3 μl of sample was then dispensed on Quantifoil R1.2/1.3 200-mesh grids (Quantifoil Micro Tools GmbH) that had been freshly glow discharged for 30 s at 20 mA. Grids were blotted using blot force +2, for 5 s using Whatman No. 1 filter paper and immediately plunge-frozen into liquid ethane cooled by liquid nitrogen using a Vitrobot Mark IV plunger (Thermo Fisher Scientific) equilibrated to ∼95% relative humidity, 4°C. Movies of frozen-hydrated specimens were collected using Glacios Cryo-TEM (Thermo Fisher Scientific) operating at 200 keV and equipped with a Falcon 4 Direct Electron Detector (Thermo Fisher Scientific). For additional analysis of monovalent DARPin R2, 4 μl of purified S-ectodomain (18 μM) was mixed with 1 μl of 100 μM DARPin, and incubated for 60 s at room temperature. Grids were prepared as described above, and movies were collected using a Titan Krios Cryo-TEM (Thermo Fisher Scientific) operating at 300 keV and equipped with a Falcon 4 Direct Electron Detector (Thermo Fisher Scientific). All cryo-EM data were acquired using the EPU 2 software (Thermo Fisher Scientific) with a 30-degree stage tilt to account for preferred orientation of the samples. Movies were collected in electron counting mode at 92,000x (Glacios) or 75,000x (Titan Krios), corresponding to a pixel size of 1.1 Å/pix or 1.045 Å/pix over a defocus range of -1.25 to -2.5 μm.

### Image processing

Movie stacks were manually inspected and then imported in Relion version 3.1^60^. Drift and gain correction were performed with MotionCor2^61^, and GCTF^62^ was used to estimate the contrast transfer function for each movie. Particles were automatically picked using the Laplacian-of-Gaussian (LoG) algorithm and then Fourier binned (2 x 2) particles were extracted in a 160-pixel box. The extracted particles were subjected to two rounds of 2D classification, ignoring CTFs until the first peak. Using the ‘molmap’ command in UCSF chimera^63^, a SARS-CoV-2 spike structure (PDB ID: 6VSB)^64^ was used to generate a 50 Å resolution starting model for 3D classification. Particles selected from 2D classification were subject to a single round of 3D classification (with C1 symmetry). Particles belonging to the best classes were re-extracted unbinned in a 320-pixel box, 3D auto-refined (with C1 or C3 symmetry) and post-processed. Iterative rounds of per particle defocus estimation, 3D auto-refinement and post-processing were used to account for the 30-degree stage tilt used during data collection. When CTF refinement did not yield any further improvement in resolution, Relion’s Bayesian polishing procedure was performed on the particle stacks, with all movie frames included, followed by 3D auto-refinement and post-processing. Subsequently, additional rounds of per particle defocus estimation, 3D auto-refinement and post-processing were performed on the polished particles until no further improvement in resolution or map quality was observed. The nominal resolution for each map was determined according to the ‘gold standard’ Fourier shell correlation (FSC) criterion (FSC = 0.143) and local resolution estimations were performed using Relion. Map sharpening was performed using DeepEMhancer^65^ as implemented in COSMIC2^66^. To improve the quality of the mono-DARPin R2 density in the fully open spike reconstruction, a focused 3D classification approach was employed. Briefly, each particle contributing to the final C3-symmetry–imposed reconstruction was assigned three orientations corresponding to its symmetry related views using the “relion_particle_symmetry_expand” tool. A soft mask was placed over the map to isolate the mono-DARPin R2-bound RBD, and the symmetry-expanded particles were subjected to masked 3D classification without alignment using a regularization parameter (‘T’ number) of 20. Particles corresponding to the 3D class with the best resolved DARPin density were re-extracted in a 200-pixel box and centered on the mask used for focused classification. In conjunction with this, the signal for the protein outside the masked was subtracted. The re-extracted particles were then 3D auto-refined (with C1 symmetry) using local angular searches (1.8 degrees) and sharpened using DeepEMhancer^65^. Three copies of the locally refined map were aligned to the globally refined map using the UCSF Chimera ‘fit in map’ tool and resampled using the ‘vop resample’ command. Finally, a composite map was generated using the “vop add” command. An overview of the image processing workflows is shown in supplementary Figure 2A.

### Molecular modeling of mono and multivalent DARPin molecules

Homology models of monovalent DARPin molecules R1, R2 and R3 were generated with Rosetta^67–69^. The consensus designed ankyrin repeat domain PDB ID:2XEE was used as template. Mutations were introduced with RosettaRemodel with fixed backbone, and the structure was refined with RosettaRelax. Forty refined structures were clustered using RosettaCluster with 0.3 Å radius, and the lowest-energy model from the largest cluster served as the final model. The UCSF Chimera ‘fit in map’ tool was used to fit the monovalent DARPin R2 model into the cryo-EM map produced from focused refinement. This fitted model of DARPin R2, together with the RBD domain (PDB ID:6M0J) was further refined with Rosetta. The structure was pre-relaxed for docking and served as input for local, high-resolution docking with RosettaDock with fixed backbone. Five hundred models were generated and clustered with 1 Å radius (RosettaCluster). Two largest clusters were inspected and the lowest-energy model from more conserved group (i.e., with lower rigid-body perturbation from the input structure) was taken further for additional all-atom refinement with RosettaRelax, with protocol optimized for interfaces (InterfaceRelax2019). Fifty models were generated, and the lowest scoring model was selected. This model was used to describe the interactions between DARPin R2 and the RBD. The PDB file with the coordinates of the trimer of DARPin R2:RBD was used as an input structure for the conceptual modeling of ensovibep bound to the spike ectodomain as shown in Figure 1G. The linkers were generated using Rosetta modeling tools. Figures were generated using LigPlot^70^, UCSF Chimera^63^, UCSF ChimeraX^71^, PyMOL (The PyMOL Molecular Graphics System, Version 2.0, Schrödinger, LLC) and BioRender (BioRender.com).

### Generation of monoclonal antibodies

Publicly available sequences of variable domains from monoclonal antibodies were used to synthetize the corresponding cDNA fragments and cloned into a proprietary expression vector at Evitria AG (Schlieren, Switzerland). Generated vectors containing the constant immunoglobulin chains were used for transfection in Chinese hamster ovary cells by Evitria. Sterile filtered cell supernatants were purified via affinity purification with HiTrap MabSelect column followed by a size exclusion chromatography using HiLoad 26/600 Superdex 200 column in PBS pH7.4. Selected fractions were pooled and quality controlled (by SDS-PAGE, size exclusion chromatography and endotoxin measurement) before use in assays.

### VSV-SARS-CoV-2 pseudotype mutation-vector generation

Plasmid pCAGGS containing the Wuhan-hu-1 spike protein of SARS-CoV-2^31^ was used as a reference and as a template for generation of single and multiple spike protein mutants. Forward and reverse complementary primers encoding the mutation were synthesized by Microsynth (Balgach, Switzerland). High-fidelity Phusion polymerase (New England Biolabs, USA) was used for all DNA amplification.

Single mutations of the spike protein were generated via two PCR fragments of the spike ORF using high-fidelity Phusion polymerase (New England Biolabs, USA). The first fragment was generated via a generic forward primer (pCAGGS-5) annealing upstream of the spike ORF and the specific reverse primer encoding the mutation. The second fragment was generated using the specific forward primer encoding the mutation and a reverse primer (rbglobpA-R). The two fragments were gel-purified and used as input for an assembly PCR without addition of flanking primers.

For multi-mutation spike proteins, a complementary pair of primers (forward and reverse) encoding each mutation was designed. Fragment 1 was generated with forward primer pCAGGS-5 and reverse primer encoding mutation 1. Fragment 2 was generated using forward primer encoding mutation 1 and reverse primer encoding mutation 2. All subsequent fragments were generated analogously. DNA fragments were gel-purified and mixed in equimolar amounts. This mix was used for re-assembly of the full spike ORF using outer primers pCAGGS-5 (GGTTCGGCTTCTGGCGTGTGACC) and rbglobpA-R (CCCATATGTCCTTCCGAGTG).

For both single as well as multi-mutation spike protein, the full-length spike ORF was isolated from an agarose gel, digested by restriction enzymes NheI/EcoRI and inserted into the pCAGGS vector backbone. The correct sequence was verified via sequencing the whole ORF of the spike protein by Microsynth (Balgach, Switzerland).

### VSV-SARS-CoV-2 pseudotype neutralization assay for mutational analysis

The pseudotype viral system was based on the recombinant VSV*DELG-Luc vector in which the glycoprotein gene (G) had been deleted and replaced with genes encoding green fluorescent protein and luciferase. For the neutralization assay of ensovibep, MP0420 or their his-tagged variants ALE049/ALE070, an initial dilution of the compounds was followed by three-fold dilutions in quadruplicates in DMEM-2 % [vol/vol] FCS supplemented with 20 μM human serum albumin (CSL Behring). The mixture was mixed with an equal volume of DMEM-2 % FCS containing 250 infectious units (IU) per well of SARS-CoV-2 pseudoviruses and incubated for 90 min at 37°C. The mix was inoculated onto Vero E6 cells in a clear bottom white walled 96-well plate during 90 min at 37°C. The inoculum was removed and fresh medium added, and cells further incubated at 37°C for 16 h. Cell were lysed according to the ONE-Glo™ luciferase assay system (Promega, Madison, US) and light emission was recorded using a Berthold TriStar LB941 luminometer. The raw data (relative light unit values) were exported to GraphPad Prism v8.4.3. IC50/IC90 were modelled with a nonlinear regression fit with settings for log (inhibitor) vs normalized response curves. Data points are plotted by the mean ± SEM (standard error of mean) of quadruplicate data.

### SARS-CoV-2 lentivirus-based pseudovirus neutralization assay

The neutralizing activity of therapeutic antibodies against SARS-COV-2 variants was measured using lentiviral particles pseudotyped with spike proteins of SARS-COV-2 variants, as previously described^72^. Briefly, pseudoviruses bearing the spike proteins and carrying a firefly luciferase ^73^ reporter gene were produced in 293T cells by co-transfection of pCMVΔR8.2, pHR’CMVLuc and pCDNA3.1-spike variants. Plasmids encoding human codon-optimized spike genes with the desired mutations were purchased (GenScript, Piscataway, NJ). Supernatants containing pseudoviruses were collected 48 h post-transfection, filtered through a 0.45 μm low protein binding filter, and stored at -80oC. Pseudovirus titers were measured by infecting 293T-ACE2.TMPRSS2s cells for 48 h prior to measuring luciferase activity (luciferase assay reagent, Promega, Madison, WI). For neutralization assays, pseudoviruses with titers of approximately 106 relative luminescence units (RLU)/ml were incubated with serially diluted DARPin for two h at 37°C before adding the pseudovirus and DARPin mixtures (100 μl) onto 96 well plates pre-seeded one day earlier with 3.0 x 104 293T-ACE2.TMPRSS2s cells/well. Pseudovirus infection was scored 48 h later by measuring luciferase activity. The DARPin concentration causing a 50% reduction of RLU compared to control (ID50) was reported as the neutralizing DARPin titer. Titers were calculated using a nonlinear regression curve fit (GraphPad Prism software Inc., La Jolla, CA). The ratio of the neutralizing DARPin titer of the variant compared to the neutralizing DARPin titer of wild-type reference was calculated. The D614G mutation background was used for the variants and the reference virus. The mean titer from at least two independent experiments with intra-assay duplicates was reported as the final titer. This work was performed independently by investigators at the US Food and Drug Administration, Center for Biologics Evaluation and Research as part of Therapeutics Research Team for the US government COVID-19 response efforts.

### SARS-CoV-2 lentivirus-based pseudovirus neutralization assay (Setup 2)

Neutralizing activity was measured in an assay that utilized lentiviral particles pseudotyped with full-length SARS-CoV-2 Spike protein and containing a firefly luciferase (Luc) reporter gene for quantitative measurements of infection by relative luminescence units (RLU). The backbone vector used in pseudovirus creation, F-lucP.CNDOΔU3, encodes the HIV genome with firefly luciferase replacing the HIV env gene. A codon-optimized version of the full-length spike gene of the Wuhan-1 SARS-CoV-2 strain (MN908947.3; GenScript) was cloned into the Monogram proprietary env expression vector, pCXAS-PXMX, for use in the assay. The D614G spike mutation was introduced into the original Wuhan sequence by site-directed mutagenesis. Sequences of the spike gene and expression vector were confirmed by full-length sequencing using Illumina MiSeq NGS.

Pseudovirus stock was produced in HEK 293 cells via a calcium phosphate transfection using a combination of spike plasmid (pCXAS-SARS-CoV-2-D614G) and lentiviral backbone plasmid (F-lucP.CNDOΔU3). Transfected 10 cm^2^ plates were re-fed the next day and harvested on Day 2 post transfection. The pseudovirus stock (supernatant) was collected, filtered and frozen at -70°C in single use aliquots. Pseudovirus infectivity was screened at multiple dilutions using HEK293 cells transiently transfected with ACE2 and TMPRSS2 expression vectors. RLUs were adjusted to ∼ 50,000 for use in the neutralization assay. Neutralization was performed in white 96-well plates by incubating pseudovirus with 10 serial threefold dilutions of samples for one hour at 37°C.

HEK293 target cells, which had been transfected the previous day with ACE2 and TMPRSS2 expression plasmids, were detached from 10 cm^2^ plates using trypsin/EDTA and re-suspended in culture medium to a final concentration that accommodated the addition of 10,000 cells per well. Cell suspension was added to the serum-virus mixtures and assay plates were incubated at 37°C in 7% CO_2_ for 3 days. On the day of assay read, Steady Glo (Promega) was added to each well. Reactions were incubated briefly and luciferase signal (RLU) was measured using a luminometer. Neutralization titers represent the inhibitory concentration (IC) of samples at which RLUs were reduced by either 50% (IC_50_) or 90% (IC_90_) compared to virus control wells (no sample wells). Data of single runs are represented. This work was performed independently by investigators at Monogram Biosciences, CA, US, for the US government COVID-19 response efforts.^74^

### Cells and pathogenic virus

Vero E6 cells (kindly provided by Prof. Volker Thiel, University of Bern, Switzerland) were passaged in Minimum Essential Medium (MEM) (Cat N° M3303) containing 10% fetal bovine serum (FBS) and supplements (2 mM L-Glutamine, 1% Non-essential amino acids, 100 units/ml Penicillin, 100 μg/ml Streptomycin, 0.06% Sodium bicarbonate, all from Bioswisstec, Schaffhausen, Switzerland) at 37°C, >85% humidity and 5% CO_2_. Vero E6/TMPRSS2 cells^75,76^ obtained from the Centre For AIDS Reagents (National Institute for Biological Standards and Control) were passaged in Dulbecco’s Modified Eagle Medium (DMEM) (Cat N° M1452) containing 10% fetal bovine serum (FBS) and supplements (2 mM L-Glutamine, 1% Non-essential amino acids, 100 U/mL Penicillin, 100 μg/mL Streptomycin, 0.06% Sodium bicarbonate and 2% Geneticin G418, all from Bioswisstec, Schaffhausen, Switzerland) at 37°C, >85% humidity and 5% CO_2_.

SARS-CoV-2 (2019-nCoV/IDF0372/2020), kindly provided by Dr. Sylvie van der Werf from the National Reference Centre for Respiratory Viruses hosted by Institut Pasteur (Paris, France) was propagated in Vero E6 cells in MEM containing 2% FBS and supplements (2%-FBS-MEM) at 37°C, >85% humidity and 5% CO_2_. SARS-CoV-2 variants (B.1.1.7, B.1.351 and P.1) were provided from University Hospital of Geneva, Laboratory of Virology ^25^ and propagated in Vero E6/TMPRSS2 cells in DMEM containing 2% FBS and supplements (2%-FBS-DMEM) at 37°C, >85% humidity and 5% CO_2_. Viral titer was determined by standard plaque assay, by incubating 10-fold serial dilutions of the virus for 1 h at 37°C on a confluent 24-well plate with Vero E6 cells. Then inoculum was removed and 1 mL overlay medium (20 ml Dulbecco’s Modified Eagle’s Medium, 5 ml FBS, 100 U/mL Penicillin, 100 μg/mL Streptomycin, and 25 ml Avicel rc581) was added. After 3 days incubation at 37°C the overlay was removed and the plates stained with crystal violet solution (spatula tip (∼4 mg) crystal violet powder (Sigma Aldrich) solved in 30 ml 37% formalin and 120 mL PBS (Sigma Aldrich).

### Viral passaging experiment with authentic SARS-CoV-2

Virus escape studies were adapted from a previously published protocol by Baum et al.^27^. Briefly, 1:5 serial dilutions of DARPin molecules and monoclonal antibodies from 100 μg/mL to 0.032 μg/mL were prepared in Minimum Essential Medium (MEM) containing 2% FBS, supplements and 10 μM human serum albumin (HSA; CSL Behring, Switzerland; 2%-FBS-MEM + HSA). 500 µL of virus suspension containing 1.5 x 10^6^ plaque forming units (pfu) SARS-CoV-2 in 2%-FBS-MEM + HSA were mixed with 500 μL of serially diluted DARPin molecules or monoclonal antibodies and subsequently incubated for 1 hour at 37°C. The mixtures were then transferred to confluent Vero E6 cells in 12 well plates and incubated for 4 days at 37°C, >85% humidity and 5% CO_2_. Each culture well was assessed for cytopathic effect (CPE) by microscopy. Supernatant was removed from wells with the highest DARPin or antibody concentrations showing significant CPE (≥20%) and used for total RNA extraction and further passaging. For subsequent rounds of passaging, remaining 900 μL supernatant of selected wells was diluted in 4 mL in 2%-FCS-MEM + HSA and from the 4.9 mL, 500 μL mixed with serial dilutions of DARPin molecules or antibodies, incubated and the mixture transferred to 12-well plates with fresh Vero E6 cells as described above. Cell culture wells were assessed for CPE again after 4 days and the supernatant of wells with highest DARPin or antibody concentrations with evident viral replication (CPE) harvested and used for additional passages. A total of 4 passages were performed this way.

### Deep sequencing of viral passages

RNA of the cell culture supernatant was extracted using the RNeasy Universal Plus kit (Qiagen, Basel, Switzerland) according to the manufacturer’s protocol. 10.5 µL of the extract was reverse transcribed using Superscript VILO (ThermoFisher Scientific, Reinach, Switzerland) following the manufacturer’s instructions. Barcoded libraries were prepared on the Ion Chef Instrument (ThermoFisher Scientific) using the Ion AmpliSeq SARS-CoV-2 Research Panel (ThermoFisher Scientific). 8-16 barcoded samples were pooled and loaded on one Ion 530 chip using the Ion Chef Instrument (ThermoFisher Scientific) and sequenced on the Ion S5 System with 550 flows.

The resulting BAM files were converted to fastq format using Samtools 1.10^77^ and subjected to adapter and quality trimming using Trimmomatic 0.39^78^ (options: ILLUMINACLIP:adapters.fasta:2:30.10, LEADING: 3, TRAILING: 3, SIDINGWINDOW:4:15, MINLEN:36). Reads were aligned to the SARS-CoV-2 reference genome (NC_045512.2) using bwa 0.7.17^79^ and variants were determined using LoFreq v2.1.5^80^. Variants were filtered for a minimal depth (DP) of 400X and a minimal allele frequency (AF) of 3% using bcftools 1.10^77^. Functional annotation of the variants was performed using SNPEff 5.0^81^. Variants were visualized in R 3.6.1 using ComplexHeatmap 2.2^82^.

### Virus neutralization of authentic wild type and variants of SARS-CoV-2 determined by Cell Titer-Glo

Virus neutralization capacity of mono-valent DARPin candidate and multispecific DARPin molecules was determined for 100 TCID_50_ SARS-CoV-2 variants from lineage B.1.1.7 (H69_V70del, Y145del, N501Y, A570D, D614G, P681H, T716I, S982A, D1118H), B.1.351 (L18F, D80A, D215G, L242_L244del, T302T, K417N, E484K, N501Y, D571D,D614G, A701V) and P.1 (L18F, T20N, P26S, D138Y, R190S, K417T, E484K, N501Y, D614G, H655Y, T1027I, V1176F) in reference to a wild-type French isolate (with the following differences to the Wuhan wild-type: V367F; E990A) by measuring ATP levels of protected cells in a cell viability assay. DARPin molecules were serially diluted 1:4 from 40 nM to 2.4 pM (in triplicates) in 100 μL cell culture medium (2%-FBS-DMEM) supplemented with 10 μM HSA in 96 well plates. The diluted DARPin antivirals were mixed with 100 TCID_50_ SARS-CoV-2 in 100 μL 2%-FBS-MEM + HSA and incubated for 1 h at 37°C. DARPin/virus mixtures (200 μL) were transferred onto confluent Vero E6/TMPRSS2 cells. The controls consisted of cells exposed to virus suspension only, to determine maximal cytopathic effect and of cells incubated with medium only, to determine baseline cell viability. The plates were incubated for 3 days at 37°C, >85% humidity and 5% CO_2_. Cell viability was determined by removing 100 μL supernatant from all wells and adding 100 μL CellTiter-Glo reagent to the cells as described in the manufacturers protocol (CellTiter-Glo® Luminescent Cell Viability Assay, Promega, Madison, USA). After 2 minutes shaking on an orbital shaker, lysis of the cells during 10 min and transfer to an opaque-walled plate at room temperature, luminescence was read using a GloMax instrument (Promega).

### Surface plasmon resonance (SRP) affinity determination of ensovibep and individual RBD-binding domains

SPR assays were used to determine the binding affinity of monovalent DARPin as well as multivalent DARPin molecules to the spike protein of SARS-CoV-2. All SPR data were generated using a Bio-Rad ProteOn XPR36 instrument with PBS-T (0.005% Tween20) as running buffer. A new neutravidin sensor chip (NLC) was air-initialized and conditioned according to Bio-Rad manual.

Monovalent DARPin molecules R1, R2, R3: Chemically biotinylated (via lysines) SARS-CoV-2 spike protein 20 (Sino Biologics) was captured to ∼3400 RUs (30 μg/mL, 30 μL/min, 300 s). Two buffer injections (100 μL/min, 60 s) followed by two 12.5 mM NaOH regeneration steps (100 μL/min, 18 s) were applied before the first injections. Mono-domain DARPin proteins were injected (at 50/16.7/5.6/1.9/0.6 nM) for 180 s at 100 μL/min for association and dissociation was recorded for 3600 s (at 100 μL/min). The ligand was regenerated with a 12.5 mM NaOH pulse (100 μL/min, 18 s). The data was double referenced against the empty surface and a buffer injection and fitted according to the 1:1 Langmuir model.

Multivalent DARPin molecules: Avi-tagged biotinylated SARS-CoV-2 S protein (Acro Biosystems) was captured to ∼1200 RUs (1.33 ug/mL, 30 μl/min, 300 s) on a precoated neutravidin chip (NLC). Two buffer injections (100 μL/min, 60 s) followed by three 12.5 mM NaOH regeneration steps (100 μL/min, 18s) were applied before the first injections. One single concentration of 20 nM of ensovibep was injected for 180 s at 100 μL/min for association and dissociation was recorded for 36’000 s (at 100 μL/min). The data was double referenced against the empty surface and a buffer injection. Due to avidity gain, no significant dissociation could be recorded during the measured time.

### Surface plasmon resonance (SRP) affinity determination of wt-RBD and RBD F486V to ACE2

SPR assays were used to determine the binding affinity of wt-RBD as well as RBD-F486V human ACE2 protein. SPR data were generated using a Bruker Sierra SPR-32 Pro instrument with PBS-T (0.005% Tween20) as running buffer. A Bruker biotin tag capture sensor chip (BTC) was initialized and conditioned according to Bruker manual.

Avi-tagged biotinylated monomeric human ACE2 (Acro Biosystems) was captured to ∼170 RUs (3.3 μg/mL, 10 μL/min, 60 s). SARS-CoV-2 S protein RBD (wt, Acro biosystems, 500nM-0.229nM, threefold dilution series) and SARS-CoV-2 S protein RBD-F486V (in-house produced, 1500nM-0.229nM, threefold dilution series) were injected for 240 s at 25 μL/min for association and dissociation was recorded for 300 s (at 25 μL/min). After each injection, a 15min pause was performed to ensure full dissociation of analyte from the ligand. The data was double referenced against the empty surface and a buffer injection and fitted according to the 1:1 Langmuir model.

### Roborovski dwarf hamster model for the assessment of antiviral potency of ensovibep on wild type SARS-CoV-2 and the B.1.1.7 (alpha) variant

#### Materials and Methods

##### 1. Cells and viruses

For in vivo experiments, SARS-CoV-2 isolates BetaCoV/Germany/BavPat1/2020^83^ and BetaCoV/Germany/ChVir21652/2020 (B.1.1.7) were grown on Vero E6 cells and whole genome sequenced prior to infection experiments to confirm genetic integrity. Particularly the presence and integrity of the furin cleavage site in the majority of the virus population was confirmed. All virus stocks were titrated on Vero E6 cells prior to infection.

##### 2. Animals and infection

A total of 120 female and male Roborovski dwarf hamsters (Phodopus roborovskii) was used for infection experiments. Animals were housed in groups of 3 to 6 animals of the same sex in individually ventilated GR900 cages (Tecniplast, Buguggiate, Italy) and provided with food and water ad libitum and bountiful enrichment (Carfil, Oud-Turnhout, Belgium). Infection was performed by intranasal administration of 1×10^5^ pfu SARS-CoV-2 in 20 µL cell culture medium under general anesthesia^38^. All animal procedures were performed in accordance with relevant institutional and legal regulations and approved by the responsible state authority, Landesamt für Gesundheit und Soziales Berlin, Germany, permit number G 0086/20.

##### 3. Treatment

DARPin molecules and monoclonal antibodies were administered intraperitoneally in sterile PBS. The final drug concentration was adjusted based on the desired dose and respective animal weight to a 100 µL injection volume. For intraperitoneal administration the animal was fixed by grasping the neck skin and the back skin between thumb and fingers. Subsequently, the hand was turned over so that the animal rests with its back in the palm of the hand. The head of the animal was kept downwards to prevent injection/damage in/of the organs and the needle was inserted left of the median line in the groin area, between the 4th and the 5th mammary gland/nipple. Finally, the needle was removed in a smooth motion. All animals in this study were treated once at the indicated time point, 0, 6 or 24 hours post infection.

##### 4. Experimental groups

From a total of 120 Roborovski dwarf hamsters, 54 were used to determine dose and time dependency of treatment success. In these cohorts, 6 animals per group were infected with 1×10^5^ pfu SARS-CoV-2 WT (BetaCoV/Germany/BavPat1/2020) and treated with either 3, 10 or 20 mg/kg ensovibep at the time of infection, with 1 or 20 mg/kg 6 h post infection, or with 10 mg/kg 24h post infection, a placebo (PBS) treatment group with 6 animals was also included in each of three studies performed for this purpose. (Suppl. Figure 7)

To compare efficacy of ensovibep and Regeneron antibody cocktail treatment, 60 animals were infected with 1×10^5^ pfu SARS-CoV-2 variant B.1.1.7 (BetaCoV/Germany/ChVir21652/2020). Subjects were divided into groups of 12 animals and treated with 10 mg/kg ensovibep, 10 mg/kg Regeneron mAb cocktail or a placebo (PBS) at the time of infection or with 10 mg/kg ensovibep or 10 mg/kg Regeneron mAb cocktail 24 h post infection. An additional 6 animals served as non-infected control group.

In all animal experiments performed in this study, half of each respective group was scheduled for take out at 3 dpi, the other half was to be terminated at 5 dpi. In some of the experiments, several animals had to be terminated at time points other than these for humane reasons. Defined humane endpoints included a body temperature < 33°C, body weight loss > 15% together with signs of respiratory distress, body weight loss > 20% or a combination of these factors. Animals were monitored at least twice a day to prevent any prolonged suffering.

##### 5. Virological analysis

RNA was extracted from throat swabs and lung tissue using the innuPREP Virus DNA/RNA Kit (Analytic Jena, Jena, Germany). Viral RNA was quantified using a one-step RT qPCR reaction with the NEB Luna Universal Probe One-Step RT-qPCR (New England Biolabs, Ipswich, MA, USA) and the 2019-nCoV RT-qPCR primers and probe (E_Sarbeco)^84^ on a StepOnePlus RealTime PCR System (Thermo Fisher Scientific, Waltham, MA, USA) according to the manufacturer’s instructions. Standard curves for absolute quantification were generated from serial dilutions of SARS-CoV-2 DNA obtained from a full-length virus genome cloned as a bacterial artificial chromosome and propagated in E. coli. Duplicate 10-fold serial dilutions were used to determine replication competent virus titers on confluent layers of Vero E6 cells. To this end, serial dilutions of lung tissue homogenates were made and incubated on Vero E6 monolayers for 2 hours at 37 °C. Cells were washed and overlaid with semi-solid cell culture medium containing 1.5% microcrystalline cellulose (Avicel) and incubated for 48 h at 37 °C after which plates were fixed with 4% formalin and stained with 0.75% crystal violet for plaque counting.

##### 6. Histology

For histopathology, the left lung lobe was carefully removed, immersion-fixed in formalin, pH 7.0, for 48 h, embedded in paraffin, and cut in 2 μm sections. Slides were stained with hematoxylin and eosin (HE) after dewaxing in xylene and rehydration in decreasing ethanol concentrations. Lung sections were microscopically evaluated in a blinded fashion by a board-certified veterinary pathologist to assess the character, distribution and severity of pathologic lesions using lung-specific inflammation scoring parameters as described for other lung infection models before. Three different scores were used that included the following parameters: (1) lung inflammation score including severity of (i) interstitial pneumonia (ii) bronchiolitis, (iii) necrosis of bronchial and alveolar epithelial cells, and (iv) hyperplasia of alveolar epithelial type II cells as well as (v) hyperplasia of bronchial epithelial cells; (2) immune cell infiltration score taking into account the presence of (i) neutrophils, (ii) macrophages, and (iii) lymphocytes in the lungs as well as (iv) perivascular lymphocytic cuffing; and (3) edema score including (i) alveolar edema and (ii) perivascular edema. HE-stained slides were analyzed and images were taken using an Olympus BX41 microscope with a DP80 Microscope Digital Camera and the cellSens™ Imaging Software, version 1.18 (Olympus Corporation, Münster, Germany). For the display of overviews of whole lung lobe sections, slides were automatically digitized using the Aperio CS2 slide scanner (Leica Biosystems Imaging Inc., Vista, CA, USA), and image files were generated using the Image Scope Software (Leica Biosystems Imaging Inc.). The percentages of lung tissues affected by inflammation were determined histologically by an experienced board certified experimental veterinary pathologist (O.K.) as described previously ^85^. Lung inflammation scores were determined as absent, (1) mild, (2) moderate or (3) severe and quantified as described previously ^85^. Immune cell influx scores and edema scores were rated from absent to, (1) mild, (2) moderate, or (3) severe.

##### 7. Whole genome sequencing of SARS-CoV-2 isolated from treated hamsters

Following RNA extraction from swabs and lung samples, libraries were prepared and sequenced using Illumina technology (Illumina, San Diego, California, USA). For library preparation, a multiplexed amplicon-based whole-viral-genome approach using the NEBNext® ARTIC SARS-CoV-2 Library Prep Kit (Illumina®) was employed (New England Biolabs, Ipswich, Massachussets, USA). Briefly, this approach relies on cDNA synthesis from total RNA and amplification of target SARS-CoV-2 cDNA using the V3 ARTIC primers; these amplicons then undergo the usual library preparation steps for Illumina sequencing (end repair, adaptor ligation and PCR enrichment). Quantification of enriched sequencing libraries was performed using the NEBNext® Library Quant Kit for Illumina® (New England Biolabs, Ipswich, Massachussets, USA). Libraries were then pooled and sequenced on an Illumina Miseq System (Illumina, San Diego, California, USA).

The generated Illumina sequencing data were processed with Trimmomatic v.0.39^78^ and mapped against genome reference MT270101.1, using the Burrows-Wheeler aligner v.0.7.17^79^. Mapping statistics were generated using Samtools v1.10^86^ and alignments were visualized using IGV v2.9.4 for Linux^87^. For detection of single-nucleotide polymorphisms (SNPs), Freebayes, a Bayesian genetic variant detector was used. All SNPs with a minimum mapping quality of 5, minimum count of 3 and minimum fraction of 0.01 were considered. Consensus sequences for each sample were obtained using BCFtools. All SNP-containing open reading frame (ORFs) sequences were extracted from these consensus genomes and translated using the Expasy^88^⁸. Translate tool. The resulting protein sequences were then aligned to the corresponding reference protein sequences using the Expasy⁸ SIM Protein Alignment tool. For SNPs that resulted in amino acid substitutions, their possible effect on protein function was gauged using two predictors: PROVEAN Protein^89 90^ and SIFT^91^. Results from both predictors were taken into account, except on instances where the SIFT predictor could not resolve the proposed substitution or made “low confidence” predictions, then PROVEAN’s prediction was prioritized as its protein database is larger and newer.

### Hamster pharmacokinetic study

Single intraperitoneal injections of 10 mg/kg were administered to female hamsters. Fifteen animals were enrolled in each study (n=3 per time point). Blood was sampled from individual animals at 2 h, 24 h, 48 h, 72 h and 168 h post administration and processed to serum. MP0420 serum concentrations were determined by sandwich ELISA using an anti-DARPin antibody as capture reagent and biotinylated RBD and HRP conjugated Streptavidin as detection reagent and quantified against a standard curve. Serum concentrations for detection of both antibodies REGN10933 and REGN10987 were determined by sandwich ELISA using an anti-IgG antibody as capture reagent and biotinylated RBD and HRP conjugated Streptavidin as detection reagent and using a standard curve.

Pharmacokinetic parameters were determined with non-compartmental analyses using the software Phoenix WinNonLin (Certara, Princeton, USA) or GraphPadPrism (GraphPad Software, La Jolla,USA). For the *in vivo* efficacy study, terminal bleed samples were collected at 2, 3 or 5 days p.i. according to study description.

## Supplementary Figures

**Supplementary Figure 1:**
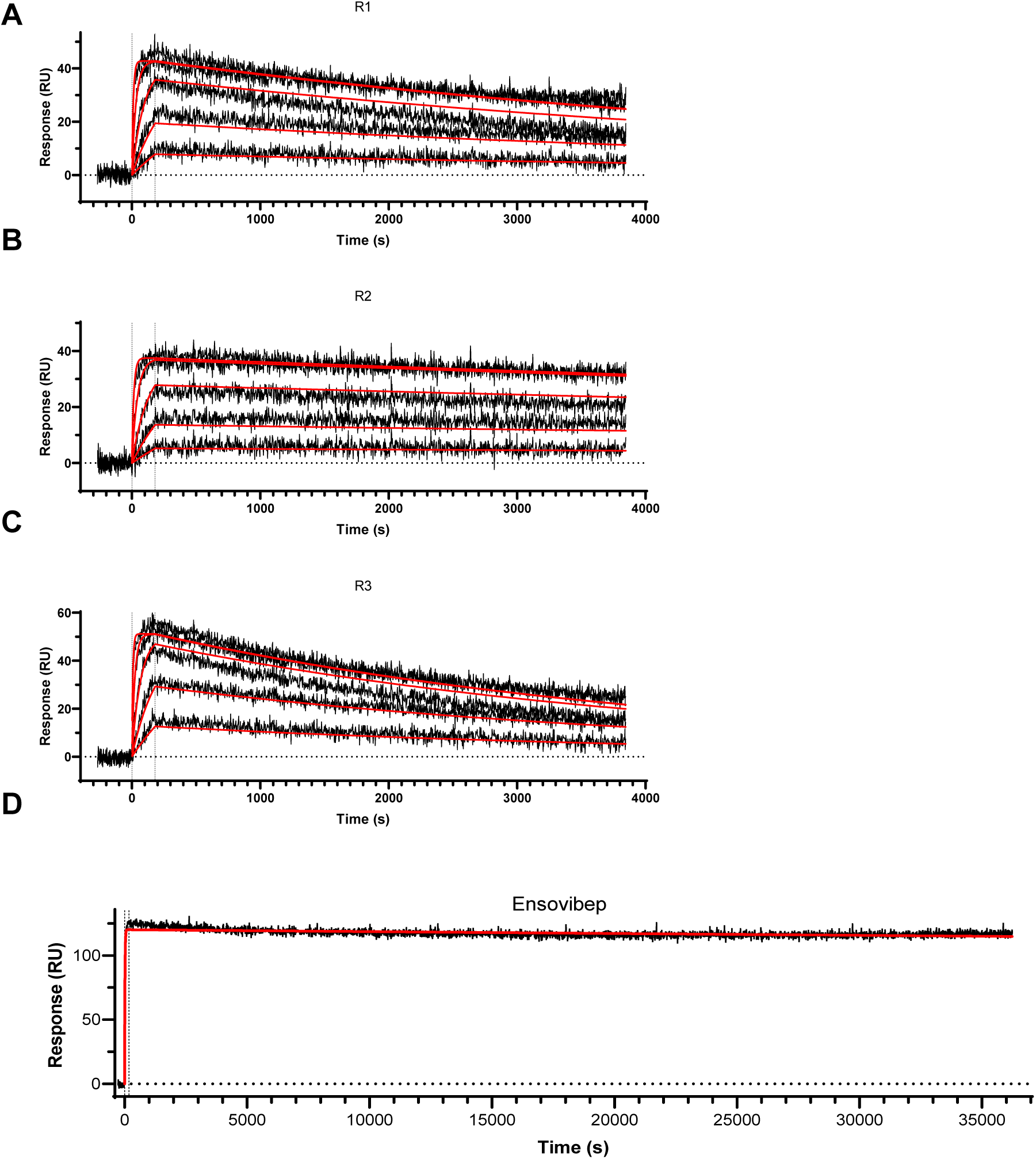
A-C) Surface plasmon resonance (SPR) sensorgrams of the monovalent DARPin modules (R1, R2, R3), incorporated in ensovibep binding to immobilized trimeric spike protein. DARPin concentrations for A-C: 50/16.67/5.56/1.85/0.62 nM. Determined K_D_ values: A) 80 pM, B) 30 pM, C) 90 pM. D) SPR sensorgram of ensovibep binding to immobilized spike protein. Off-rate was measured over 10 h and no physical off-rate could be determined by SPR due to very strong avidity of the three interlinked RBD binding modules.

**Supplementary Figure 2:**
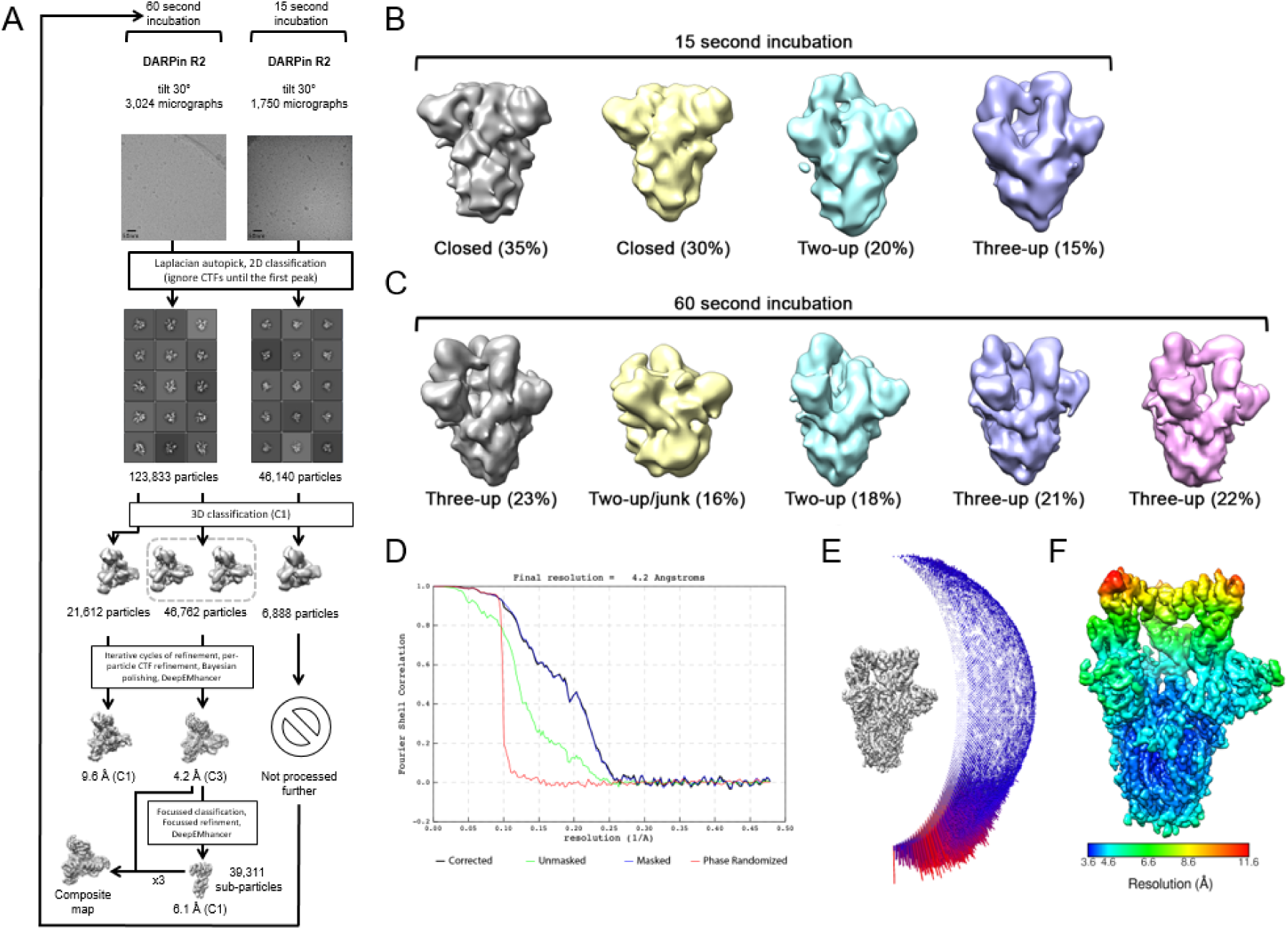
Single-particle cryo-EM data processing. A) Single-particle cryo-EM image processing workflow for the monovalent DARPin R2 data collections. B) 3D classes obtained from spike ectodomains incubated with monovalent DARPin R2 for 15 seconds, and C) for 60 seconds. D) Gold-standard Fourier shell correlation (FSC) curve generated from the independent half maps contributing to the 4.2 Å resolution density map. E) Angular distribution plot of the final C3 refined EM density map. F) The EM density map of the spike ectodomain bound to three copies of monovalent DARPin R2, colored according to local resolution.

**Supplementary Figure 3:**
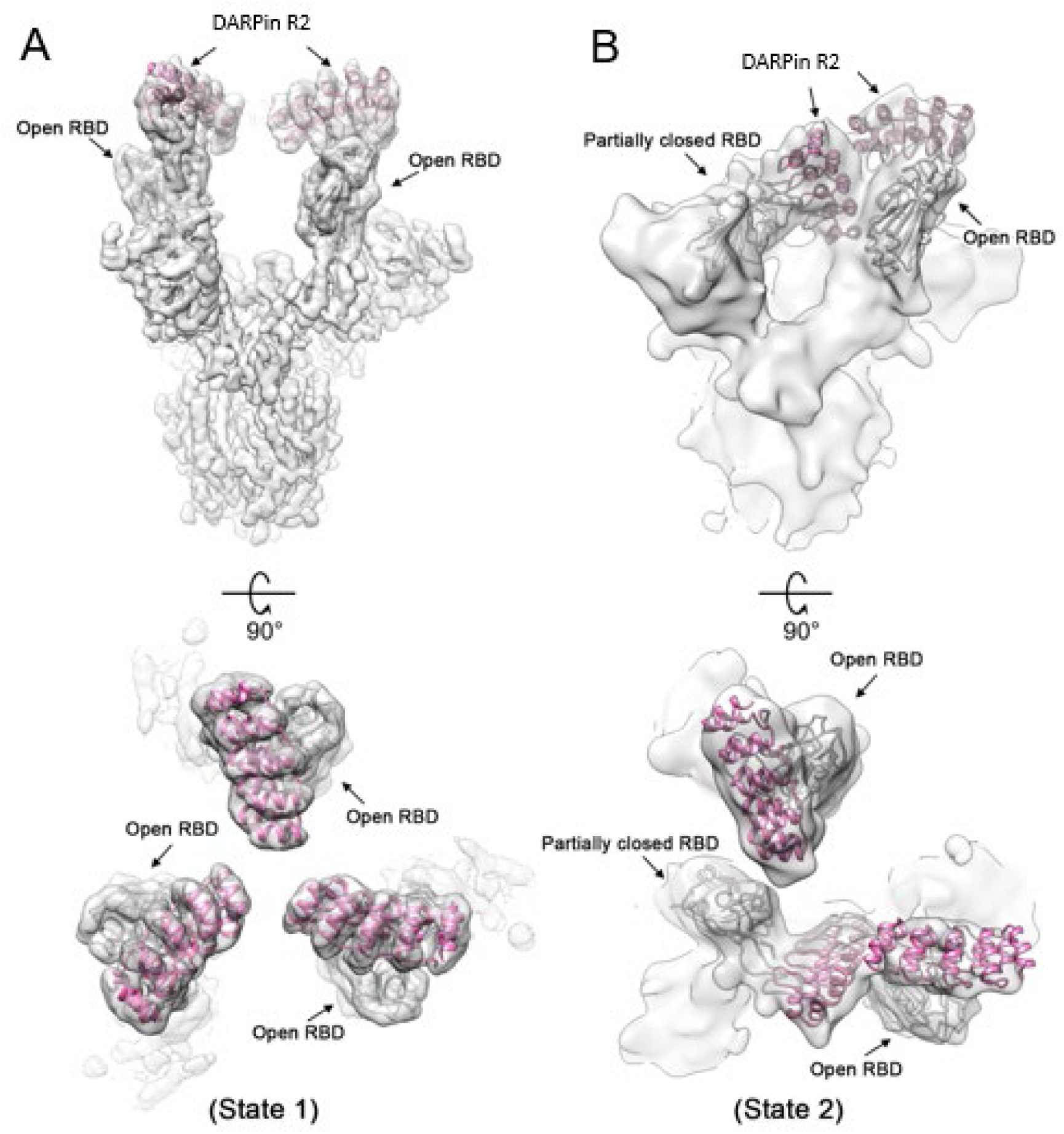
Monovalent DARPin R2 prevents full closure of the RBD. A) Cryo-EM density for state 1 and B) state 2 of the SARS-CoV-2 spike ectodomain in complex with the RBD targeting monovalent DARPin R2, shown as two orthogonal views. The pseudo-atomic model of monovalent DARPin R2 in complex with RBD, derived from molecular docking experiments, is fitted in each of the spike protomers and colored grey and pink, respectively.

**Supplementary Figure 4:**
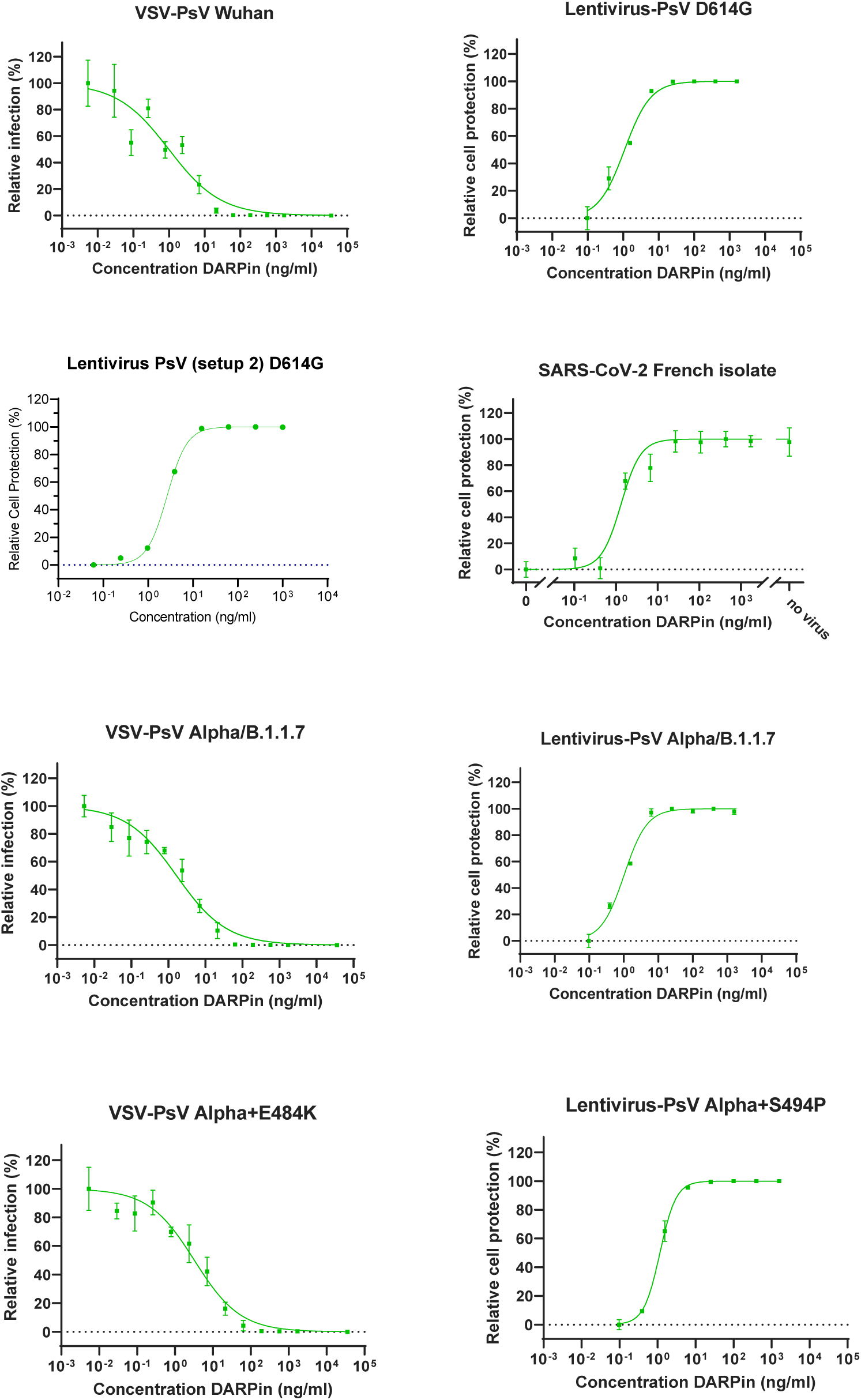
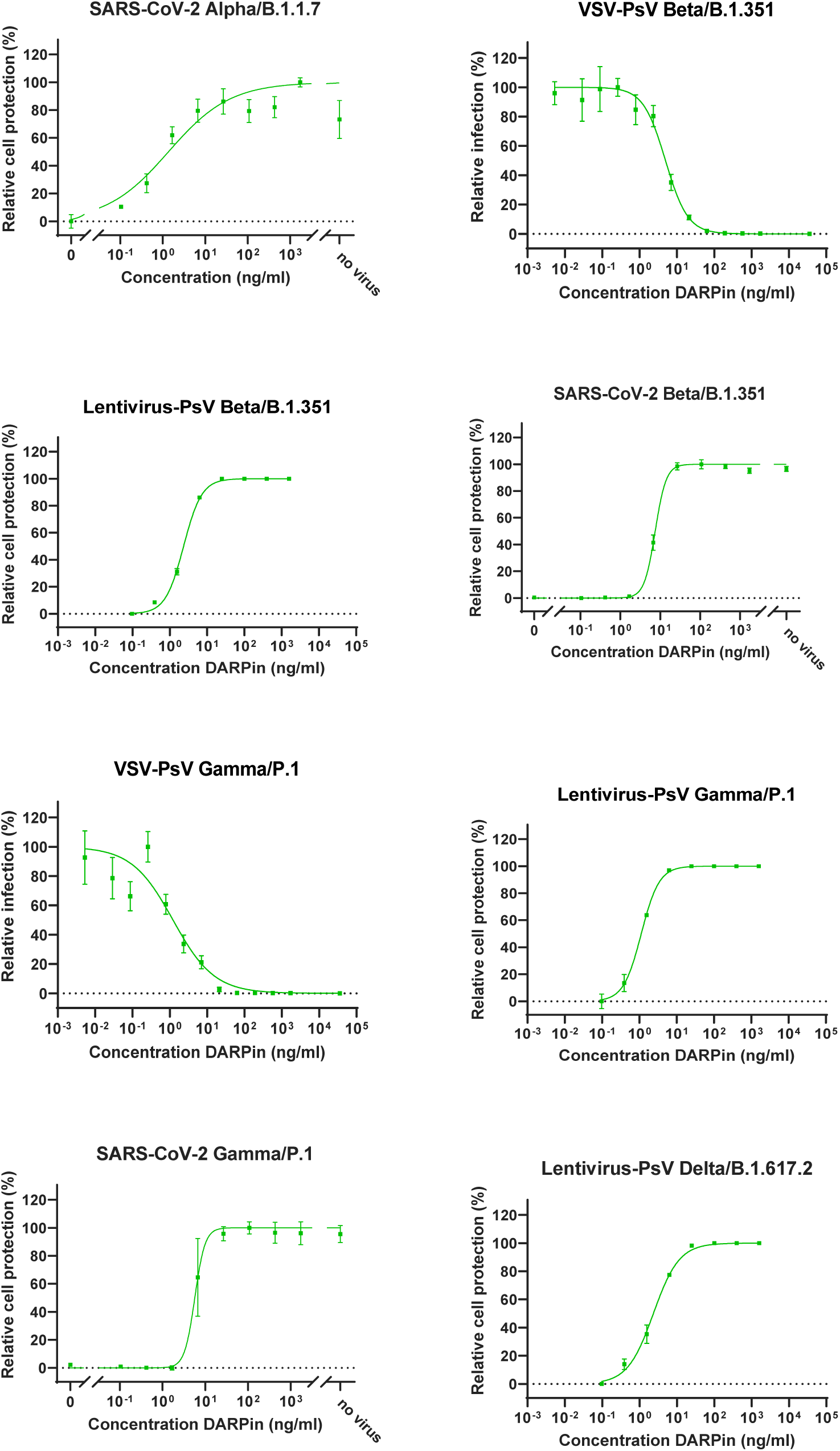
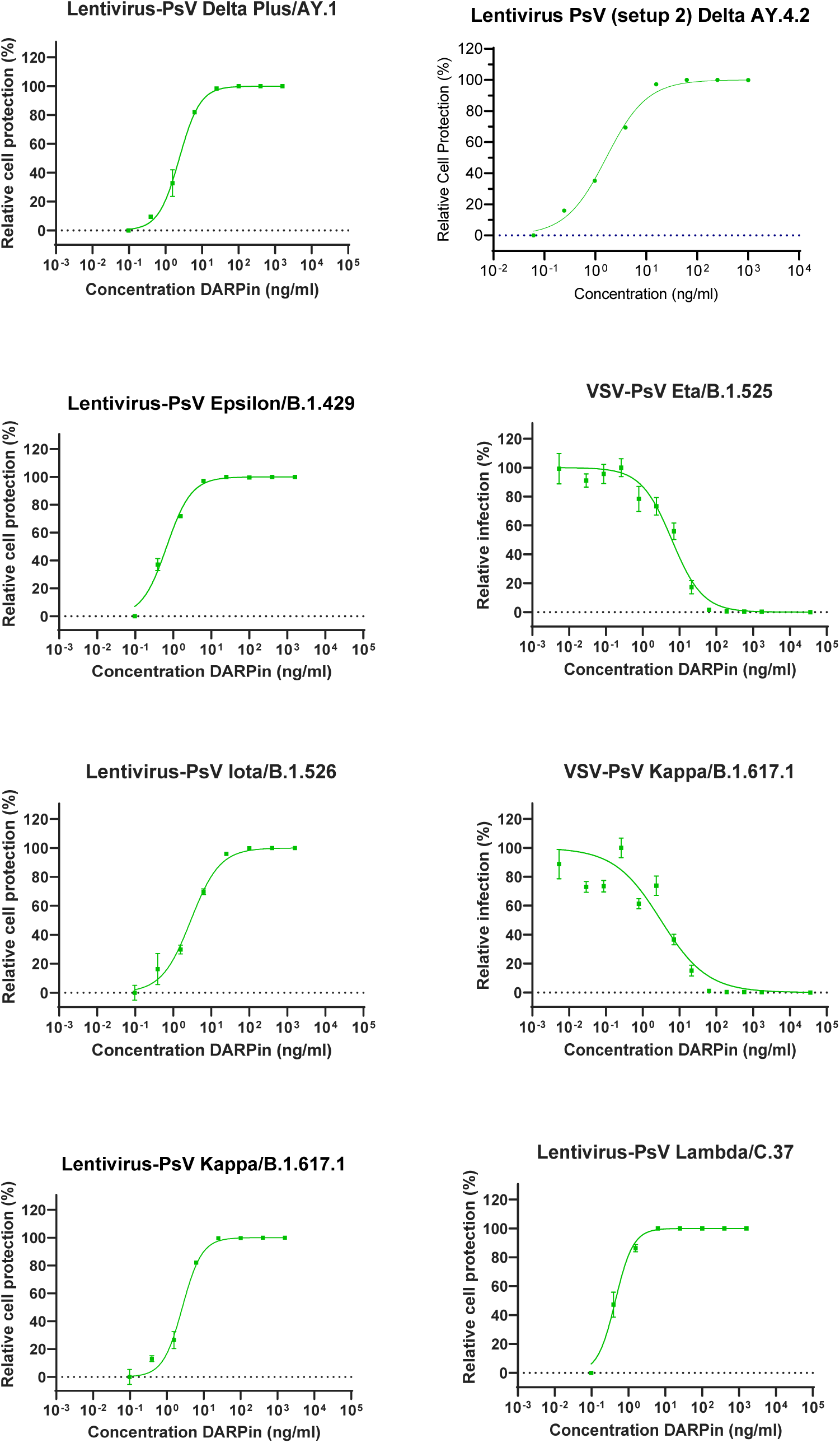
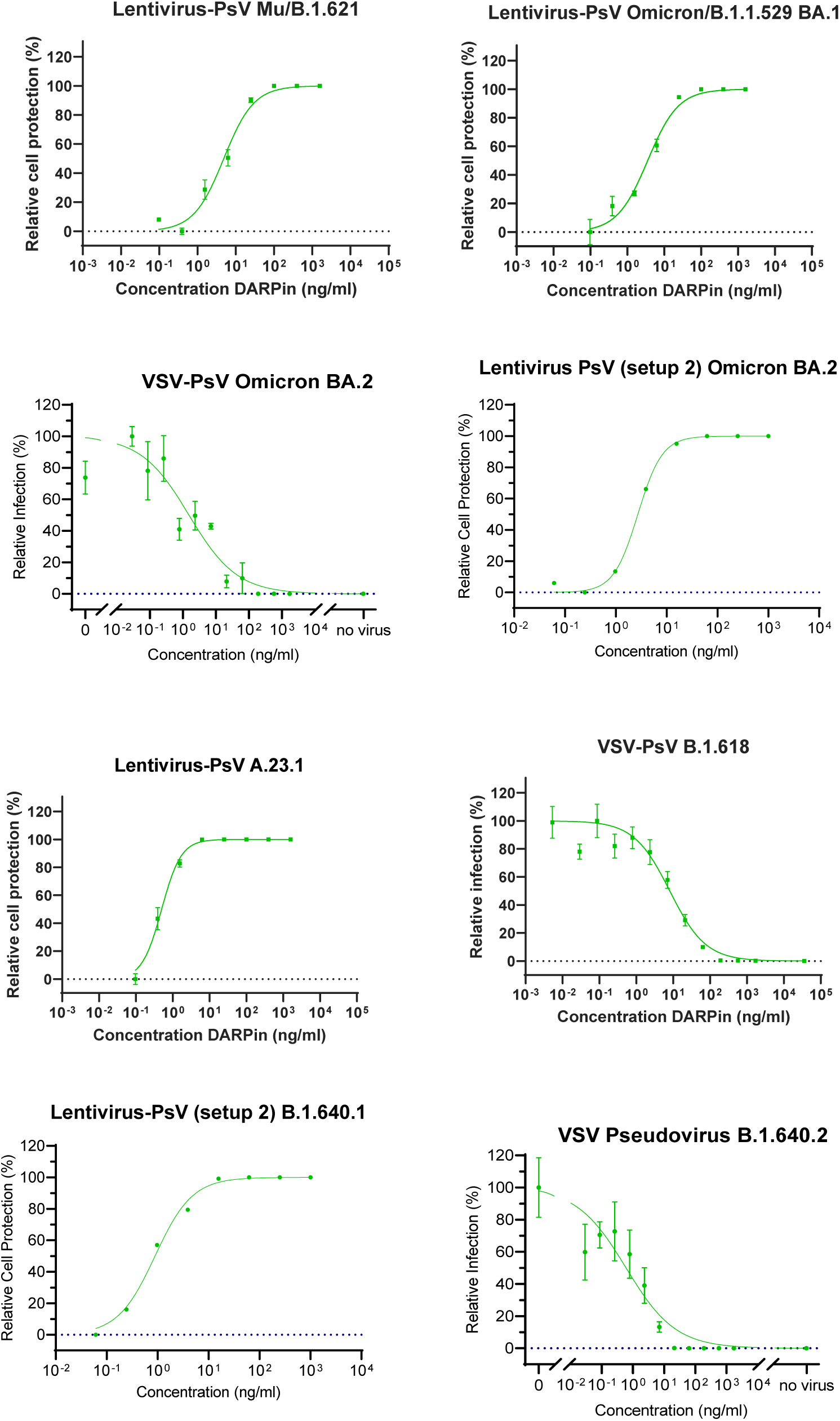
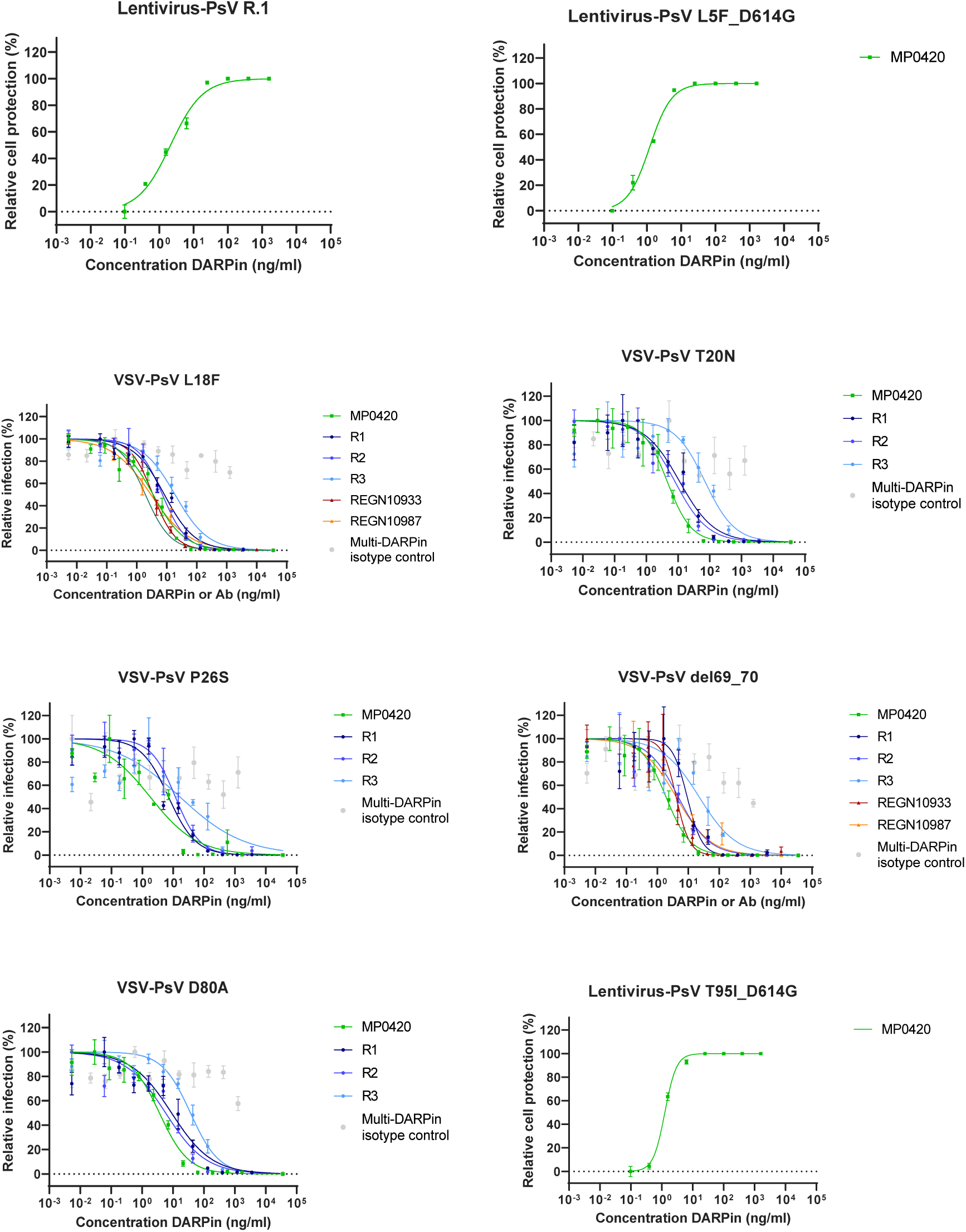
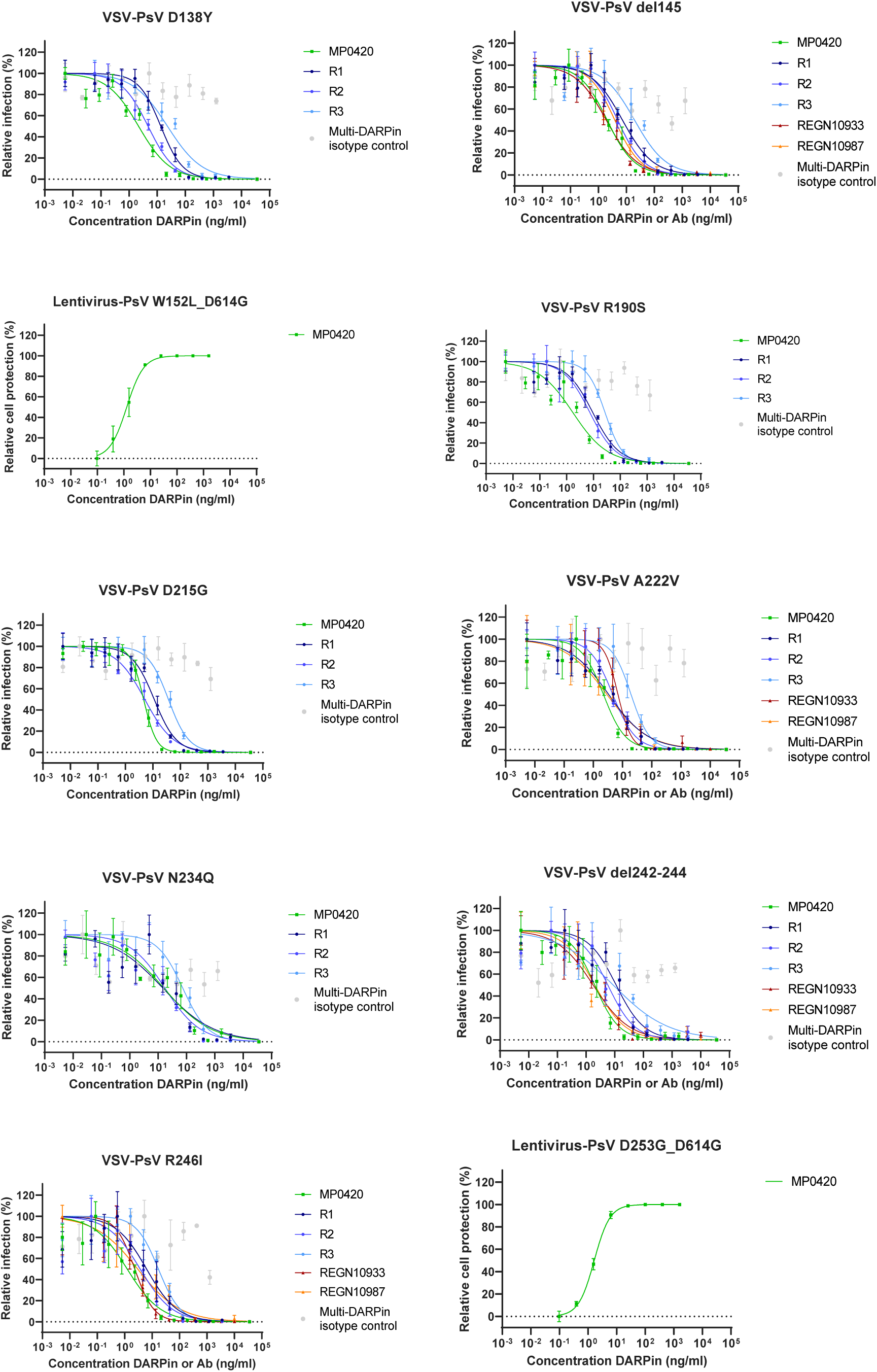
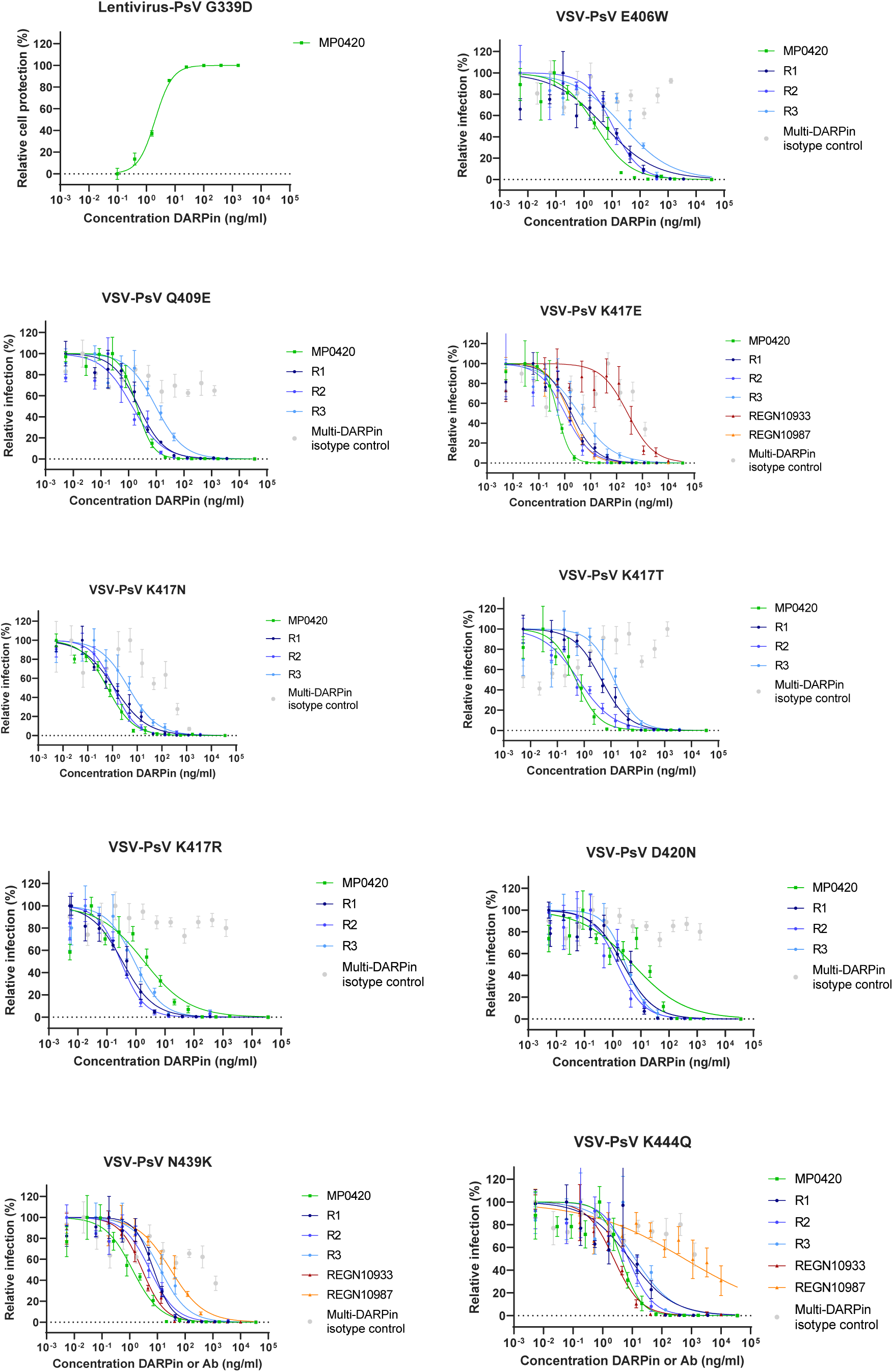
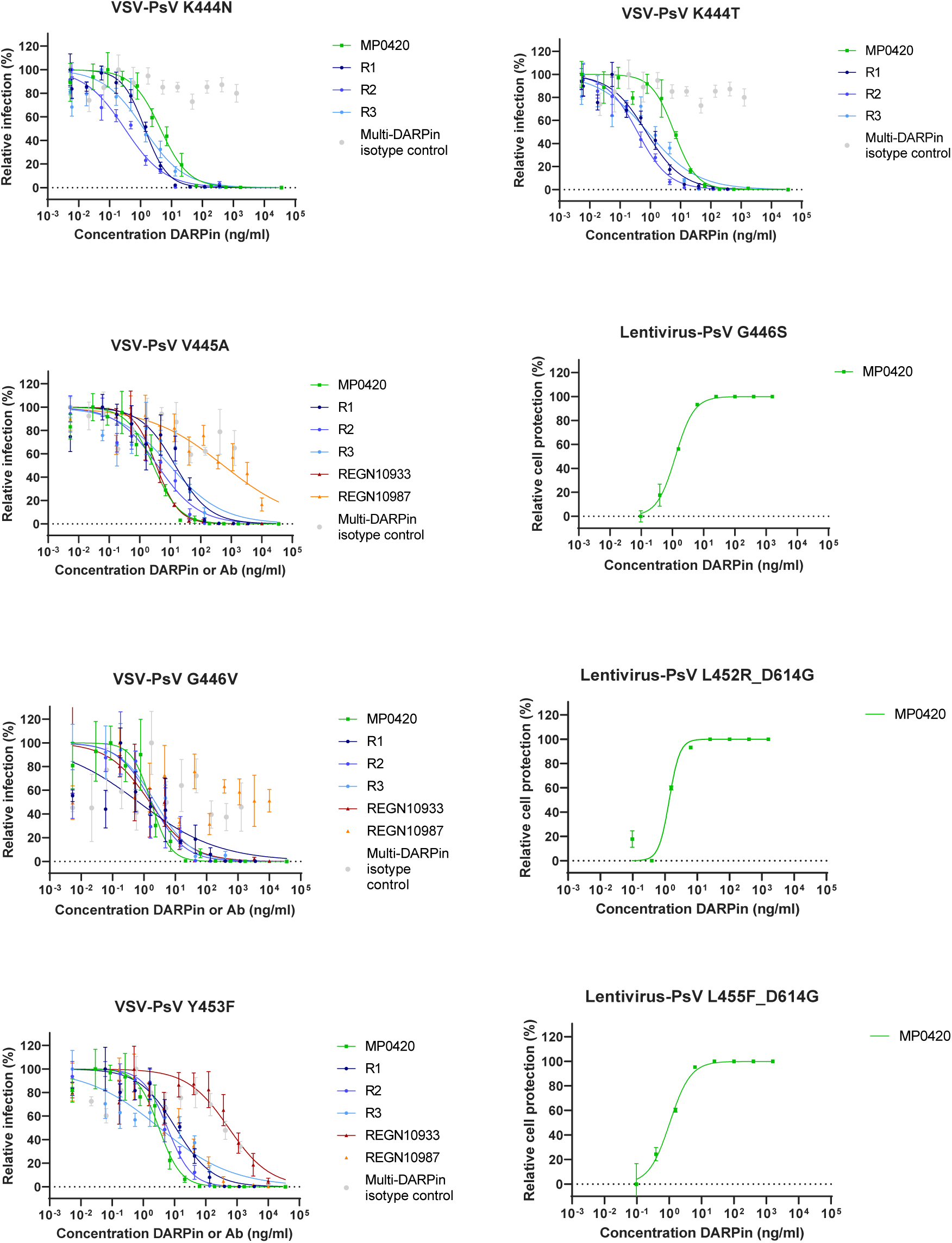
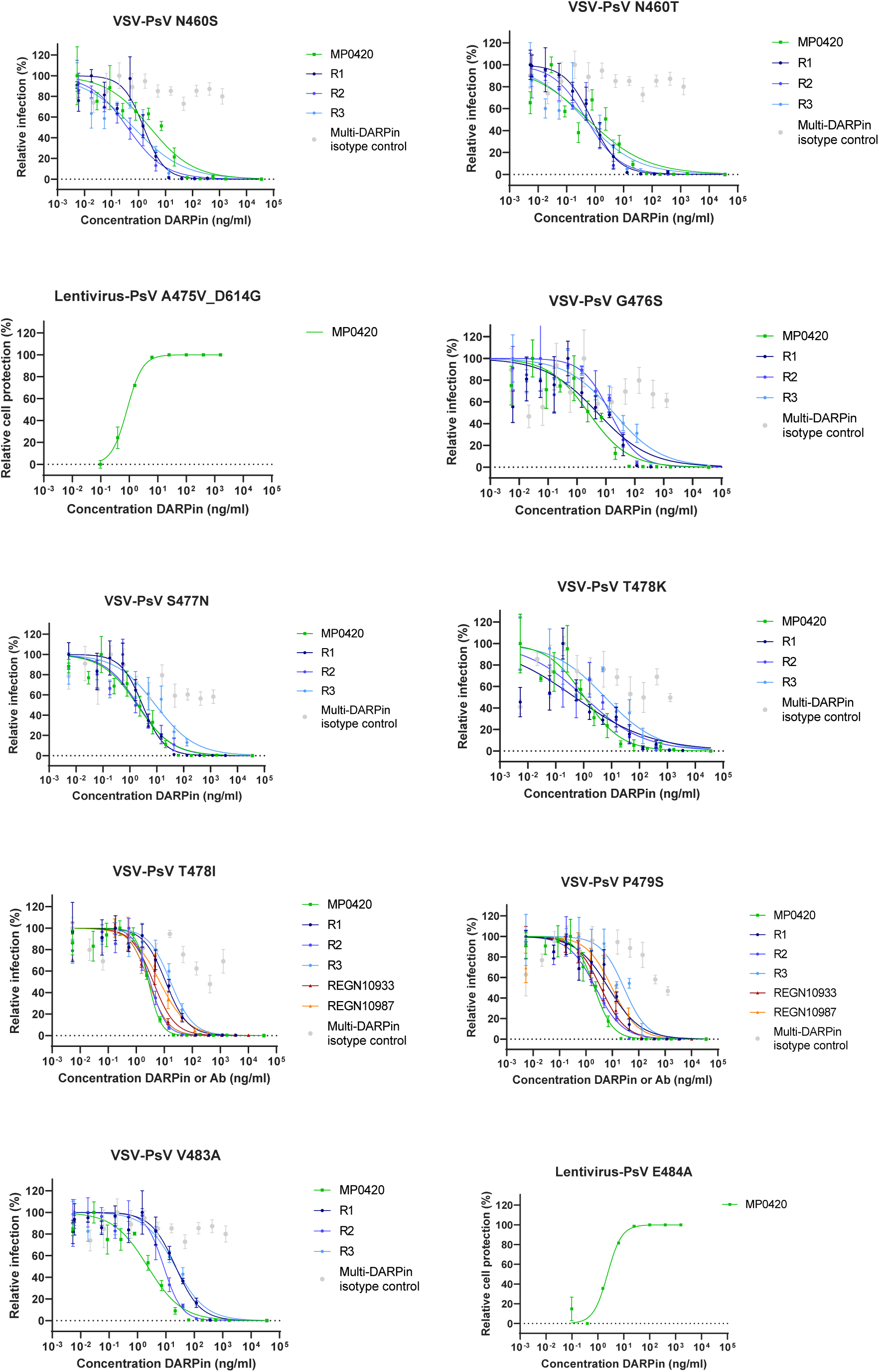
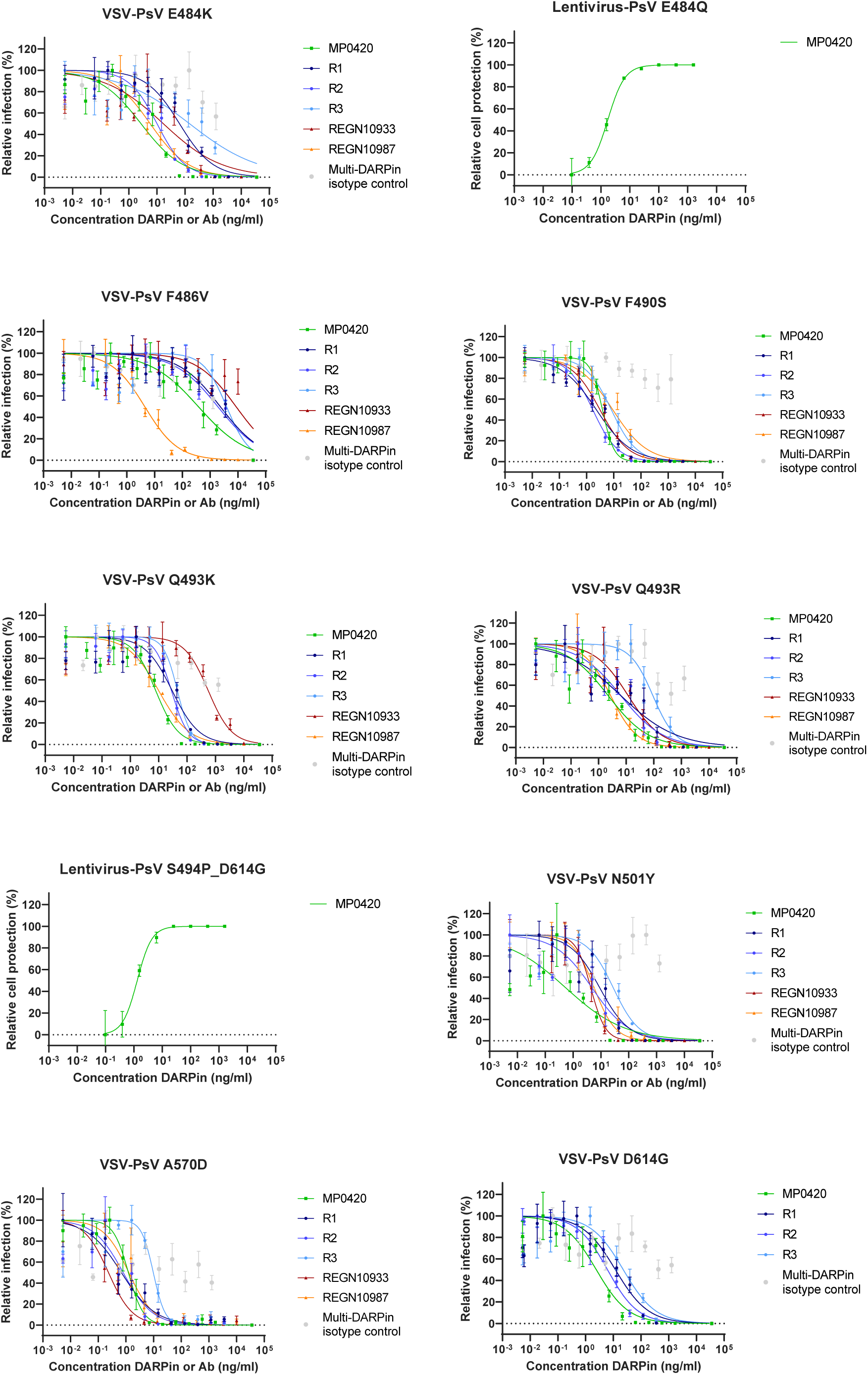
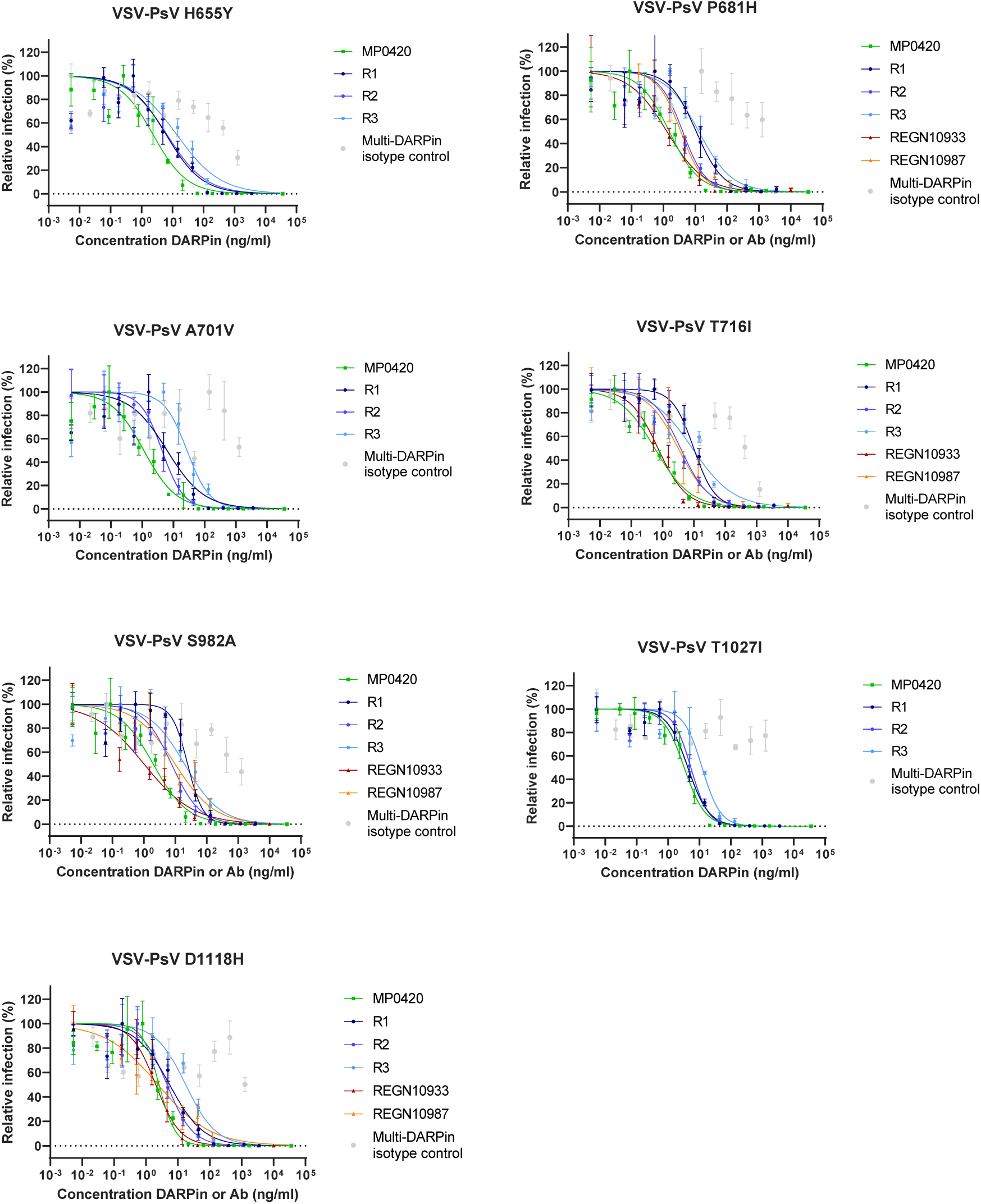
Titration curves for ensovibep (MP0420) and its RBD-binding domains (i.e. R1, R2 and R3), REGN10933 and REGN10987 to determine IC_50_ neutralization potencies on multiple spike mutants or only for ensovibep (MP0420) on the variants, which are summarized in Figure 2, Table 2 and Table 3. Reported is the mean +/− SEM (standard error of the mean).

**Supplementary Figure 5:**
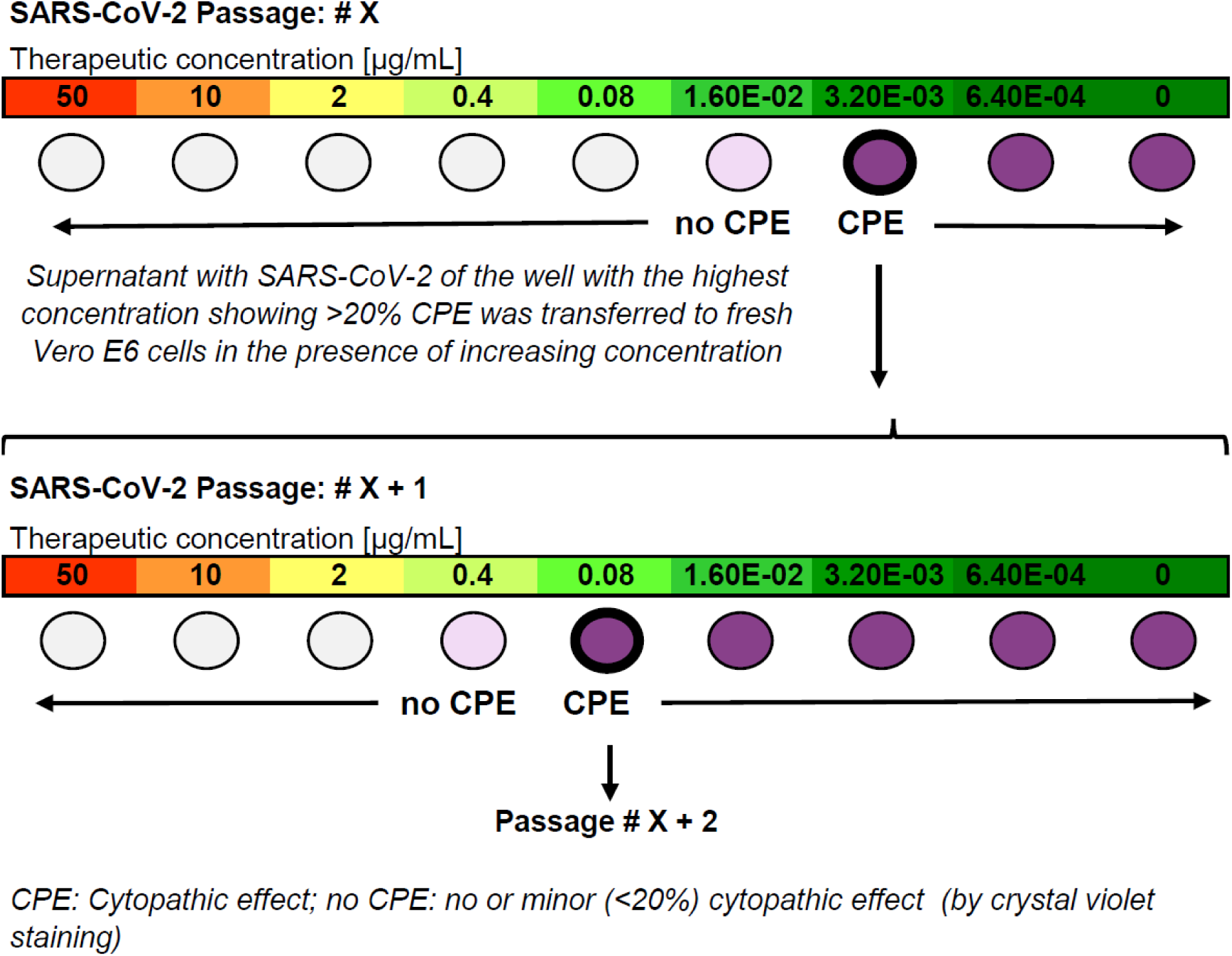
Overview of the experimental protocol for viral passaging: A patient SARS-CoV-2 isolate from early 2020 (1.5 ×10^6^ pfu) was incubated in presence of increasing concentrations of DARPin candidate or antibody for 4 days on Vero E6 cells and virus-induced cytopathic effects (CPE) were determined by microscopy. For each DARPin and antibody condition, cultures showing significant cytopathic effect (≥20%) under the greatest selective pressure were selected and virus-containing supernatant collected to start a new culture passage on Vero E6 cells (bold circle), again under increasing concentrations of the corresponding DARPin candidate or antibody condition. Passaging of virus containing supernatant was continued in the same manner for a total of 4 passages.

**Supplementary Figure 6:**
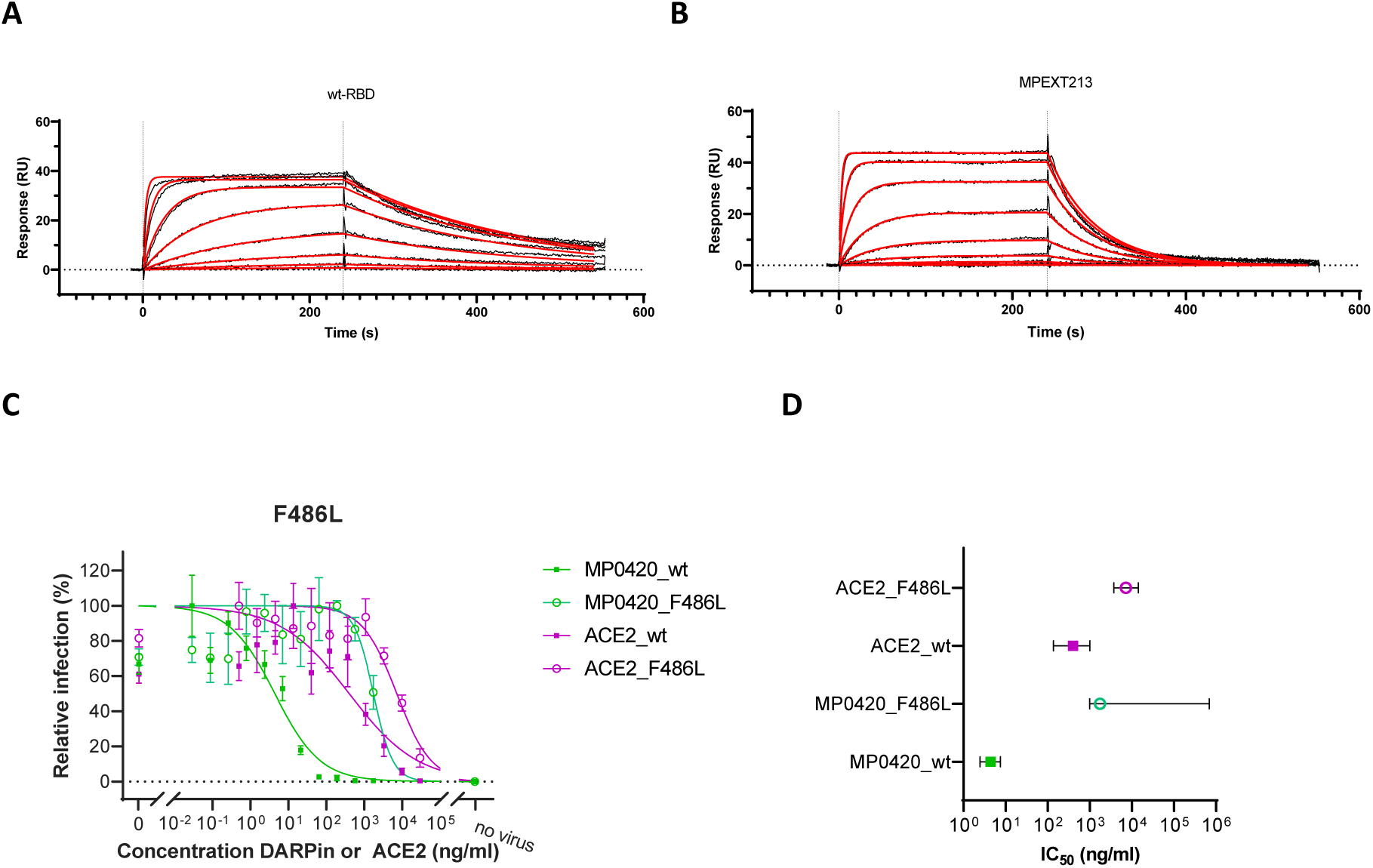
Impact of mutation F486V/L in the RBD on ACE2 binding and neutralization potency of ACE2 or ensovibep (MP0420) in a pseudotype assay. A, B) Binding kinetics for different concentrations of ACE2 was determined by SPR (surface plasmon resonance) with A) immobilized wild type RBD and B) immobilized RBD with the substitution F486V (MPEXT213). Consequently, a drop in affinity was observed upon tested substitution from a KD of 7.8 nM for wild type (k_on_: 6.0E+05; k_off_: 4.8E-03) to a KD of 68.1 nM for F486V (k_on_: 2.7E+05; k_off_: 1.8E-02) C, D) Titration of ACE2 and ensovibep (MP0420) for neutralization of a VSV pseudotype with SARS-CoV-2 wild type spike protein compared to F486L substituted in the spike protein. D) IC_50_ values with 95% confidential interval for the titrations shown in C) demonstrating the loss of potency for ACE2 and ensovibep due to the F486L substitution. In accordance with the SPR measurement, a >10-fold drop in neutralization potency was observed in a VSV pseudotype assay, when ACE2 was used as a competitor. In relation, a >100-fold drop in potency was observed for ensovibep based on the F486L substitution. Reported is the mean +/− SEM (standard error of the mean). Shown experiments further underlines the reduction in binding of ACE2 to F486 substitutions and the importance of F486 for the SARS-CoV-2 virus to maintain the interaction with the human ACE2 receptor. So far, based on the global SARS-CoV-2 database sequences published in GISAID, mutations in position F486 (the core RBD-interaction residue for ensovibep) occur at very low frequencies.

**Supplementary Figure 7:**
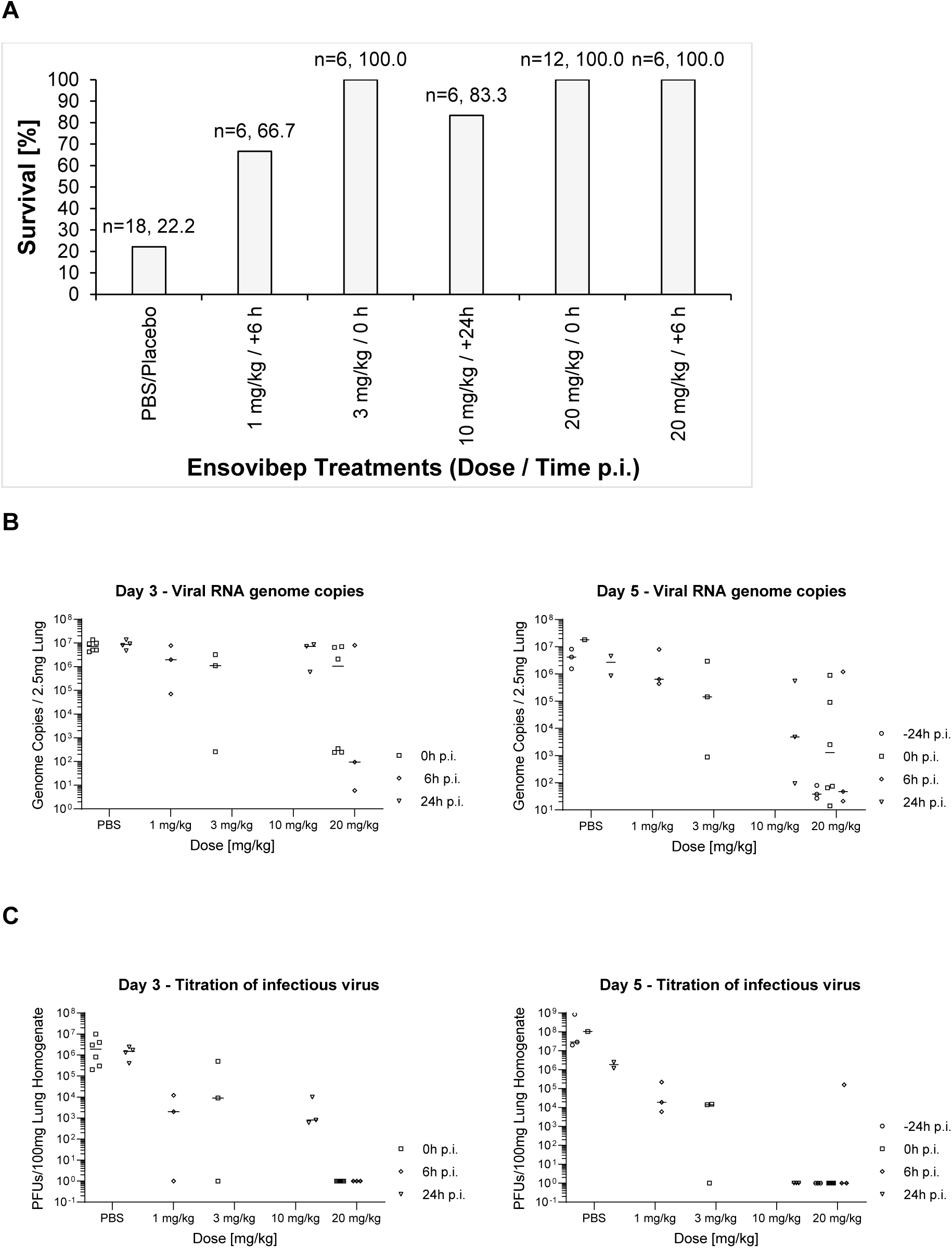
Summarized previous in vivo studies with Roborovski dwarf hamster infected with WT SARS-CoV-2 and treated with ensovibep at various doses and administration time points. A) Animal survival, end-point analysis, animals that had to be euthanized according to score sheet criteria were considered non-survived, animals that reached their respective defined take-out at day 3 or 5 post infection were considered survived. B) qPCR analysis of virus gRNA copy numbers in oropharyngeal swabs and lung homogenates at day 3 or day 5 post infection C) Titration of replication competent virus from lung homogenates as plaque assay on Vero E6 cells at day 3 or day 5 post infection. Reported are the values of the individual animals and the median.

**Supplementary Figure 8:**
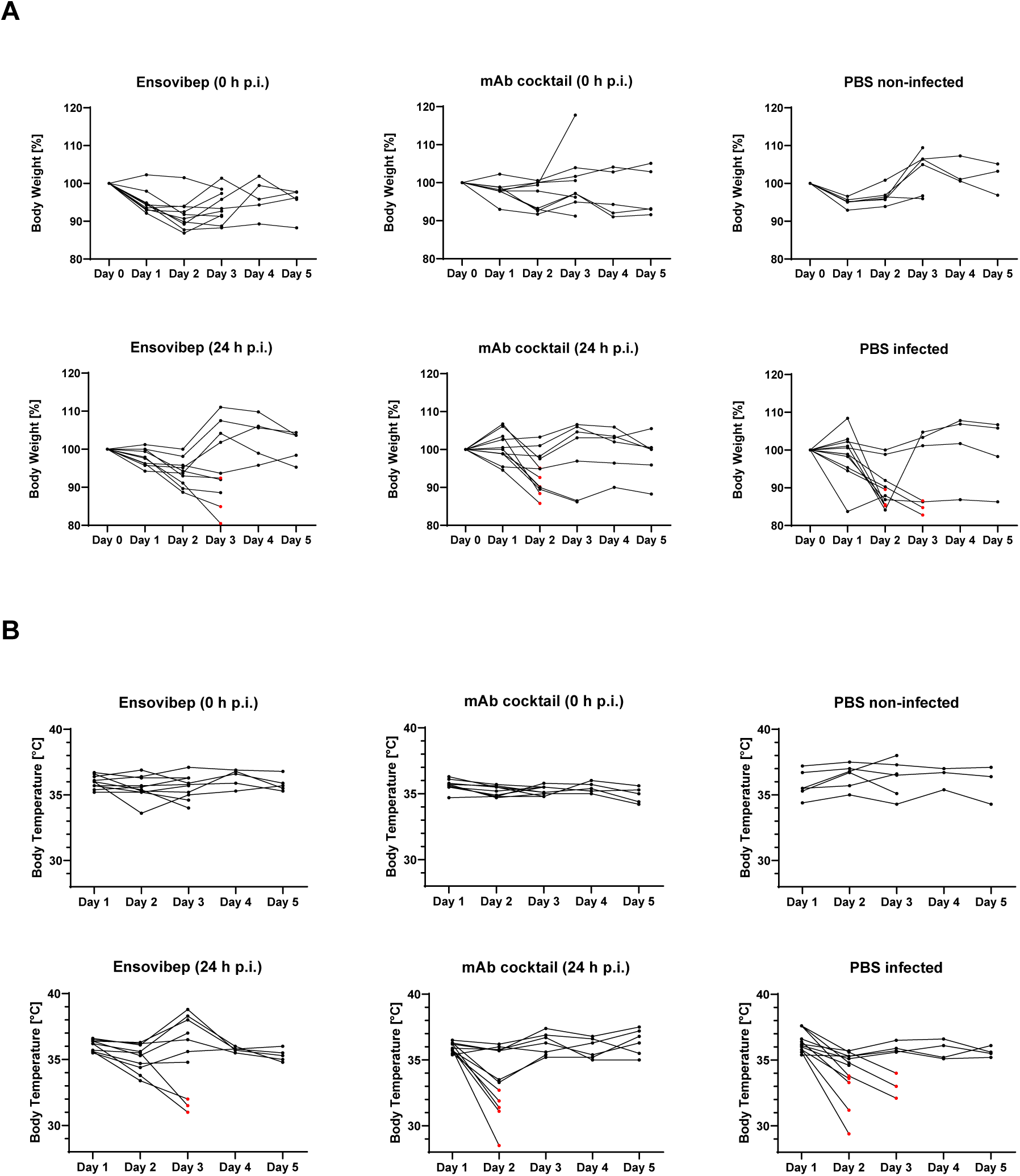
Clinical Parameters of individuals over the course of infection, (mean +/- SD presented in Figure 5C) A) Body weight changes of individual hamsters B) Body temperatures of individual hamsters. Animals that had to be euthanized based on score sheet criteria are marked in red.

**Supplementary Figure 9:**
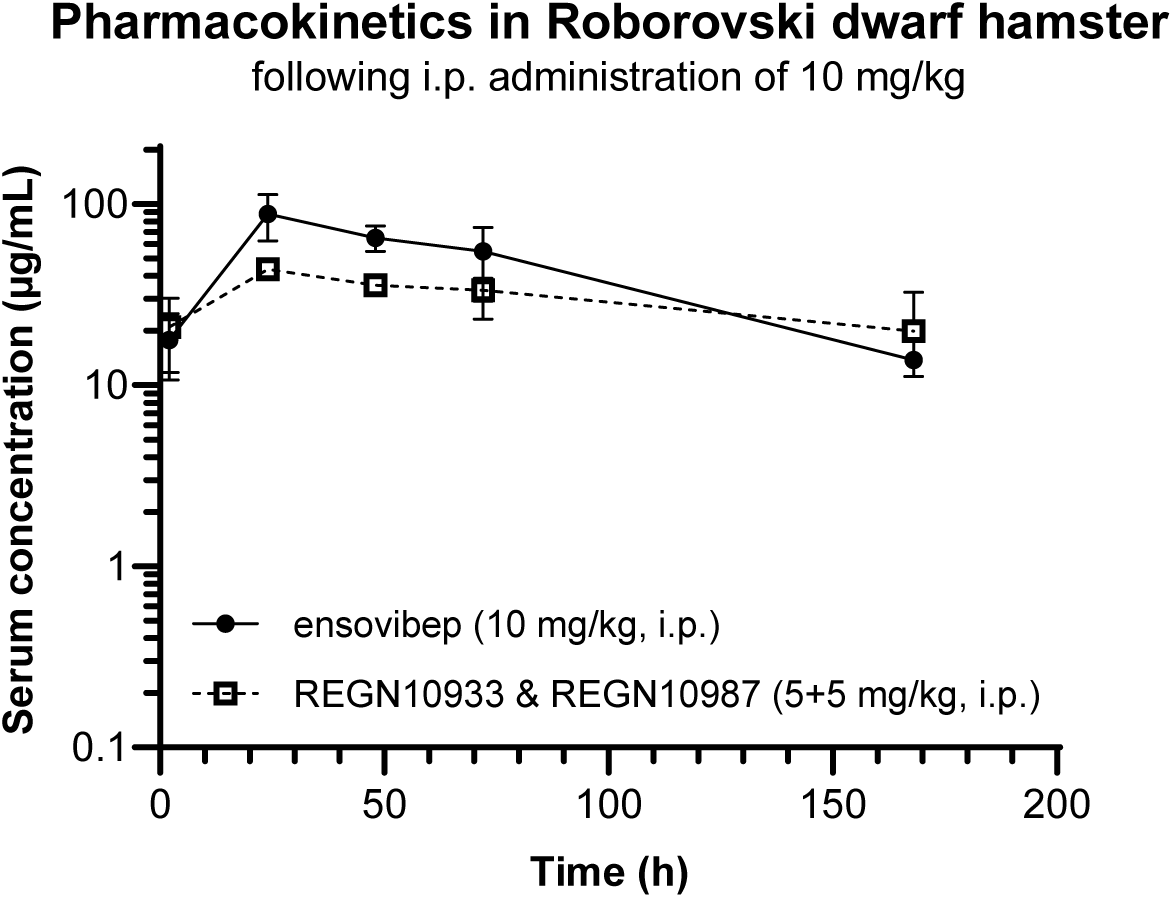
Pharmacokinetics profiles of non-infected Roborovski dwarf hamsters injected i.p. with either 10 mg/kg of ensovibep or the cocktail of REGN10933 and REGN10987 at 5 mg/kg for each of the monoclonal antibodies. Three animals were sacrificed for determination of the therapeutic concentration in the serum of the terminal bleeds. Obvious outliers due to likely a failure of the intraperitoneal injection were removed from the evaluation. Reported is the mean +/− SEM (standard error of the mean). Pharmacokinetic parameters for ensovibep: T_1/2_: 52.0 h; C_max_: 87.8 μg/mL; T_max_: 24 h. Pharmacokinetic parameters for the cocktail of REGN10933 and REGN10987: T_1/2_: 139 h; C_max_: 43.5 μg/mL; T_max_: 24h.

**Supplementary Table 1:**
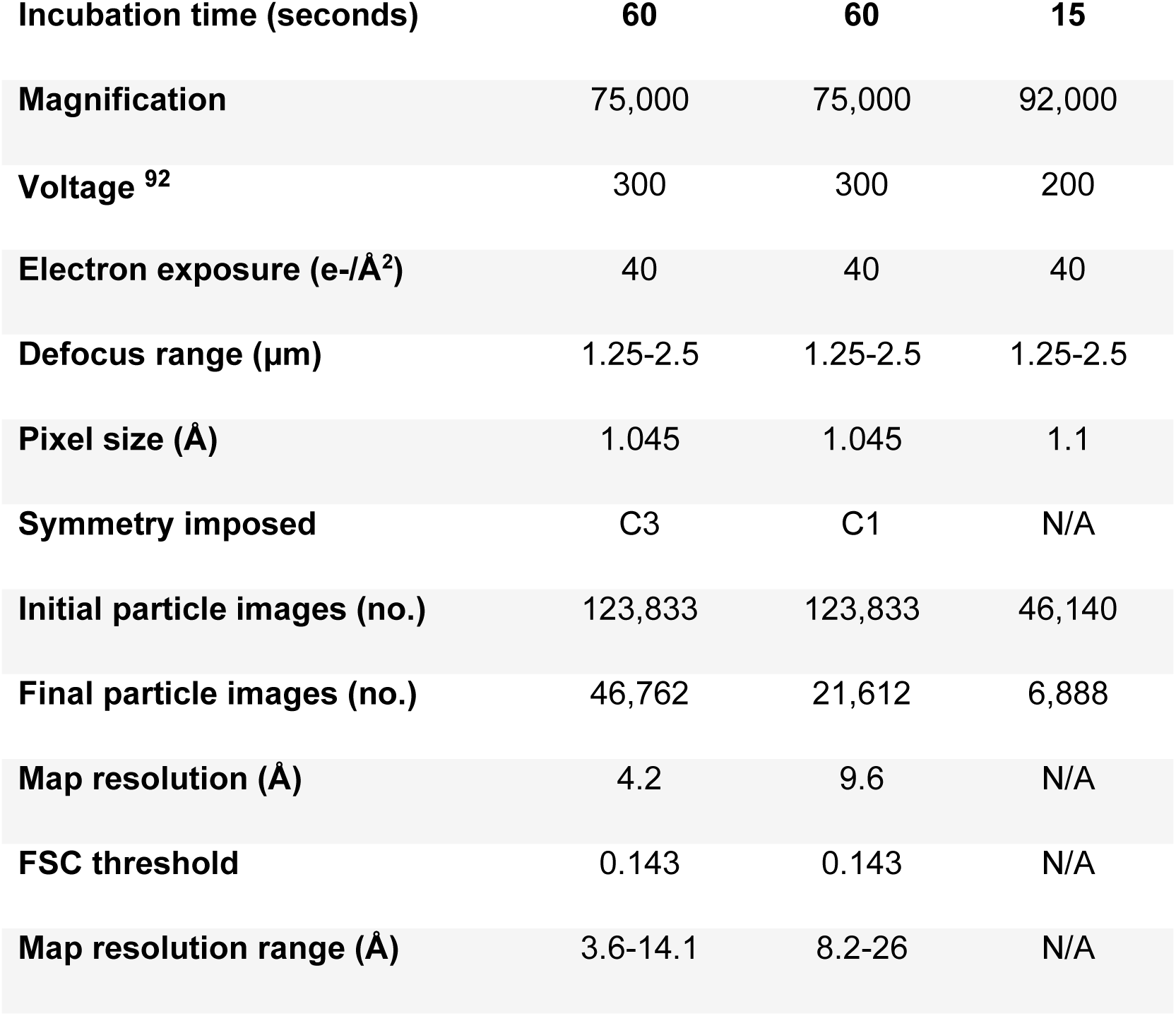
Cryo-EM data collection and image processing information.

| **Incubation time (seconds)** | **60** | **60** | **15** |
| --- | --- | --- | --- |
| **Magnification** | 75,000 | 75,000 | 92,000 |
| **Voltage <sup>92</sup>** | 300 | 300 | 200 |
| **Electron exposure (e-/Å<sup>2</sup>)** | 40 | 40 | 40 |
| **Defocus range (µm)** | 1.25-2.5 | 1.25-2.5 | 1.25-2.5 |
| **Pixel size (Å)** | 1.045 | 1.045 | 1.1 |
| **Symmetry imposed** | C3 | C1 | N/A |
| **Initial particle images (no.)** | 123,833 | 123,833 | 46,140 |
| **Final particle images (no.)** | 46,762 | 21,612 | 6,888 |
| **Map resolution (Å)** | 4.2 | 9.6 | N/A |
| **FSC threshold** | 0.143 | 0.143 | N/A |
| **Map resolution range (Å)** | 3.6-14.1 | 8.2-26 | N/A |

**Supplementary Table 2:** In vitro protection against emerging SARS-CoV-2 variants for ensovibep

| Variant | Substitutions / Deletions | Assay Type | Neutralizing IC<sub>50</sub><br>[ng/mL] |
| --- | --- | --- | --- |
| **References** | Wuhan wild type | VSV pseudotype | 1 |
|  | D614G background | Lentivirus pseudotype | 1.1 |
|  | D614G background | Lentivirus pseudotype<br>Setup 2 | 2.7 |
|  | French isolate: V367F; E990A | Authentic virus | 1.3 |
| **Alpha / B.1.1.7** | 69-70 del, del145, N501Y, A570D, D614G, P681H, T716I, S982A, D1118H | VSV | 1.7 |
|  | 69-70 del, del145, E484K, N501Y, A570D, D614G, P681H, T716I, S982A, D1118H | VSV | 3.2 |
|  | 69-70 del, del145, N501Y, A570D, D614G, P681H, T716I, S982A, D1118H | Lentivirus pseudotype | 0.9 |
|  | 69-70 del, del145, S494P, N501Y, A570D, D614G, P681H, T716I, S982A, D1118H | Lentivirus pseudotype | 0.8 |
|  | H69_V70del, Y145del, N501Y, A570D, D614G, P681H, T716I, S982A, D1118H | Authentic | 1.3 |
| **Beta / B.1.351** | D80A, D215G, E484K, N501Y, A701V | VSV | 5.5 |
|  | L18F, D80A, D215G, Del242-244, R246I, K417N, E484K, N501Y, D614G, A701V | VSV | 5 |
|  | L18F, D80A, D215G, Del242-244, K417N, E484K, N501Y, D614G, A701V | Lentivirus pseudotype | 1.2 |
|  | L18F, D80A, D215G, L242_L244del, T302T, K417N, E484K, N501Y, D571D, D614G, A701V | Authentic | 7.5 |
| **Gamma / P.1** | L18F, T20N, P26S, D138Y, R190S, K417T, E484K, N501Y, D614G, H655Y, T1027I | VSV | 1.2 |
|  | L18F, T20N, P26S, D138Y, R190S, K417T, E484K, N501Y, D614G, H655Y, T1027I, V1176F | Lentivirus pseudotype | 0.7 |
|  | L18F, T20N, P26S, D138Y, R190S, K417T, E484K, N501Y, D614G, H655Y, T1027I, V1176F | Authentic | 5.7 |
| **Delta/DeltaPlus / B.1.617.2** | T19R, G142D, E156G, F157del, R158del, L452R, T478K, D614G, P681R, D950N | Lentivirus pseudotype | 2.4 |
|  | T19R, T95I, G142D, E156G, F157del, R158del, W258L, K417N, L452R, T478K, D614G, P681R, D950N | Lentivirus pseudotype | 2.6 |
|  | T19R, T95I, G142D, Y145H, E156G, F157-, R158-, A222V, L452R, T478K, D614G, P681R, D950N | Lentivirus pseudotype<br>(Setup 2) | 1.6 |
| **Epsilon/B.1.429** | S13I, W152C, L452R, D614G | Lentivirus pseudotype | 0.9 |
|  | S13I, P26S, W152C, L452R, D614G | Lentivirus pseudotype | 0.5 |
| **Iota/B.1.526** | L5F, T95I, D253G, E484K, D614G, A701V | Lentivirus pseudotype | 3.0 |
| **Kappa / B.1.617.1** | T95I, G142D, E154K, L452R, E484Q, D614G, P681R, Q1071H | Lentivirus pseudotype | 2.0 |
|  | G142D, E154K, V382L, L452R, E484Q, D614G, P681R, Q1071H, D1153Y | Lentivirus pseudotype | 1.9 |
| **Lambda / C37** | G75V, T76I, R246del, S247-G252del, D253N, L452Q, F490S, D614G, T859N | Lentivirus pseudotype | 0.4 |
| **Mu / B.1.621** | T95I, Y144S, Y145N, R346K, E484K, N501Y, D614G, P681H, D950N | Lentivirus pseudotype | 6.1 |
| **Omicron / B.1.1.529 / BA.1** | A67V, Δ69-70, T95I, G142D, Δ143-145, Δ211, L212I, ins214EPE, G339D, S371L, S373P, S375F, K417N, N440K, G446S, S477N, T478K, E484A, Q493R, G496S, Q498R, N501Y, Y505H, T547K, D614G, H655Y, N679K, P681H, N764K, D796Y, N856K, N969K, L981F | VSV pseudotype | 2.2 |
|  | A67V, Δ69-70, T95I, G142D, Δ143-145, Δ211, L212I, ins214EPE, G339D, S371L, S373P, S375F, K417N, N440K, G446S, S477N, T478K, E484A, Q493K, G496S, Q498R, N501Y, Y505H, T547K, D614G, H655Y, N679K, P681H, N764K, D796Y, N856K, Q954H, N969K, L981F | VSV pseudotype | 2.1 |
|  | A67V, Δ69-70, T95I, G142D, Δ143-145, Δ211, L212I, ins214EPE, G339D, S371L, S373P, S375F, K417N, N440K, G446S, S477N, T478K, E484A, Q493R, G496S, Q498R, N501Y, Y505H, T547K, D614G, H655Y, N679K, P681H, N764K, D796Y, N856K, Q954H, N969K, L981F | Lentivirus pseudotype | 3.6 |
| **Omicron / B.1.1.529 / BA.2** | T19I, L24-, P25-, P26-, A27S, G142D, V213G, G339D, S371F, S373P, S375F, T376A, D405N, R408S, K417N, N440K, S477N, T478K, E484A, Q493R, Q498R, N501Y, Y505H, D614G, H655Y, N679K, P681H, N764K, D796Y, Q954H, N969K | Lentivirus pseudotype (Setup 2) | 2.7 |
|  | T19I, L24-, P25-, P26-, A27S, G142D, V213G, G339D, S371F, S373P, S375F, T376A, D405N, R408S, K417N, N440K, S477N, T478K, E484A, Q493R, Q498R, N501Y, Y505H, D614G, H655Y, N679K, P681H, N764K, D796Y, Q954H, N969K | VSV pseudotype | 1.5 |
| **B.1.640.1** | P9L_E96Q_C136del_N137del_D138del_P139del_F140del_L141del_G142del_V143del_Y144del_R190S_I210T_R346S_N394S_Y449N_F490R_N501Y_D614G_P681H_T859N_D936H | Lentivirus pseudotype (Setup 2) | 0.9 |
| **B.1.640.2** | P9L, E96Q, delC136-Y144, R190S, D215H, R346S, N394S, Y449N, E484K, F490S, N501Y, D614G, P681H, T859N, D1139H | VSV pseudotype | 0.6 |
| **R.1** | W152L, E484K, D624G, G769V | Lentivirus pseudotype | 2.4 |
| **A.23.1** | F157L, V367F, Q613H, D614G, P681R | Lentivirus pseudotype | 0.3 |

**Supplementary Table 3:** *In vitro* protection against SARS-CoV-2 spike protein substitutions or deletions for ensovibep.

| Amino acid position | Substitution / deletion | Assay Type | Neutralizing IC<sub>50</sub> [ng/mL] |
| --- | --- | --- | --- |
| L18 | F | VSV | 3.5 |
| T20 | N | VSV | 4.6 |
| P26 | S | VSV | 1.8 |
| 69-70 | del | VSV | 1.9 |
| D80 | A | VSV | 3.6 |
| T95 | I | Lenti | 0.9 |
| D138 | Y | VSV | 2.2 |
| 145 | del | VSV | 2.1 |
| W152 | L | Lenti | 1.2 |
| R190 | S | VSV | 1.7 |
| A222 | V | VSV | 2.2 |
| N234 | Q | VSV | 16.2 |
| 242-244 | del | VSV | 2.0 |
| G339 | D | Lenti | 2.0 |
| E406 | Q | Lenti | 1.5 |
|  | W | VSV | 2.7 |
| Q409 | E | VSV | 2.0 |
| K417 | E | VSV | 0.5 |
|  | N | VSV | 0.6 |
|  | R | VSV | 2.1 |
|  | T | VSV | 0.5 |
| D420 | N | VSV | 5.6 |
| N439 | K | VSV | 1.3 |
| K444 | E | Lenti | 0.8 |
|  | N | VSV | 4.4 |
|  | Q | Lenti | 1.3 |
|  | T | VSV | 6.1 |
| V445 | A | Lenti | 1.3 |
| G446 | V | VSV | 1.7 |
|  | S | Lenti | 1.3 |
| N450 | D | Lenti | 0.9 |
| L452 | R | Lenti | 0.4 |
| Y453 | F | VSV | 3.2 |
| L455 | F | Lenti | 1.1 |
| N460 | S | VSV | 2.6 |
|  | T | VSV | 0.6 |
| A475 | V | Lenti | 0.9 |
| G476 | S | VSV | 1.5 |
| S477 | N | VSV | 1.9 |
| T478 | I | VSV | 2.7 |
|  | K | Lenti | 1.5 |
| P479 | S | VSV | 2.1 |
| V483 | A | VSV | 2.3 |
| E484 | A | Lenti | 2.4 |
|  | K | VSV | 2.7 |
|  | Q | Lenti | 2.3 |
| G485 | D | VSV | 28.5 |
| F486 | V | VSV | >100 |
|  | L | VSV | >100 |
| F490 | S | VSV | 3.8 |
| Q493 | K | VSV | 7.9 |
|  | R | VSV | 2.2 |
| S494 | P | Lenti | 1.3 |
| N501 | Y | VSV | 0.6 |
| A570 | D | VSV | 1.2 |
| D614 | G | VSV | 2.4 |
| H655 | Y | VSV | 2.4 |
| P681 | H | VSV | 1.5 |
| A701 | V | VSV | 1.1 |
| T716 | I | VSV | 0.6 |
| S982 | A | VSV | 2.0 |
| T1027 | I | VSV | 3.3 |
| D1118 | H | VSV | 2.6 |

**Supplementary Table 4:** drug exposure levels in serum at day of euthanization. Animals with drug exposure levels below 10% of the group average were removed from the study analysis (depicted in bold).

| Ensovibep<br>(10 mg/kg i.p.; 0 h p.i) |  |  | Ensovibep<br>(10 mg/kg i.p.; 24 h p.i) |  |  |
| --- | --- | --- | --- | --- | --- |
| # | Animal ID | Serum concentration<br>[µg/mL] | # | Animal ID | Serum concentration<br>[µg/mL] |
| 1 | DN5_1 | 86.9 | 1 | DN5_25 | 17.0 |
| 2 | **DN5_2** | **2.7** | 2 | **DN5_26** | **1.8** |
| 3 | DN5_3 | 169.7 | 3 | DN5_27 | 39.1 |
| 4 | DN5_4 | 44.5 | 4 | DN5_28 | N/A |
| 5 | DN5_5 | 92.9 | 5 | DN5_29 | 48.0 |
| 6 | DN5_6 | 39.2 | 6 | **DN5_30** | **5.7** |
| 7 | **DN5_7** | **1.5** | 7 | DN5_31 | 225.1 |
| 8 | DN5_8 | 109.4 | 8 | DN5_32 | 27.3 |
| 9 | DN5_9 | 51.8 | 9 | DN5_33 | 66.6 |
| 10 | DN5_10 | 127.9 | 10 | DN5_34 | 109.5 |
| 11 | DN5_11 | 70.2 | 11 | DN5_35 | 128.4 |
| 12 | DN5_12 | 38.6 | 12 | DN5_36 | 78.3 |

| REGN10933 & REGN10987<br>(5 + 5 mg/kg i.p.; 0 h p.i) |  |  | REGN10933 & REGN10987<br>(5 + 5 mg/kg i.p.; 24 h p.i) |  |  |
| --- | --- | --- | --- | --- | --- |
| # | Animal ID | Serum concentration<br>[µg/mL] | # | Animal ID | Serum concentration<br>[µg/mL] |
| 1 | DN5_13 | 29.0 | 1 | DN5_37 | 72.0 |
| 2 | **DN5_14** | **0.3** | 2 | DN5_38 | 60.9 |
| 3 | DN5_15 | 26.5 | 3 | DN5_39 | 42.8 |
| 4 | **DN5_16** | **1.1** | 4 | DN5_40 | 32.6 |
| 5 | DN5_17 | 34.0 | 5 | DN5_41 | 46.7 |
| 6 | **DN5_18** | **1.4** | 6 | DN5_42 | 41.9 |
| 7 | DN5_19 | 41.9 | 7 | DN5_43 | 43.7 |
| 8 | DN5_20 | 30.3 | 8 | DN5_44 | 45.3 |
| 9 | DN5_21 | 43.5 | 9 | DN5_45 | 46.1 |
| 10 | DN5_22 | 38.0 | 10 | DN5_46 | 61.9 |
| 11 | DN5_23 | 28.1 | 11 | DN5_47 | 37.2 |
| 12 | DN5_24 | 32.7 | 12 | DN5_48 | 41.8 |

N/A: Not available due to low amount of serum extracted from terminal bleeds
Bold: animals removed from the study data due to low therapeutic exposure
Red: animals euthanized at 2 dpi
Blue: animals euthanized at 3 dpi
Green: animals euthanized at 5dpi

**Supplementary Table 5:**
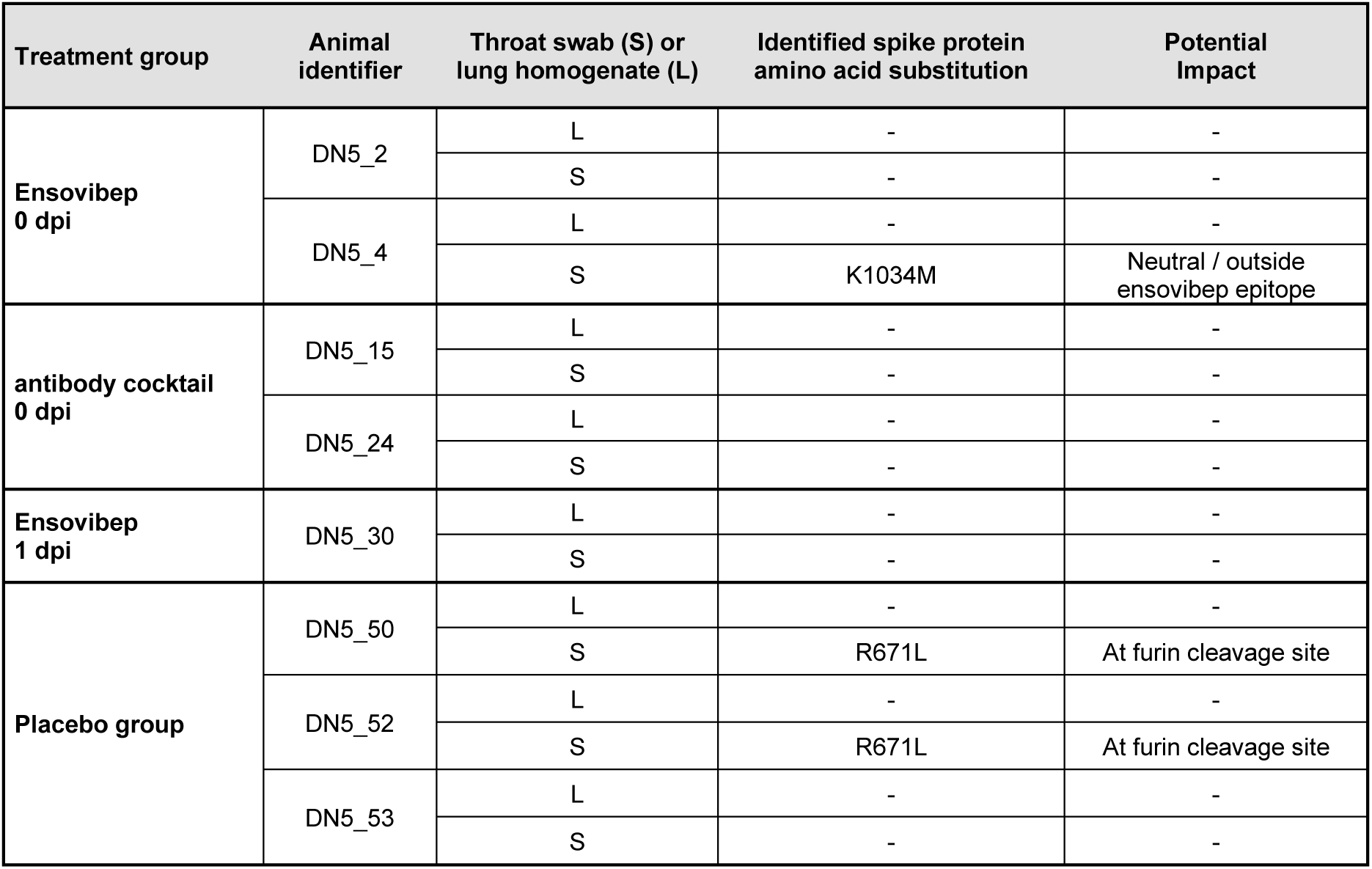
Identification of escape mutations by deep sequencing of SARS-CoV-2 Alpha variant B.1.1.7 in animals at day 5 p.i., which indicated remaining viral titers. As a control, three non-treated animals were also deep sequenced. Deep Sequencing was performed from either swab (S) or lung (L) extracted RNA.

| Treatment group | Animal identifier | Throat swab (S) or lung homogenate (L) | Identified spike protein amino acid substitution | Potential Impact |
| --- | --- | --- | --- | --- |
| Ensovibep 0 dpi | DN5_2 | L | - | - |
|  |  | S | - | - |
|  | DN5_4 | L | - | - |
|  |  | S | K1034M | Neutral / outside ensovibep epitope |
| antibody cocktail 0 dpi | DN5_15 | L | - | - |
|  |  | S | - | - |
|  | DN5_24 | L | - | - |
|  |  | S | - | - |
| Ensovibep 1 dpi | DN5_30 | L | - | - |
|  |  | S | - | - |
| Placebo group | DN5_50 | L | - | - |
|  |  | S | R671L | At furin cleavage site |
|  | DN5_52 | L | - | - |
|  |  | S | R671L | At furin cleavage site |
|  | DN5_53 | L | - | - |
|  |  | S | - | - |

**Supplementary Table 6:**
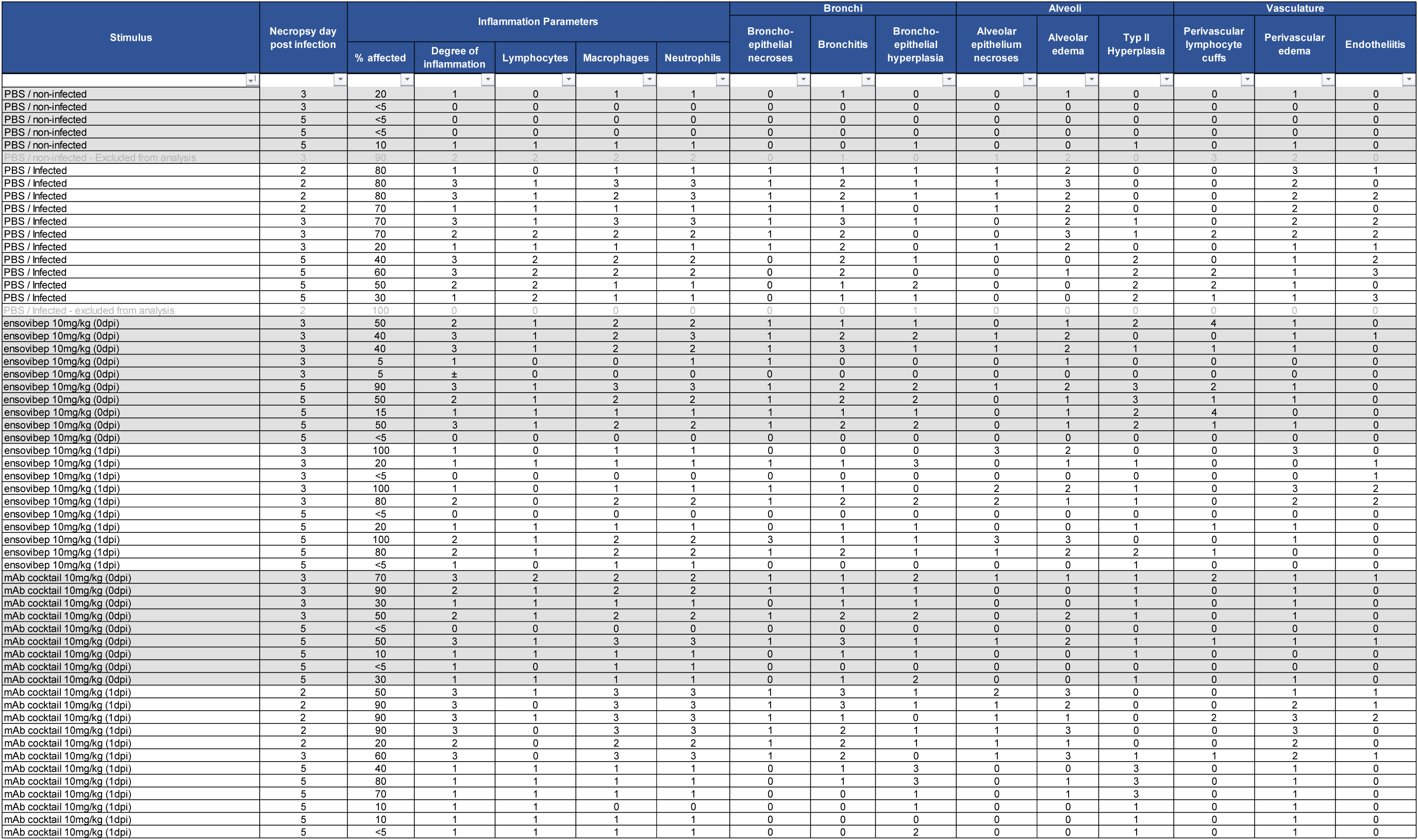
Histopathology data table

